# GM-CSF Programs Alveolar Macrophages to Maintain Pulmonary Lymphocyte Homeostasis

**DOI:** 10.64898/2026.09.28.753165

**Authors:** Paritha Arumugam, Minzhe Guo, Claire Williams, Brenna Carey, Matthew Wessendarp, David Read, Michelle Mantilla, Dana Jackson, Jennifer Stock, Rathnakumar Kumaragurubaran, Takuji Suzuki, Grant Waterer, Cole Trapnell, Bruce C. Trapnell

## Abstract

Pulmonary lymphocytosis is a common feature of chronic lung disease, but the mechanisms governing lymphocyte accumulation within the lung remain poorly understood. Here, we identify granulocyte-macrophage colony-stimulating factor (GM-CSF) as a central regulator of pulmonary lymphocyte homeostasis. Using genetic models, single-nucleus and spatial transcriptomics, cell trafficking analyses, and pulmonary macrophage replacement studies, we show that loss of GM-CSF signaling causes profound pulmonary lymphocytosis characterized by lymphoid follicle formation, local immunoglobulin production, and expansion of multiple lymphocyte lineages. Mechanistically, GM-CSF signaling deficiency blocks terminal alveolar macrophage differentiation and reprograms alveolar macrophages into inflammatory states marked by reduced PU.1 and PPARγ expression, enhanced NF-κB signaling, and expression of lymphocyte-recruiting chemokines. These changes promote lymphocyte recruitment, local proliferation, and lymphoid lineage commitment. Restoration of GM-CSF-responsive alveolar macrophages re-established macrophage identity, suppressed inflammatory chemokine programs, and reversed pulmonary lymphocytosis. Together, these findings identify alveolar macrophages as gatekeepers of adaptive immune homeostasis in the lung.

## INTRODUCTION

Mammalian lungs comprise a vast, ultrathin gas-exchange surface juxtaposed to an equally vast capillary network that increases the surface-to-volume ratio sufficiently to support efficient gas exchange. Exposure to environmental air occurs via a network of branching airways terminating in alveoli that are physically interconnected via complex parenchymal tethering, which balances intraluminal pressure and alveolar/airway recoil in synchronous, tensegrity-like fashion,^1^ thereby keeping distal airways open during the breath cycle and minimizing air-trapping during exhalation.^2^ Immune protection of the lung surface is achieved by (topologically exterior) tissue-resident alveolar macrophages (AMs) and (topologically interior) circulating innate and adaptive immune cells in blood-filled capillaries. Thus, circulating lymphocytes are continuously available on an as-needed basis and can be recruited rapidly throughout the lung surface via peribronchovascular high endothelial venules.^3^ Notwithstanding, pulmonary lymphocyte recruitment is tightly regulated^4^ and bronchus-associated lymphoid tissues (BALT) is minimal in healthy lungs and, importantly, increased pulmonary lymphocyte recruitment and the resulting hyperplasia of peribronchiolar BALT can distort and narrow small airways, disrupt airflow, and impair lung function.^5^

Pulmonary lymphocytosis—the abnormal accumulation of lymphocytes in lung tissue—occurs in multiple chronic lung diseases and is characterized by distortion of pulmonary architecture by peribronchovascular, alveolar, and interstitial lymphocyte accumulation, follicle formation, constrictive bronchiolitis, fibrosis, and cyst development.^6^ Pulmonary lymphocytosis occurs in individuals with complete deficiency of ‘C-C’ motif chemokine receptor 2 (CCR2) in association with a reduction in alveolar macrophages and results in constrictive cellular bronchiolitis and polycystic lung disease caused by increased intraluminal pressure from a check-valve mechanism.^7^ Pulmonary lymphocytosis occurs in numerous other chronic lung diseases including hypersensitivity pneumonitis,^8,9^ asthma,^10,11^ chronic obstructive disease,^12–14^ fibrotic diseases,^6^ autoimmune lung diseases,^15^ and pulmonary alveolar proteinosis (PAP)-causing diseases.^7,16,17^ Since their discovery in the late 1970s^18^, our understanding of chemokines, their cognate receptors, and the pathways by which they recruit lymphocytes have been extensively characterized(reviewed in ^19^). However, the mechanisms that initiate and sustain pulmonary lymphocytosis in lung diseases remain largely unknown.

We studied pulmonary lymphocytosis caused by granulocyte/macrophage-colony stimulating factor (GM-CSF) signaling deficiency was because it arises from disruption of signaling by a single molecule, GM-CSF, providing a uniquely defined etiologic driver and because GM-CSF signaling deficiency causes an identical lung disease, PAP, in humans (e.g., autoimmune PAP, hereditary PAP) and mice (e.g., *Csf2*^KO^, *Csf2ra*^KO^, and *Csf2rb*^KO^) ^20–24^ GM-CSF a central regulator of myeloid cell differentiation,^25^ GM-CSF is required for the development and maintenance of AMs,^26–28^ and establishes AM identity via transcriptional programs including PU.1, C/EBPβ, and PPARγ.^26–29^ Without GM-CSF stimulation, AMs have impaired innate and homeostatic functions including clearance of surfactant, cell debris, and inhaled particulate matter.^23,24,26,30,31^ Beyond its role in macrophage biology, GM-CSF shapes innate-adaptive immune interactions through effects on cytokine production, antigen presentation, and lymphocyte activation,^32–35^ promotion of leukocyte recruitment, and by amplifying adaptive immune responses in inflammatory and autoimmune diseases.^36^ Notwithstanding, how GM-CSF signaling contributes to lymphocyte homeostasis in health and its deficiency contributes to pulmonary lymphocytosis in disease remains unclear. Here, using integrated single-cell, spatial, and functional analyses, we identify inflammatory AM populations that emerge in response to impaired GM-CSF signaling and are associated with pulmonary lymphocytosis. Restoration of GM-CSF signaling in AMs suppresses these programs and reduces lymphocyte accumulation in the lung. Together, our findings identify GM-CSF-dependent AMs as gatekeepers of pulmonary lymphocyte homeostasis and reveal how disruption of tissue macrophage identity remodels the adaptive immune architecture of the lung.

## RESULTS

### GM-CSF signaling deficiency Causes Severe Pulmonary Lymphocytosis

Granulocyte–macrophage colony-stimulating factor (GM-CSF) signaling deficient (GMSD) mice spontaneously developed increased accumulation of lymphocytes compared to wild type (WT) mice, particularly in peribronchovascular regions, irrespective of whether the disruption was caused by absence of the ligand or disruption of the GM-CSF-specific receptor α chain or the β chain common to receptors for GM-CSF, IL-3, and IL-5 (**Figures 1A-C** and **S1A-F**). The abnormal cell foci also contained macrophages (**Figures 1D, S1C, and S1F**) and formed lymphoid follicles (**Figure 1E-J**) containing B220+, CD3+, and CD68+ cells corresponding to B cells, T cells, and macrophages, respectively (**Figure 1F-H** and **S1G-H**). Cells in lymphoid aggregates also stained positive for Pax5 and Ki67 consistent with in-situ replication of lymphocytes (**Figures 1I-J** and **S1I**). Transmission electron microscopy confirmed that peribronchovascular cell aggregates contained primarily lymphocytes but also included plasma cells (**Figure 1K-M**). The increase in plasma cells in the lungs of GMSD mice was confirmed by an increase in CD138+ cells among lung digest cells (**Figure 1N**). The increase in abnormal lymphocyte foci was similar in *Csf2^KO^*, *Csf2ra^KO^*, and *Csf2rb^KO^* mice (**Figure 1O**) identifying pulmonary lymphocytosis as a consistent feature of GM-CSF signaling disruption. Pulmonary histopathology also included patchy accumulation of surfactant (**Figure 1B and S1J**), cholesterol crystals (**Figure S1K**), and foamy-appearing alveolar macrophages and non-cleared debris (**Figure S1L**), which was not necessarily coincident with pulmonary lymphocyte accumulation in GMSD mice. Spatial transcriptomic analysis of peribronchovascular regions in GMSD and WT lung tissues (**Figure S2**) revealed that lymphocyte accumulations in GMSD mice showed increased expressed genes involved in T and B cell receptor signaling, activation, differentiation, and proliferation and expression of multiple immunoglobin class genes (IgM, IgA, IgG1, IgG2b, IgG3 and J-chain) (**Figures 1P** and **S2D-F**). In lymphoid cell accumulations, downregulated genes included regulation of endothelial cell migration, cell junction assembly and organization.

**Figure 1.**
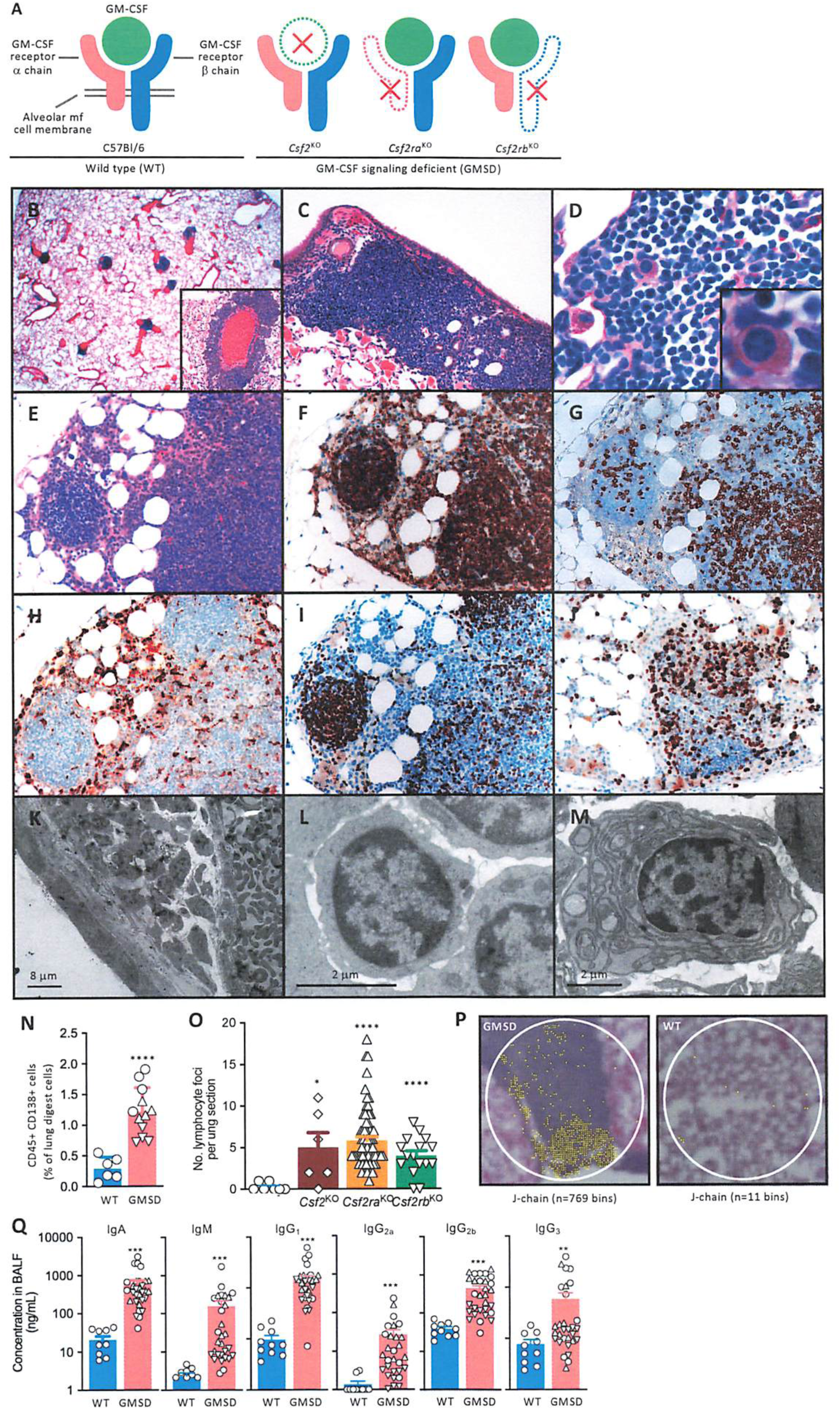
GM-CSF signaling deficiency drives pulmonary lymphocytosis. Loss of GM-CSF signaling results in prominent peribronchovascular lymphoid cell accumulations and immunoglobulin production in the lung. (A) Schematic of the GM-CSF ligand, GM-CSF receptor (*Csf2ra*/*Csf2rb*) complex and GM-CSF signaling-deficient mouse models (*Csf2*-/-, *Csf2ra*-/- and *Csf2rb*-/-used in this study. Red Xs denote deletion of the corresponding ligand or receptor genes. (B-D) Representative H&E-stained lung sections from *Csf2ra*−/− mice showing progressive peribronchovascular cellular accumulations. Magnification: B-1x, inset 10X; C-10X, D-63X (E-J) Characterization of lymphoid aggregates by H &E (E), immunostaining for B220 (F), CD3 (G), CD68 (H), PAX5 (I), and Ki67 (J), Magnification-20x. (K-M) Transmission electron micrographs showing lymphocyte and plasma cell accumulation in peribronchovascular regions (Magnification-K-2480x, L, M-7480x,). (N) Quantification of CD138+ plasma cells among lung CD45+ cells by flow cytometry. (O) Histopathological scoring of pulmonary lymphocytosis in WT and GM-CSF signaling-deficient (GMSD) mice. Data are the sum of abnormal peribronchovascular lymphocyte foci obtained by counting low-power (1x) micrographs for 32-38 lung tissue sections from 5 mice per condition. Each symbol represents one region of lymphocyte accumulation in lung sections. (P) Spatial transcriptomic analysis of *Jchain* expression within peribronchovascular lymphoid aggregates. (Q) Immunoglobulin concentrations in bronchoalveolar lavage (BAL) fluid, including IgA, IgM, IgG1, IgG2a, IgG2b, and IgG3. Each symbol represents data from one mouse. GMSD mice *Csf2*-/-(white diamonds), *Csf2ra*-/-(white upward triangles) and *Csf2rb*-/-(white downward triangles). Abbreviations: GMSD=GM-CSF signaling deficient (i.e., *Csf2*-/-, *Csf2ra*-/- and *Csf2rb*-/-), WT=wild type. N=6-10 mice/genotype). Data shown are the means ± SEM. Significance was assessed using Mann-Whitney U tests; *P<0.05, **P<0.01, ***P<0.001 and ****P<0.0001.

The increased expression of immunoglobulin genes in pulmonary lymphocytes was confirmed by the observation of marked increases in IgM, IgA, IgG1, IgG2a, IgG2b, and IgG3 in the BAL fluid from GMSD compared to WT mice (**Figure 1Q**).

### Pulmonary alveolar macrophages are present in the absence of GM-CSF signaling

To further define the pulmonary consequences of GM-CSF signaling deficiency, we performed single-nucleus RNA sequencing (snRNAseq) analysis on 33,576 lung cells from 8 WT and 16 GMSD mice (**Figure S3A**). Cells corresponding to the five major lung cell lineages—epithelial, endothelial, mesenchymal, myeloid, and lymphoid—were similar in number in WT and GMSD mice except for lymphoid cells, which were increased in GMSD mice (**Figures 2A** and **S3B-D**). Compared to WT, all three GMSD mouse lines exhibited a similar pattern of differential gene expression for each of the corresponding major lung cell lineages (**Figures S3E**). Differentially expressed genes were analyzed using the Kyoto Encyclopedia of Genes and Genomes (KEGG) database to compare the five major lung cell lineages in GMSD and WT mice (**Figure S4, Supplemental table S3**). Lymphocytes had increased expressed of genes associated with lymphocyte activation, antigen receptor signaling, and proliferation (**Figure S4A**). Epithelial cells had increased expression genes associated with tissue injury, remodeling, stress, metabolic adaptation, mucosal defense, barrier function, and cell turnover consistent with dysregulated epithelial cell homeostasis and environmental stress (**Figure S4B**). Endothelial cells expressed genes related to vascular remodeling and leukocyte trafficking (**Figure S4C**). Mesenchymal cells had increased expression of genes encoding extracellular matrix proteins, matrix remodeling enzymes, cytoskeletal regulators, and contractile proteins consistent with myofibroblast-like differentiation, activation of a tissue remodeling program, and fibrosis (**Figure S4D**). Myeloid cells had increased expression of genes associated with inflammatory and immune signaling (**Figure S4E**).

**Figure 2.**
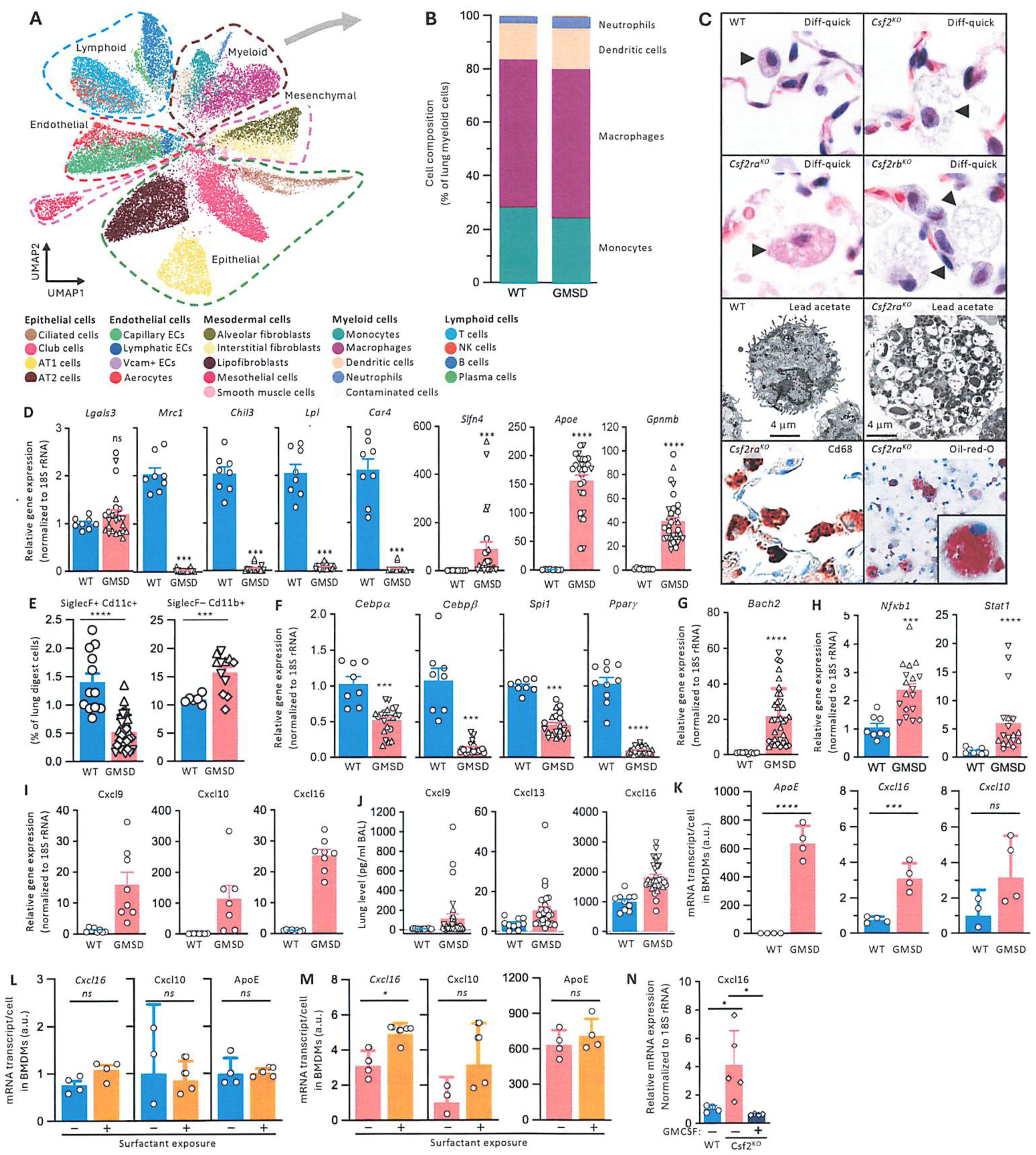
Pulmonary lymphocytosis occurs despite the presence of alveolar macrophages. GM-CSF signaling-deficient lungs harbor phenotypically remodeled alveolar macrophages with features of altered differentiation and activation. (A) Integrated UMAP of all lung cells identified by snRNA-seq. (B) Relative abundance of major epithelial, endothelial, mesenchymal, myeloid, and lymphoid cell lineages in WT and GMSD lungs. (C) Morphologic characterization of BAL cells demonstrates the persistence of alveolar macrophages in all GMSD strains, including ultrastructural, CD68, and Oil Red O analyses. (D) Expression of mature alveolar macrophage genes and inflammatory macrophage genes. (E) Quantification of Siglec-F+CD11c+ alveolar macrophages and Siglec-F−CD11b+ myeloid cells in lung digests. (F) Gene expression of lineage defining transcription factors *Cebpα, Cebpβ, Spi1 and Pparγ*, (G) *Bach2*(H) *Pparγ*-associated signaling genes *NF-κb* and *Stat1*. (I and J) Expression and BAL concentrations of chemokines in WT and GMSD lungs. (K) Inflammatory and chemokine gene expression in WT and GMSD alveolar macrophages. (L and M) Chemokine expression in WT and GMSD macrophages at baseline and after exposure to PAP-derived surfactant (BMDMs) N=4 samples/group. (N) Chemokine expression in *Csf2*-/-BMDMs after GM-CSF addition. Each symbol represents data from one mouse. GMSD mice *Csf2*-/-(white diamonds), *Csf2ra*-/-(white upward triangles) and *Csf2rb*-/-(white downward triangles). Abbreviations: GMSD=GM-CSF signaling deficient (i.e., *Csf2*-/-, *Csf2ra*-/- and *Csf2rb*-/-), WT=wild type. N=6-10 mice/genotype). Data shown are the means ± SEM. Significance was assessed using Mann-Whitney U tests; NS-not significant, *P<0.05, **P<0.01, ***P<0.001 and ****P<0.0001.

Because the pattern of differential gene expression between GMSD and WT mice was most prominent in myeloid cells compared other lineages (**Figure S3E**), a detailed analysis was undertaken. All the main myeloid lineages—neutrophils, dendritic cells, macrophages, and monocytes—were present and in roughly similar proportions in GMSD and WT mice (**Figures 2B**) and the cell types identified included monocytes, neutrophils, four dendritic cell subpopulations, interstitial macrophages, and five alveolar macrophage subpopulations (**Figure S5A-C**). Microscopic examination of cells recovered by bronchoalveolar lavage (BAL) confirmed that alveolar macrophages were present in GMSD mice but, compared to WT, were enlarged, foamy-appearing, and contained numerous intracytoplasmic inclusion vesicles (**Figure 2C**). GMSD alveolar macrophages also stained strongly with oil-red-O (**Figures 2C and S5D**), consistent with accumulation of esterified cholesterol-rich intracellular lipid droplets as reported for GMSD alveolar macrophages in humans^37^ and mice^38^.

Compared to WT, GMSD alveolar macrophages had normal expression of a canonical macrophage gene (*Lgals3)*, reduced expression of genes normally expressed in healthy mature alveolar macrophages (*Mrc1, Chil3, Lpl,* and *Car 4*), and abnormally increased expression of genes related to lipid metabolism and inflammation (*Slfn4, Apoe, and Gpnmb*) (**Figure 2D**). Gpnmb was also increased on GMSD CD45+ lung digest cells and in soluble form in GMSD BAL fluid compared to WT (**Figures S5E-F**). CD68, a phenotypic marker of macrophages, was readily detected on GMSD alveolar macrophages recovered by BAL (**Figure 2C**); however, Siglec-F, a phenotypic marker of healthy mature alveolar macrophages was reduced on GMSD CD11c+ lung digest cells and CD11b, a phenotypic marker of myeloid cells not normally present on alveolar macrophages^39,40^ was increased on GMSD Siglec-F–lung digest cells (**Figures 2E** and **S5G**). GMSD alveolar macrophages recovered by BAL also had reduced expression of transcription factors important in alveolar macrophage differentiation (*Cebpα*, *Cebpβ*, *Spi1*, and *Pparγ*) (**Figure 2F**) and increased expression of a transcription factor normally expressed in lymphocytes (*Bach2*) (**Figure 2G**). Compared to WT, BAL from GMSD mice was milky-appearing and had increased turbidity, surfactant, and cell debris (**Figures S5H-J**, **S1L** and **2C**), consistent with prior observations of impaired clearance of surfactant, particles, and cell debris by GMSD alveolar macrophages.^23,24^

### Absence of GM-CSF stimulation reprograms alveolar macrophages to a pro-inflammatory state

Since Pparγ is reported to suppress Nfκb signaling-stimulated chemokine expression in macrophages^41–44^ and *Pparγ* expression was markedly reduced in GMSD alveolar macrophages,^45^ we evaluated the expression of genes related to Nfκb signaling, chemokine secretion, and inflammatory mediators.

Compared to WT, GMSD alveolar macrophages recovered by BAL had increased expression of *Nffib1* and *Stat1* (**Figure 2H**), increased expression of multiple chemokines including *Cxcl9*, *Cxcl10*, *Cxcl16* (**Figure 2I**) and BAL from GMSD mice had increased levels of multiple chemokines including Cxcl9, Cxcl13, Cxcl16, Ccl2, Ccl7, Ccl8 (**Figures 2J** and **S5K**) and inflammatory mediators (**Figures S5F** and **S5L**). To determine if the increased chemokine secretion by GMSD alveolar macrophages was a primary abnormality caused by GM-CSF signaling deficiency or a secondary response from exposure to uncleared surfactant and debris, primary bone marrow-derived macrophages (BMDMs) from WT and GMSD mice were evaluated *ex vivo*. After culturing in media containing M-CSF ex vivo for 5 days, GMSD BMDMs spontaneously expressed *Apoe*, *Cxcl16*, and *Cxcl10* at increased levels compared to WT BMDMs (**Figure 2K**). Exposure to surfactant-rich lung lavage sediment from PAP patients resulted in no change in the expression of these genes in WT BMDMs (**Figure 2L**) and minimal to no change in GMSD BMDMs (**Figure 2M**). Importantly, the increased *Cxcl16* expression in GMSD BMDMs was reduced to normal by the addition of GM-CSF to the culture media (**Figure 2N**).

To explore the mechanism(s) by which alveolar macrophages are reprogramed, we evaluated the expression of cytokines reported to regulate alveolar macrophages differentiation.^27–29,46–49^ While levels of GM-CSF detectable in BAL varied by genotype as expected, compared to WT, BAL from all three GMSD mouse lines had increased levels of M-CSF and IL-34 (**Figure S6A-C**) and increased TGFβ only in older mice (**Figure S6D**). While GM-CSF receptor-mediated signaling and cellular proliferation was detectable only in WT BMDMs (**Figure S6E-G**), expression of M-CSF receptors was detected in all (**Figure S6H**) and M-CSF receptor-mediated signaling and cellular proliferation was detectable and similar in WT and GMSD BMDMs (**Figure S6I-J**). Compared to WT, GMSD mice had increased M-CSF in BAL fluid and GMSD alveolar macrophages had increased M-CSF mRNA and M-CSF secretion and all three findings were reversed in *Csf2*^KO^ mice by transgenic expression of GM-CSF exclusively in the lungs (**Figure S6K-N**). In primary alveolar macrophages, M-CSF expression varied inversely with expression of *Spi1* encoding the master macrophage transcription factor PU.1.^26^ In cultured alveolar macrophage cell lines, M-CSF secretion, mRNA levels, and transcription were increased in the *Csf2*^KO^ compared to WT and were reduced by transduction-mediated *Spi1* expression and M-CSF mRNA stability was unaffected by *Csf2* expression (**Figure S6O-R**). Interestingly, the chemokines Ccl2, Ccl7, and Ccl8 followed a similar pattern with respect to levels of the protein in BAL and transcription in cultured alveolar macrophage cell lines (**Figure S6S-V**).

To further define alveolar macrophage cell states in GMSD and WT mice, we analyzed the lung snRNAseq data and identified 5 distinct alveolar macrophage subpopulations (AM1–5) (**Figure 3A-B**). Although the total numbers of macrophages were similar between GMSD and WT mice (**Figure 2B**), the relative proportions of corresponding alveolar macrophage subpopulations in each differed markedly (**Figure 3A-B**). In WT lungs, alveolar macrophages were predominantly composed of the AM1 population, with a smaller contribution from AM2 cells and minimal representation of AM3-AM5 states. In contrast, GMSD alveolar macrophages were shifted towards expansion of the AM2 population, and accumulation of AM3-AM5 inflammatory macrophage states and significant reduction of AM1 cells. (**Figure 2A-B**). Alveolar macrophage subpopulations exhibited distinct differential gene expression profiles (**Figure S7A**) and KEGG pathway activation (**Figure S7B, Supplemental Table 3**). AM1 cells exhibited a differential gene expression profile partially overlapping with AM2 cells (**Figure S7A**) and included the expression of canonical markers of mature alveolar macrophages including *Chil3*, *Plet* 1, *Lpl*, *Car4*, and *Ccl6* (**Figures 3C**). AM3 cells had an expression profile differing extensively from other subpopulations (**Figures S7A**) and included increased expression of genes involved in lipid metabolism, cholesterol clearance, and efferocytosis (**Figures 3C**), suggesting a cell state with heightened macrophage activity and tissue remodeling. AM4 and AM5 exhibited a partial overlapping differential gene expression profile that was distinct from that of AM1, AM2, and AM3 cells (**Figures S7A**) and included multiple interferon-induced genes, proinflammatory genes, and chemokines capable of recruiting lymphocytes (**Figures 3C**). Spatial transcriptomics analysis revealed the increased expression of T-, NK-, and B-cell chemokines (Cxcl9, Cxcl10, Cxcl13, Cxcl16, others) in the lungs of GMSD mice occurred in regions of peribronchovascular lymphocyte accumulation in a pattern consistent with lung-vascular gradients (**Figure 3D**).

**Figure 3.**
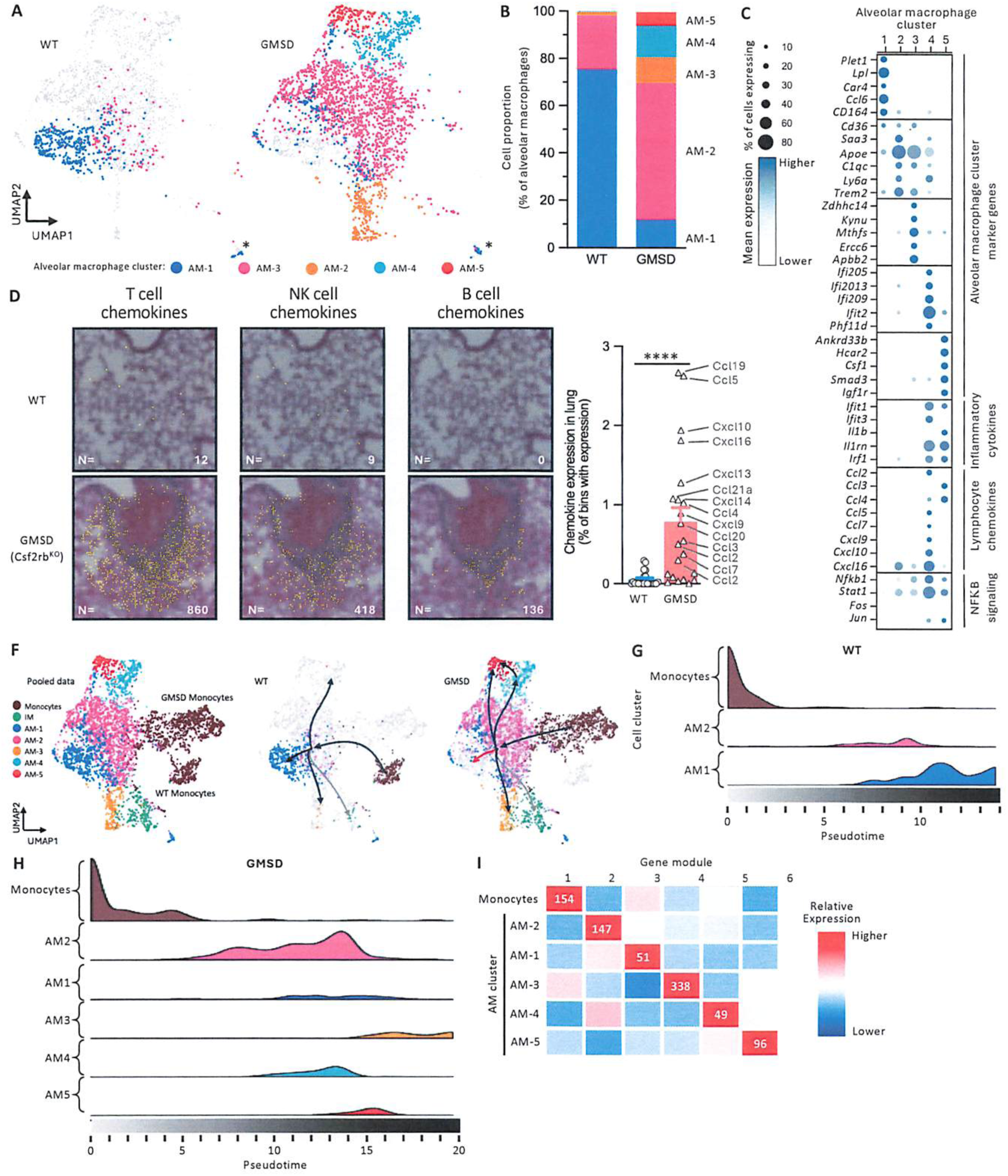
GM-CSF deficiency reprograms alveolar macrophages toward a chemokine-producing pro-inflammatory state. SnRNA-seq analyses identify distinct inflammatory alveolar macrophage states in GMSD lungs. (A) UMAP visualization of alveolar macrophage populations from WT and GMSD mice. (B) Quantification of alveolar macrophage cluster composition across genotypes. (C) Cluster-defining gene expression signatures, including alveolar macrophage genes, inflammatory cytokines, lymphocyte-attracting chemokines, and NF-κB pathway components. (D-E) Spatial identification of peribronchovascular T, B and NK-cell chemokine gradients in GMSD lungs. (F) Pseudo time analysis demonstrates divergence of WT and GMSD alveolar macrophage states. (G-H) Distribution of alveolar macrophage clusters across pseudotime trajectories. (I) Heatmap showing temporally regulated gene expression modules associated with macrophage reprogramming. Abbreviations: GMSD=GM-CSF signaling deficient (i.e., *Csf2*-/-, *Csf2ra*-/- and *Csf2rb*-/-), WT=wild type. N=4-8 mice/genotype), AM-alveolar macrophages. Data shown are the means ± SEM. Significance was assessed using Mann-Whitney U tests; ****P<0.0001.

Because the differential accumulation of alveolar macrophage subpopulations in GMSD mice compared to WT—markedly reduced AM1 cells and increased AM2 cells—suggested a block in terminal differentiation, pseudotime analysis was performed. Monocytes were used as the pseudotime origin since they are capable of differentiating into macrophages,^50^ were present in WT and GMSD mice (**Figure S5B**), and expressed canonical monocyte markers similarly in both (**Figure S8A**) albeit with a small split in the UMAP positions indicating differential effects of GM-CSF on monocytes in the lungs (**Figure 3F**). In WT mice, monocytes differentiated along pseudotime into AM2 cells and finally to AM1 cells (**Figures 3G**). In GMSD mice, monocytes differentiated along pseudotime into AM2 where they appeared to accumulate; only a few progressed to AM1 with larger numbers progressing into AM3, AM4, and AM5 (**Figure 3H**). Alveolar macrophage subpopulations expressed distinct gene expression ‘modules’ (**Figures 3I** and **S8C-D**), and cell functions as determined by KEGG analysis (**Figure S7B, Supplemental Table 3**). AM4 and AM5 more strongly expressed macrophage M1 phenotype genes while AM1, AM3, monocytes, and interstitial macrophages expressed M2 phenotype genes (**Figure S8E-F**).

### GM-CSF signaling deficiency results in increased pulmonary lymphocyte recruitment, in-situ proliferation, and lymphoid lineage commitment

To determine the mechanism(s) driving pulmonary lymphocyte accumulation in GMSD mice, we analyzed pulmonary lymphocyte gene expression, trafficking and proliferation markers. Compared to WT, GMSD mice had generalized increase in pulmonary B-, T-, NK-, and plasma cells (**Figure 4A-C**) with preservation of the relative proportions of subpopulations (**Figure 4D**). The expression of genes associated with recruitment of lymphocytes into tissues was increased in endothelial cells and pulmonary lymphocytes in GMSD compared to WT (**Figure 4E**). Lymphocyte trafficking was evaluated by passive transfer of congenic green fluorescent protein (GFP)-expressing splenocytes, which demonstrated increased numbers of GFP+ splenocytes in the lungs of GMSD mice compared to WT, particularly in peribronchovascular regions associated with lymphocyte accumulation (**Figure 4F-G**).

**Figure 4.**
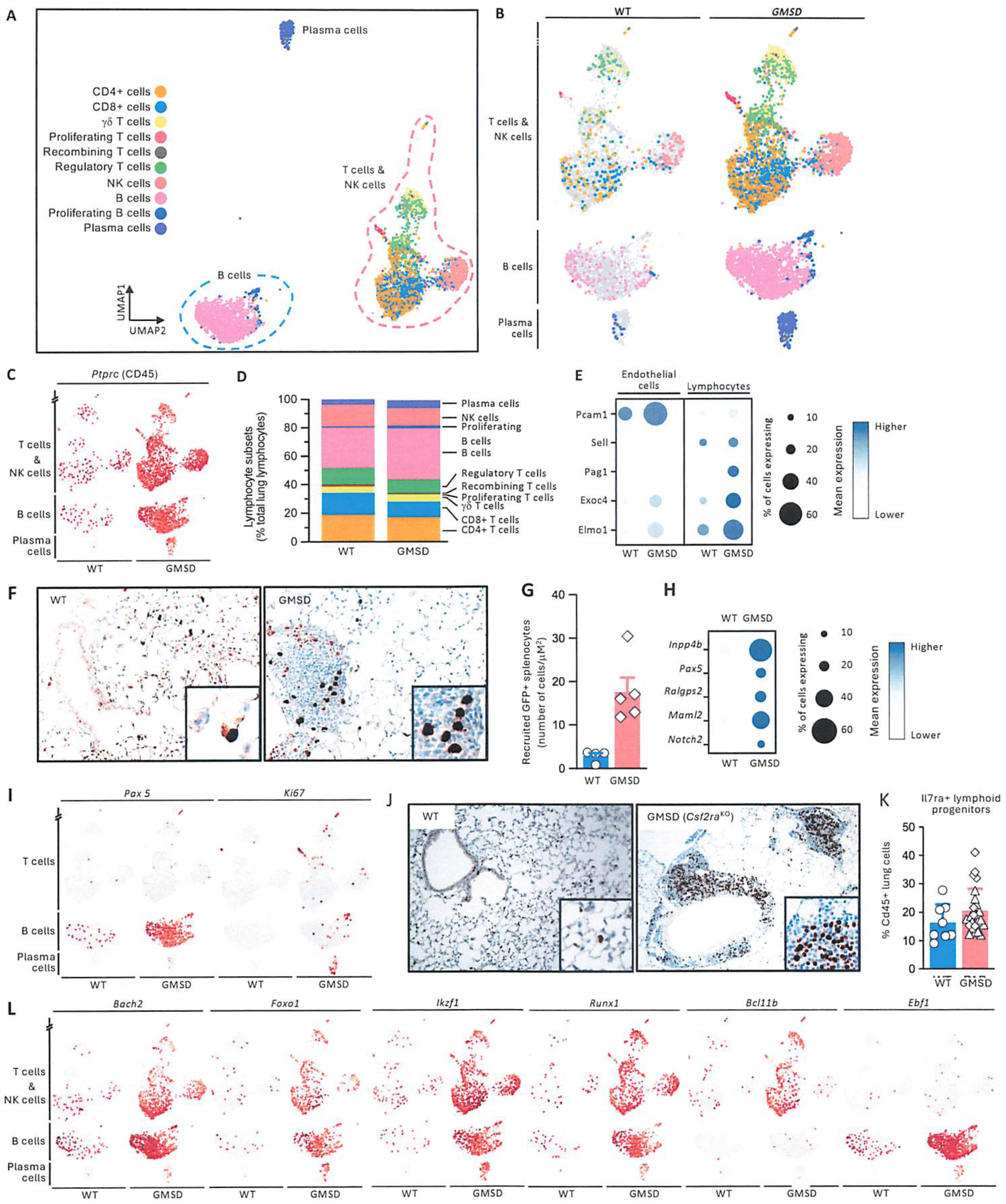
Inflammatory macrophage reprogramming promotes chemokine-dependent lymphocyte recruitment and pulmonary lymphocytosis. Loss of GM-CSF signaling promotes recruitment and expansion of lymphocyte populations within the lung. (A and B) UMAPs showing integrated and genotype-specific analyses of pulmonary lymphoid populations. (C) Feature plots illustrating lineage-specific lymphocyte (CD45) marker expression. (D) Quantification of lymphocyte subset abundance in WT and GMSD lungs. (E) Expression of genes associated with lymphocyte trafficking (leukocyte adhesion and migration cascade) in endothelial and lymphoid populations. (F and G) Adoptive transfer experiments demonstrating enhanced recruitment of GFP-labeled lymphocytes into *Csf2ra*−/− lungs, Magnification-20x (panel F), Inset 40X. (H and I) Analyses of proliferation markers (Inpp4b, Ralgps2) and lymphoid transcriptional regulator (Pax5, Notch2, Maml2) gene expression in pulmonary lymphocyte populations. (J) Representative Ki67 immunostaining demonstrating increased lymphocyte proliferation in WT and GMSD lungs, Magnification-5x., inset-zoomed-in view (K) IL-7rα expression in CD45+ lung cells. (L) Expression of lineage commitment genes across pulmonary lymphocyte subsets. Each symbol represents data from one mouse. Abbreviations: GMSD=GM-CSF signaling deficient (i.e., *Csf2*-/-(white diamonds), *Csf2ra*-/-(white upward triangles) and *Csf2rb*-/-(white downward triangles)), WT=wild type. N=4-8 mice/genotype). Data shown are the means ± SEM. Significance was assessed using Mann-Whitney U tests; NS-not significant, *P<0.05.

In situ lymphocyte proliferation was evaluated by gene expression analysis and immunohistochemical staining, which demonstrated increased expression of replication-associated genes (**Figure 4H-I**) and increased Ki67 immunostaining in the lungs of GMSD mice compared to WT (**Figure 4J**). Compared to WT, the lungs of GMSD mice had increased numbers of lymphoid precursors (**Figure 4K**), and increased expression of transcription factors involved in lymphoid lineage commitment – *Bach2, Foxo1, IKzf1, Runx1, Bcl11b*, and *Ebf1* (**Figure 4L**).

### Pulmonary macrophage transplantation to restore GM-CSF signaling in alveolar macrophages restores pulmonary lymphocyte homeostasis

Given the marked increased expression of multiple chemokines by alveolar macrophages in GMSD mice, we tested if restoring GM-CSF signaling exclusively to the alveolar macrophage population would normalize pulmonary lymphocyte homeostasis. To accomplish this, *Csf2ra*^KO^ recipient mice received congenic *Csf2ra* gene-corrected macrophages by direct pulmonary instillation—pulmonary macrophage transplantation or PMT hereafter.^51^ Compared to untreated mice, alveolar macrophages recovered by BAL from recipient mice 2-months after PMT had increased expression of *Cebpβ, Chil3, Plet 1, Mrc1, and Lpl*, and reduced expression of *Bach2*, *Gpnmb, ApoE, Cxcl9, Cxcl10, and Cxcl16* (**Figure 5A-D**), consistent with restoration of a GM-CSF responsiveness to the alveolar macrophage population. BAL from recipient mice also had reduced levels of Cxcl16 compared to untreated mice (**Figure 5E**). One year after PMT, alveolar macrophages from recipient mice had markedly increased expression of *Pparγ*, reduced expression of *Nfκb* signaling-related genes (*Nffib1*, *Stat1*, *Fos*), increased levels of the mature alveolar macrophage marker Siglec-F (**Figure 5F-H**). One year after PMT, BAL fluid from recipient mice had markedly reduced turbidity indicating a reduction in abnormal surfactant accumulation (**Figure 5I**). Compared to untreated mice, PMT-treated mice had fewer histologic foci of pulmonary lymphocyte accumulation after 6-, and 12-months (**Figure 5J-K**), normal numbers of B220+ and CD3+ in lung digests (Figure 5L), and reduced levels of immunoglobulins in BAL fluid (**Figure 5M**).

**Figure 5.**
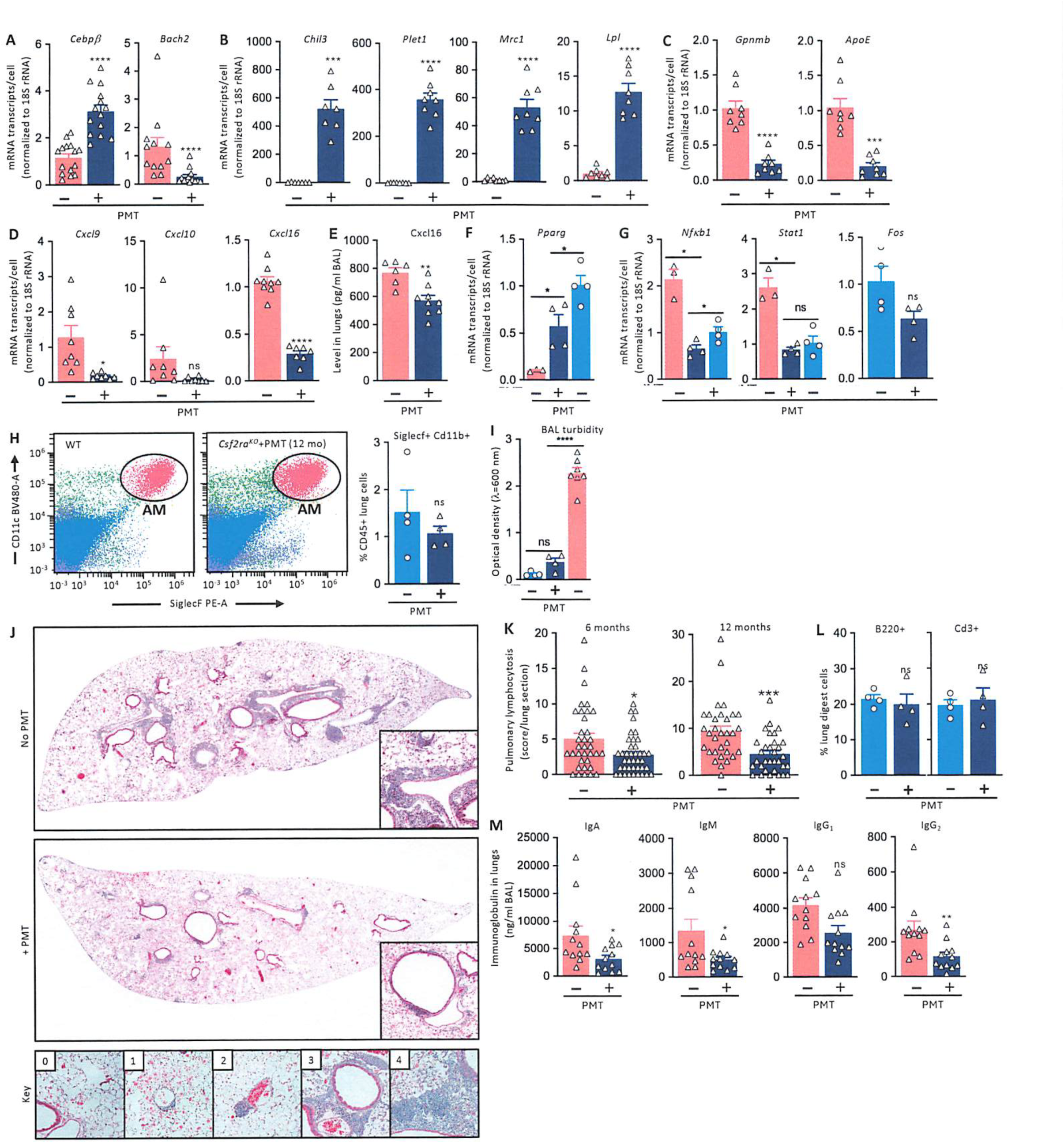
Restoration of GM-CSF signaling in alveolar macrophage re-establishes pulmonary immune homeostasis. Pulmonary macrophage transplantation re-establishes alveolar macrophage identity and suppresses pulmonary lymphocytosis in GMSD mice. (A-E) Expression of lineage-defining transcription factors, mature alveolar macrophage genes, inflammatory genes, chemokine genes, and BAL chemokine concentrations 2 months after pulmonary macrophage transplantation (PMT). (F and G) Expression of *Pparγ* and NF-κB signaling genes 12 months after PMT. (H) Representative flow plot and quantification of Siglec-F+CD11c+ alveolar macrophages following PMT. (I) BAL turbidity in untreated WT, untreated GMSD, and PMT-treated GMSD mice. (J) Representative lung sections of *Csf2ra*^-/-^showing reduced peribronchovascular lymphoid accumulations after PMT, Magnification-1x, inset-zoomed-in view. (K) Quantification of pulmonary lymphocytosis at 6 and 12 months following PMT. Data are the sum of abnormal peribronchovascular lymphocyte foci obtained by counting low-power (1x) micrographs for 32-38 lung tissue sections from 5 mice per condition. Each symbol represents one region of lymphocyte accumulation in lung sections. Key for scoring is detailed in methods section. (L) Quantification of pulmonary B220+ B cells and CD3+ T cells among CD45+ lung leukocytes in untreated WT and PMT-treated GMSD mice. (M) BAL immunoglobulin concentrations before and after restoration of GM-CSF signaling by PMT. Red bars-*Csf2ra*−/− mice, Dark blue bars-*Csf2ra*−/− mice that received PMT, Teal blue bars-*Csf2ra*−/− mice that received PMT, analysis at 1 year. Each symbol represents data from one mouse. N=4-8 mice/group). Abbreviations: PMT-Pulmonary macrophage transplantation. Data shown are the means ± SEM. Significance was assessed using Mann-Whitney U tests; NS-not significant, *P<0.05, **P<0.01, ***P<0.001 and ****P<0.0001.

## DISCUSSION

We demonstrate that GM-CSF signaling deficiency causes pulmonary lymphocytosis by reprograming AMs to an inflammatory, chemokine-secreting state and identify GM-CSF as a central regulator of pulmonary lymphocyte homeostasis. Disruption of GM-CSF signaling, whether through loss of ligand or receptor function, drove the formation of peribronchovascular lymphoid aggregates enriched in proliferating B cells, T cells, plasma cells, and macrophages, accompanied by robust local immunoglobulin production and widespread activation of adaptive immune programs. AMs persisted in the absence of GM-CSF signaling but exhibited profound loss of mature identity, impaired differentiation, and reprogramming into inflammatory states characterized by activation of NF-κB-associated pathways and expression of multiple lymphocyte-recruiting chemokines. Spatial and trajectory analyses linked the emergence of these inflammatory macrophage states to chemokine gradients within regions of lymphocyte accumulation and revealed a failure of terminal alveolar macrophage maturation that diverted cells toward inflammatory fates. Functionally, pulmonary lymphocytosis resulted from coordinated increases in lymphocyte recruitment, in situ proliferation, and lymphoid lineage commitment. Importantly, selective restoration of GM-CSF responsiveness in alveolar macrophages by pulmonary macrophage transplantation reversed macrophage reprogramming, suppressed chemokine production, and normalized pulmonary lymphocyte accumulation, identifying GM-CSF-dependent alveolar macrophages as central regulators of pulmonary lymphocyte homeostasis and revealing a macrophage-driven mechanism linking disruption of tissue-resident myeloid cell identity to remodeling of the adaptive immune architecture of the lung.

Our observations reframe AMs from cells whose primary function is maintenance of surfactant homeostasis and innate host defense^20,23,52^ to one as master regulators of pulmonary immune architecture. The prevailing paradigm has viewed the major consequence of impaired GM-CSF signaling as loss of AM-mediated clearance functions, resulting in surfactant accumulation and pulmonary alveolar proteinosis.^21,23,24,29,53–56^ Our findings instead reveal that GM-CSF-dependent AMs actively enforce lymphocyte homeostasis throughout the lung by maintaining a differentiated transcriptional state that suppresses inflammatory and lymphocyte-recruiting programs. When this identity program is disrupted, AMs do not simply become dysfunctional scavengers; they adopt alternative inflammatory fates that produce chemokine networks capable of orchestrating lymphocyte recruitment, proliferation, and lymphoid tissue formation. Thus, AMs emerge as gatekeepers that continuously restrain adaptive immune cell accumulation within the pulmonary interstitium and peribronchovascular niche. This conceptual shift positions alveolar macrophages as central organizers of tissue immunity whose differentiated state determines not only surfactant balance, but also the distribution, abundance, and spatial organization of adaptive immune cells within the lung. More broadly, these findings suggest that AM identity itself represents a critical mechanism of lung homeostasis and that disruption of resident macrophage programs may be a unifying driver of tertiary lymphoid structure formation and chronic inflammation across diverse pulmonary diseases.

Our results establish a genetically defined model of pulmonary lymphocytosis in which disruption of a single macrophage-regulating pathway, GM-CSF signaling, is sufficient to remodel the adaptive immune architecture of the lung (**Figure 6**). In this model, alveolar macrophages are not absent^29^ but instead fail to complete terminal differentiation and are redirected into inflammatory cell states characterized by loss of canonical alveolar macrophage identity, activation of NF-κB- and interferon-associated programs, and expression of lymphocyte-recruiting chemokines including CXCL9, CXCL10, CXCL13, and CXCL16.

**Figure 6.**
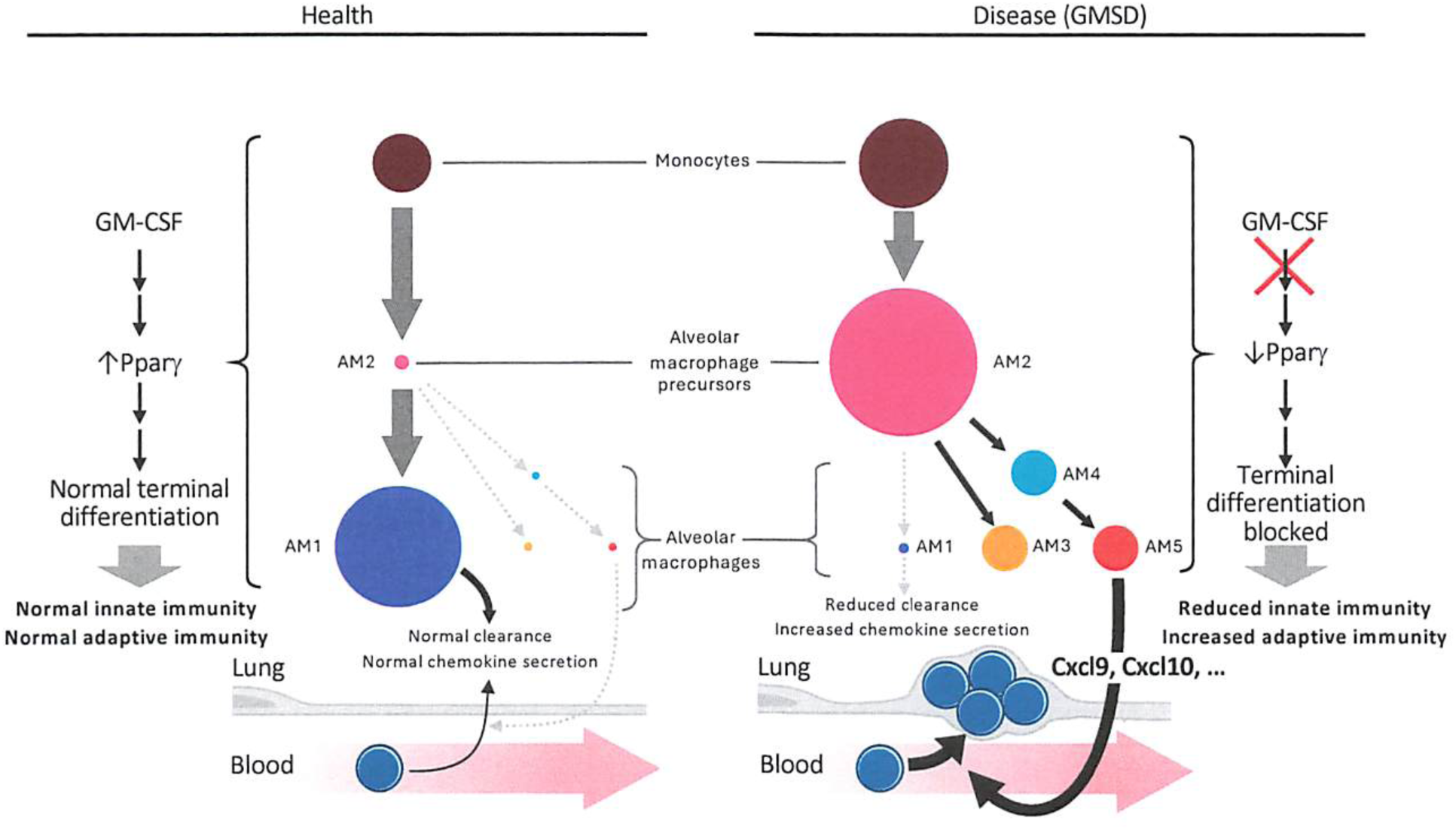
GM-CSF signaling maintains alveolar macrophage identity and prevents pulmonary lymphocytosis. In health, GM-CSF, via PPARγ, directs terminal alveolar macrophage (AM) differentiation through a transitional intermediate/precursor cell state (AM2) into the mature AM state (AM1). In the absence of GM-CSF signaling, AMs have reduced terminal differentiation into mature cells (AM1) and accumulate at the precursor cell state (AM2) and in alternative cell states (AM3-5). In two of these cell states (AM4-5), AMs are reprogramed into an inflammatory, chemokine-secreting state, which drives increased recruitment and accumulation of lymphocytes with distortion of peribronchovascular tissue.

These reprogrammed macrophages generate spatially organized chemokine gradients that promote accumulation of B cells, T cells, plasma cells, and macrophages within peribronchovascular regions, resulting in lymphoid follicle formation, local immunoglobulin production, and progressive pulmonary lymphocytosis. Pulmonary lymphocyte accumulation is driven by the combined effects of enhanced lymphocyte recruitment, in situ proliferation, and lymphoid lineage commitment, culminating in the development of tertiary lymphoid structures throughout the lung.

These findings support a mechanistic framework in which GM-CSF functions as a lineage-maintaining checkpoint that preserves alveolar macrophage identity and thereby restrains pathological adaptive immune responses in the lung. Under homeostatic conditions, GM-CSF-dependent transcriptional programs centered on PU.1, PPARγ, and C/EBP family members^26,29,45,47,57^ maintain a mature macrophage state that suppresses inflammatory chemokine expression. Loss of this program shifts alveolar macrophages from homeostatic sentinels to inflammatory organizers that actively recruit and sustain lymphocytes within the pulmonary microenvironment. The causal nature of this pathway is demonstrated by PMT, where selective restoration of GM-CSF responsiveness within the AM compartment re-established AM differentiation, suppressed chemokine production, and resolved pulmonary lymphocytosis. Thus, GM-CSF signaling deficiency links disruption of tissue-resident macrophage identity to tertiary lymphoid tissue formation and reveals alveolar macrophages as central regulators of pulmonary lymphocyte homeostasis rather than merely custodians of surfactant clearance and innate defense.

The implications of our findings likely extend well beyond GM-CSF signaling deficiency and PAP. Pulmonary lymphocytosis, tertiary lymphoid structure formation, and peribronchiolar immune aggregates are common pathological features of numerous chronic lung diseases, including hypersensitivity pneumonitis, autoimmune lung diseases, chronic obstructive pulmonary disease, severe asthma, interstitial lung diseases, and post-infectious fibrotic disorders.^6–17,58^ Our results suggest that, rather than arising solely from persistent antigenic stimulation or dysregulated lymphocyte-intrinsic pathways, these immune structures may also reflect a failure of tissue-resident macrophages to maintain local immune homeostasis. In this framework, alveolar macrophages function as active suppressors of adaptive immune organization, and chronic inflammatory, fibrotic, infectious, autoimmune, or aging-related insults that perturb macrophage identity may unleash chemokine programs that recruit and sustain lymphocytes within the lung. The observation that reprogrammed AMs generate coordinated chemokine networks capable of driving lymphocyte recruitment, proliferation, and differentiation raises the possibility that macrophage state transitions represent a common upstream mechanism linking diverse tissue injuries to tertiary lymphoid neogenesis. More broadly, these findings support a unifying model in which loss of tissue-resident macrophage identity, rather than loss of macrophage number,^7^ serves as a fundamental driver of chronic inflammation and adaptive immune remodeling across pulmonary diseases and perhaps in other organs where resident macrophages preserve tissue-specific immune equilibrium.

These findings have implications for the treatment of pulmonary lymphocytosis focused to prophylaxis. Pulmonary lymphocytosis arises upstream from disruption of a tissue-resident macrophage identity program and can be substantially reversed by restoring GM-CSF responsiveness selectively within the AM compartment. This observation raises the possibility that therapies aimed at preserving or re-establishing homeostatic AM differentiation may help limit the development of pathogenic lymphoid aggregates and tertiary lymphoid structures^6^ in diseases characterized by chronic pulmonary inflammation. More broadly, the results suggest that modulation of macrophage state, lineage-defining transcriptional programs, or downstream chemokine networks may represent complementary approaches for restoring immune homeostasis. Importantly, our findings do not imply that macrophage-directed therapies will be broadly effective across all forms of pulmonary lymphocytosis, where environmental, infectious, autoimmune, and genetic drivers are likely to contribute in varying degrees. Nevertheless, the ability of alveolar macrophage-targeted correction to normalize lymphocyte accumulation in this genetically defined model identifies tissue-resident macrophages as a potentially tractable therapeutic node and provides a rationale for exploring whether restoration of macrophage homeostatic function can modify disease progression in selected chronic lung disorders characterized by excessive adaptive immune remodeling.

An important implication of this study is the demonstration that AM replacement by PMT of GM-CSF-responsive macrophages can correct disease-driving immune dysregulation even in the setting of an intact but dysfunctional endogenous macrophage compartment. Historically, cell-based therapies for GM-CSF signaling deficiency and pulmonary alveolar proteinosis have been viewed primarily as a means to restore surfactant clearance and alveolar homeostasis.^51,59,60^ Our findings reveal a broader role: replacement of defective alveolar macrophages re-established the GM-CSF-dependent differentiation program, suppressed inflammatory chemokine production, reduced lymphocyte recruitment and proliferation, and restored pulmonary lymphocyte homeostasis. The success of pulmonary macrophage transplantation is particularly notable because it achieved these effects without directly targeting lymphocytes, immunoglobulin production, or other downstream manifestations of disease. Instead, correction of a relatively small population of tissue-resident sentinel cells was sufficient to remodel the broader pulmonary immune environment. These results provide proof-of-principle that replacement of genetically or functionally defective resident macrophages can durably reprogram tissue immunity and support the concept that macrophage-directed cellular therapies may have therapeutic value extending beyond restoration of macrophage housekeeping functions to include reconstruction of tissue-specific immune homeostasis.

### Limitations of the study

This study is subject to several limitations: results were derived exclusively from mouse models of GM-CSF signaling deficiency, the precise chemokines responsible for pulmonary lymphocyte recruitment remain unresolved, the relative contributions to pulmonary lymphocyte accumulation arising from recruitment, lineage commitment, and in situ proliferation remain unresolved, the antigenic targets of immunoglobulin production in GMSD mice and potential differences in the effects of developmental absence versus acquired loss of GM-CSF signaling were not addressed. Nevertheless, these findings establish GM-CSF-dependent alveolar macrophage programming as a fundamental mechanism governing pulmonary lymphocyte homeostasis and identify macrophages as central organizers of adaptive immune Reponses within the lungs. More broadly, our results reveal how disruption of a single macrophage differentiation pathway can reshape tissue immune architecture and drive chronic inflammatory disease.

## MATERIALS AND METHODS

### Animals

All animal procedures were approved by the Institutional Animal Care and Use Committee (IACUC) of Cincinnati Children’s Hospital Medical Center and were performed in accordance with institutional guidelines. B6.SJL-Ptprc^a^Pepc^b^/BoyJ (CD45.1; The Jackson Laboratory, stock no. 002014), *Csf2ra*^−/−^ *Csf2rb*^−/−^, and *Csf2*^−/−^ mice were maintained under specific pathogen-free conditions in the Cincinnati Children’s Hospital Research Foundation (CCHRF) vivarium. Age-matched male and female mice (12–16 weeks of age) were used for all experiments. The generation of *Csf2ra*^KO^, *Csf2rb*^KO^, and *Csf2^KO^* mice have been described previously.^23,30,31^

### Bronchoalveolar lavage and alveolar macrophage isolation

Alveolar macrophages were isolated from the lungs of mice by bronchoalveolar lavage (BAL) as previously described.^61^ Briefly, BAL fluid was collected from the lung, and cells were recovered by centrifugation at 300 x g for 10 min at room temperature. BAL turbidity and BAL cytology were determined as described previously.^51,60^ BAL cell pellets were resuspended in appropriate buffer and used for subsequent molecular analysis.^51,60^

### Mouse lung harvest for single cell suspension

Single cell suspension from mouse lungs were prepared as described.^62^ Briefly, mice were anesthetized with pentobarbital (50mg/kg body weight) via an intraperitoneal injection. The chest cavity was opened, and the right ventricle was perfused with 10ml of cold DPBS to remove circulating blood cells (red blood cells, T cells, B cells and monocytes) from the pulmonary vasculature and to accurately assess the lung immune populations. Following perfusion, the lung lobes were excised, cut into small pieces, and transferred to gentleMACS “C” tubes (Miltenyi Biotec, 130-096-334) containing 5 mL prewarmed digestion buffer consisting of 5U/mL dispase (Corning, 354235), 50U/mL DNase I (GoldBio, D-301), 480U/mL Collagenase type I (Gibco, 17100017) in DPBS. Lungs were dissociated using a gentleMACS Octo Dissociator (Miltenyi Biotec, 130-096-427) with the programs “m_lung_01_02” (36 seconds, 2 cycles), “37C_m_LIDK_1” (36 minutes and 12 seconds, once), and “m_lung_01_02” (36 seconds, once). The lung cell suspension was diluted with 40mL DPBS and was filtered through a 70-µm cell strainer and centrifuged at 300 × g for 10 min at 4°C. Sedimented cells were resuspended in ice-cold MACS buffer (Miltenyi Biotec) for processed for flow cytometry.

### Processing lung single cell suspension for Fluorescence-activated cell sorting (FACS)

Lung single cell suspensions (approximately 3×10^6^ cells) were incubated with Mouse BD Fc Block (BD Pharmingen, 553141; 1:100 dilution) for 10 min at room temperature, washed with MACS buffer, pelleted by centrifugation, and stained with the antibody panel listed in Supplementary Table 1. Cell viability was assessed using a Zombie™ Fixable Viability Dye (BioLegend). Dead cells were excluded from subsequent flow cytometric analyses based on Zombie dye positivity. Following staining, cells were washed, centrifuged, and filtered through 35-µm cell strainer caps (Falcon FACS tubes). Samples were acquired on a five-laser Cytek Aurora or Maleficient Aurora spectral flow cytometer. Unstained and single-stained controls were included for spectral unmixing. Spectral unmixing was performed using Spectro Flo software (Cytek Biosciences), and data were subsequently analyzed in SpectroFlo software following compensation adjustment and gating.

### Lung Histology and Immunohistochemistry

Mice were euthanized by pentobarbital overdose and exsanguination. Lungs were inflation-fixed via the trachea with 4% paraformaldehyde (PFA) in PBS (pH 7.4) at 25 cm H₂O pressure, ligated, and fixed at 4°C for 24 h. Lung lobes were then dissected, sectioned into ∼2-mm slices, paraffin embedded, and cut into 5-μm sections. Sections were stained with hematoxylin and eosin (H&E) and analyzed by light microscopy.^51^ Oil Red O staining and quantification of lipid accumulation in alveolar macrophages were performed as previously described.^37,63^ Immunohistochemical staining for B220 B220-Cd45 (Rabbit) Abcam 1:500 dilution, CD4 (SP35, Rabbit Ventana Roche), Pax5 (SP34, Rabbit Ventana Roche), CD68 9PG-M1 Abcam Ab 783 (1:50 dilution), Ki67 (anti-Ki67 antibody-Clone 30-9 (Roche Diagnostics, Basel, Switzerland) and GFP were performed by the CCHMC Integrated Pathology Research Facility. The slides were counter stained with species-specific secondary antibodies and detected with Opti-view DAB kit (Roche). Photographs were taken at 20X and 40X magnification using Leica DM2700 M brightfield microscope (Leica Microsystems, Buffalo, Grove, IL).

### Preparation of bone marrow derived macrophages and gene corrected macrophages

BMDMs and *Csf2ra* gene-corrected macrophages were prepared from hematopoietic progenitor cells and differentiated *in vitro* as previously described.^51,60^

### Pulmonary macrophage transplantation

BMDMs or LSK-derived macrophages were delivered to the lungs of 8–10-week-old mice by endotracheal instillation as previously described.^51,60^ Briefly, macrophages were resuspended in 50 μL PBS and were administered under isoflurane anesthesia and inhaled into the lungs during spontaneous respiration.

### Histopathological Scoring of Pulmonary Lymphocytosis

Pulmonary lymphocytosis was evaluated on lung tissue sections using a semiquantitative scoring system based on the extent and distribution of peribronchovascular lymphocyte accumulation. Each lymphocytic focus was assigned a score ranging from 0 to 4: 0, no detectable peribronchovascular lymphocyte accumulation; 1, mild accumulation characterized by minimal peribronchovascular lymphocyte infiltration; 2, moderate accumulation consisting of multiple independent or continuous foci adjacent to a vessel; 3, severe accumulation defined by extensive perivascular lymphocyte infiltration surrounding a vessel or bridging adjacent vessels; and 4, extensive lymphocyte accumulation involving the alveolar space. For each lung section, all lymphocytic foci were evaluated, and individual scores were summed to generate a cumulative lymphocytosis score for that section. Cumulative section scores were then used for statistical comparisons between experimental groups. Histopathological scoring was performed by investigator(s) blinded to experimental group allocation and genotype. The distribution of lymphocytosis scores within each group was assessed for normality and homogeneity of variance prior to statistical testing.

### Electron microscopy

For transmission electron microscopy (TEM), mice were anesthetized and inflated as described previously (Suzuki et al 2014) using freshly prepared Karnovsky’s fixative solution containing 2.5% glutaraldehyde, 2% paraformaldehyde, and 0.1 M sodium cacodylate buffer (pH 7.3). These fixed lungs were submitted to the CCHMC Integrated Pathology Research Facility for processing for TEM.^64^ Images were acquired using a Hitachi H-7650 transmission electron microscope.

### ELISA

Concentrations of Gpnmb, Cxcl9, Cxcl10, Cxcl13, GM-CSF, M-CSF, IL-34 and TGFβ in bronchoalveolar lavage (BAL) fluid were quantified using commercially available ELISA kits (Quantikine®, R&D Systems) according to the manufacturer’s instructions. Cytokine concentrations were calculated from standard curves generated using recombinant standards and are reported as pg/mL.

### Measurement of GM-CSF-stimulated STAT5 phosphorylation

GM-CSF-induced STAT5 phosphorylation in BAL cells was assessed by phospho-flow cytometry. Cells were stimulated with recombinant murine GM-CSF (10 ng/mL) for 30 min at 37°C and subsequently fixed, permeabilized, and stained with an anti-phospho-STAT5 (pY694) antibody (clone 47/Stat5; BD Biosciences) according to the manufacturer’s instructions and as previously published (Suzukji 2014, Arumugam 2019, 2024). Phosphorylated STAT5 (pSTAT5) expression was quantified by flow cytometry using an LSR Fortessa™ flow cytometer (BD Biosciences) at the CCHMC Research Flow Cytometry Core. Data was analyzed using FCS Express Research Edition software.

### Measurement of BAL fluid chemokines, biomarkers and immunoglobulins

Concentrations of chemokines, cytokines and immunoglobulins in bronchoalveolar lavage (BAL) fluid were quantified using a magnetic bead-based multiplex assay (Mouse Magnetic Luminex® Assay, R&D Systems; catalog no. LXSAMSM-22 and LXSAMSM-3, respectively) according to the manufacturer’s instructions. BAL fluid was separated from cells by centrifugation (285 × g, 10 min) and stored at −20°C until analysis. Samples were analyzed on a Luminex/MILLIPLEX analyzer (MilliporeSigma), and cytokine concentrations were determined from standard curves generated using recombinant cytokine standards. Results are reported as pg/mL.

### In vitro-cytokine stimulated proliferation assay

Bone marrow-derived macrophages (BMDMs) from WT and GM-SD mice were plated at 2.5 × 10^4^ cells per well and cultured for 72 h in serum-free macrophage medium (Macrophage-SFM; Gibco) in the presence or absence of recombinant mouse GM-CSF or M-CSF (10 ng/mL. Cell proliferation was quantified using an XTT-based colorimetric assay (Cell Proliferation Kit II, Roche Applied Science) according to the manufacturer’s instructions. Proliferation was expressed as the fold change in absorbance relative to cells cultured without cytokine supplementation. Cell viability was >95% prior to plating, as determined by trypan blue exclusion.

### Alveolar macrophage cell lines

Three previously described murine alveolar macrophage cell lines were used.^26^ WT MΦ cells were derived from *Csf2*^+/+^ mice and exhibit a mature phenotype.^65^ (*Csf2*^−/−^PU.1^neg^ cells were derived from *Csf2*^−/−^ mice and display an immature phenotype lacking spontaneous PU.1 expression. ^26^*Csf2*^−/−^rPU.1 overexpressing cells were generated by retroviral expression of PU.1 in *Csf2*^−/−^PU.1^Neg^ cells and exhibit a mature alveolar macrophage phenotype.^26^

### Quantitation of mRNA transcript abundance

Total RNA (1µg) was combined with 1 μg oligo(dT) primer (Life Technologies) in a final volume of 12 μL. Samples were denatured at 65°C for 10 min and immediately cooled on ice. Reverse transcription was performed by adding 8 μL of cDNA synthesis master mix containing 4 μL First-Strand Buffer, 2 μL 25 mM MgCl₂, 1 μL 10 mM dNTP mix, 0.5 μL SuperScript II reverse transcriptase, and 0.5 μL RNaseOUT (Life Technologies), followed by incubation at 42°C for 60 min. Reactions were terminated at 95°C for 5 min, and cDNA was stored at −80°C until analysis. *Csf1* and *Gapdh* transcript abundance was quantified by qRT-PCR. *Csf1* transcript levels were normalized to the mean *Gapdh* transcript number for each sample. Nascent *Csf1* transcription was calculated as the difference between normalized *Csf1* transcript abundance measured in RNA isolated from nuclei incubated in the presence versus absence of exogenous NTPs.

### Quantitative Reverse Transcription-Polymerase chain reaction (qRT-PCR)

Total RNA was isolated from bronchoalveolar lavage (BAL) cells using TRIzol reagent (Life Technologies, Carlsbad, CA, USA) according to the manufacturer’s instructions. First-strand cDNA was synthesized using the SuperScript III First-Strand Synthesis System (Thermo Fisher Scientific, Waltham, MA, USA). qRT-PCR was performed using 100 ng of cDNA template per reaction to determine relative gene expression levels of *Lgals3*, *Mrc1, Chil3, Lpl, Car4, Bach2, Cebpa, Cebpb, Spi1, Pparg, NFkb1, STAT1, Sfln4, Apoe, Gpnmb, Cxcl9, Cxcl10, Cxcl16,* and *Csf1r* were measured using Quant Studio 3 and analyzed using the Quant Studio^TM^ design and analysis software V1.5.2. (Applied Biosystems by Thermo Fisher scientific). The details of the primer/probe sets (Thermo Fisher Scientific) are included in **Supplementary table 2**. Target gene expression was normalized to the 18S RNA housekeeping gene and analyzed using the comparative Ct (ΔΔCt) method.

### Surfactant clearance assay

Surfactant clearance of bone marrow derived macrophages was assessed using a previously described.^66^

### Trafficking of adoptively transferred donor lymphocytes

Spleens were harvested aseptically from C57BL/6-Tg(UBC-GFP)30Scha/J ubiquitin-GFP transgenic mice and mechanically dissociated in Dulbecco’s modified Eagle medium (DMEM). Single-cell suspensions were generated by passage through a 100-µm cell strainer, followed by erythrocyte lysis using ammonium chloride-based lysis buffer for 5 min at room temperature. Cells were washed, resuspended in phosphate-buffered saline (PBS) at a concentration of 2 × 10^8 cells per 0.2 mL, and maintained on ice until adoptive transfer. Wild-type (WT) and *Csf2*^-/-^recipient mice were anesthetized with isoflurane and intravenously injected with 2-3 × 10^7^ GFP^+^ splenocytes via the retroorbital injections. Twenty-four hours after cell transfer, mice were euthanized and lungs were harvested for preparation of single-cell suspensions as described above. The frequency of donor-derived GFP^+^ cells in lung tissue were determined by flow cytometry.

### Western blotting

Primary alveolar macrophages (5 × 10^5^ cells per condition) or bone marrow-derived macrophages (BMDMs; 1 × 10^6^ cells per condition) were stimulated with GM-CSF or M-CSF (10 ng/ml) for 30 min, harvested by centrifugation, and lysed on ice for 30 min in lysis buffer containing 50 mM Tris-HCl (pH 8.0), 150 mM NaCl, 1% Nonidet P-40, 0.5% sodium deoxycholate, and 0.1% SDS supplemented with protease and phosphatase inhibitors. Lysates were clarified by centrifugation (10,000 × g, 15 min, 4 °C), mixed with Laemmli sample buffer, heated at 95 °C for 5 min, and resolved by SDS–PAGE on 4–12% Bis-Tris gradient gels (Invitrogen) under reducing conditions. Proteins were transferred to PVDF membranes and blocked overnight at 4 °C in TBST (50 mM Tris-HCl, pH 8.0, 150 mM NaCl, 0.1% Tween-20) containing 5% non-fat milk. Membranes were incubated with primary antibodies for 2 h at room temperature, washed with TBST, and subsequently incubated with horseradish peroxidase-conjugated secondary antibodies for 1 h at room temperature. Immunoreactive bands were detected using enhanced chemiluminescence (ECL Plus, GE Healthcare). The following primary antibodies were used: anti-GM-CSF receptor β (1:500, Santa Cruz Biotechnology), anti-STAT5 (1:500, Santa Cruz Biotechnology), anti-phospho-STAT5 (1:500, Upstate), anti-ERK1/2 (1:1,000, Cell Signaling Technology), anti-phospho-ERK1/2 (1:1,000, Cell Signaling Technology), and anti-β-actin (1:1,000, Santa Cruz Biotechnology).

### M-CSF secretion assay

Primary alveolar macrophages (5 × 10^5^ cells per well) or cultured macrophage cell lines (1 × 10^6^ cells per well) were cultured in fresh complete DMEM (DMEM, 10% FBS, 1% Penicillin/Streptomycin) for 24 h following two washes with PBS. M-CSF concentrations in culture supernatants were measured by ELISA as described above.

#### Tissue dissociation for single nuclear RNA sequencing

Murine lungs were harvested and immediately snap-frozen in liquid nitrogen until processing. Individual lungs were dissociated into single-nucleus suspensions using gentleMACS “C” tubes in gentleMACS tissue dissociator in ice-cold lysis/fixation buffer containing 10 mM NaCl, 10 mM sodium phosphate (pH 7.2), 3 mM MgCl₂, 5% glutaraldehyde, 10 mM vanadyl ribonucleoside complex, 0.1% Triton X-100, 1% diethyl pyrocarbonate, and 0.00015% polyvinyl sulfonic acid, as previously described.^67^ Nuclei were washed and resuspended in nuclei suspension buffer (10 mM Tris-HCl, pH 7.4, 10 mM NaCl, and 3 mM MgCl₂). Isolated nuclei were snap-frozen in nuclei suspension buffer for single-cell combinatorial indexing RNA sequencing (sci RNA-seq) and stored at −80 °C until use. A subset of nuclei was retained for quality assessment and quantification using a hemocytometer.

#### Generation of Sci RNA sequencing library

Single-nucleus RNA sequencing (snRNA-seq) libraries were generated from fixed nuclei using a two-level combinatorial indexing strategy as previously described.^68,69^ Libraries were prepared in three independent batches, each containing seven to nine samples. Reverse transcription was performed using a modified ramped incubation protocol consisting of 2 min at each of 4 °C, 10 °C, 20 °C, 30 °C, 40 °C, and 50 °C, followed by 10 min at 53 °C and 15 min at 55 °C. Nuclei were distributed across reverse transcription wells, and nuclei were sorted into each PCR well following reverse transcription as previously described.^67^ Libraries were amplified, pooled, and sequenced on an Illumina NextSeq 550 platform according to the manufacturer’s instructions.

#### snRNA-Seq data processing

Raw FASTQ files were processed into gene-by-cell count matrices using the BBI pipeline with default parameters.^68^ Low-quality nuclei were excluded based on the following criteria: >100 unique molecular identifiers (UMIs), <10% mitochondrial transcript content, and classification as a doublet by Scrublet (doublet score >0.2) (*Wolock SL, 2019 Cell Systems*). Dimensionality reduction, clustering, and visualization were performed using Monocle3 (v1.2.8).^69^ (Prior to downstream analysis, datasets were aligned across batches, genotypes, and the logarithm of total UMIs per nucleus using the align_cds function. Principal component analysis (PCA) was used for dimensionality reduction, followed by Uniform Manifold Approximation and Projection (UMAP) for visualization and Leiden community detection for clustering. Cell identities were assigned by comparing cluster-specific marker genes with published reference markers. Major cell populations were subsequently divided into subsets and reanalyzed at higher resolution through iterative UMAP projection and Leiden clustering to define cellular subpopulations. Sftpb+ endothelial and immune cells were considered as potentially contaminated cells and were excluded from the downstream analysis.

#### snRNA-Seq differential expression analysis

Differential expression (DE) tests between cell groups were performed using Seurat (v5) using LogNormalized data and FindMarkers with two-tailed Wilcoxon rank sum test.

To identify robust differentially expressed genes (DEGs) between GMSD and WT within each major cell group, we performed a DE test comparing gene expression between WT and each of the three GMSD genotypes and considered a DEG as robust if it passed significance criteria (p<0.05, fold change>=1.5, and pct>=0.2) in at least two of the GMSD genotypes when compared to WT.

To identify robust DEGs for an alveolar macrophage (AM) cluster, we performed a DE test comparing gene expression in the cluster cells and in all other AM cells within each of the four genotypes and considered a DEG as robust if it passed significance criteria (p<0.05, fold change>=1.5, and pct>=0.2) in the DE tests in at least two genotypes.

### KEGG pathway enrichment analysis

Differentially expressed genes identified for each cell population were subjected to Kyoto Encyclopedia of Genes and Genomes (KEGG) pathway enrichment analysis using Enrichr (Ma’ayan Laboratory). Gene lists were uploaded to the Enrichr platform and queried against the KEGG 2021 Mouse gene set library. Pathways with a Benjamini-Hochberg-adjusted *P* value < 0.05 were considered significantly enriched. Enrichment results were ranked by adjusted *P* value and combined score, and representative pathways were visualized in the figures.

#### snRNA-seq cell type abundance

Relative abundance of cell types across genotypes was calculated using Multivariate regressions with Poisson lognormal models as implemented in Hooke, v0.0.1 and differences in cell numbers per type per genotype were identified with beta binomial testing, as implemented previously.^70^

### snRNA-seq trajectory analysis of myeloid cells and gene module identification

Pseudotime trajectory analysis of monocyte and macrophage cell clusters was performed using Monocle3. We built the cell trajectories using only WT cells, only GMSD cells, and both WT and GMSD cells together and the starts of the trajectories were set at the graph termination points in monocytes (cells expected from a literature review to be at the beginning of myeloid differentiation in the injured lung). Gene modules representing each cluster were identified using Monocle 3 based on the pseudotime cell trajectories. For each of the inferred cell trajectories, we first identified gene expression detected in at least 50 cells and significantly changed (q value<0.05 of Monocle3 graph_test function) along with the pseudotime in the trajectory. After this gene identification for all paths, we combined all significantly changed genes, performed a clustering analysis to participate genes into distinct modules based on expression using Monocle3 find_gene_modules function, and then assigned a module to a cluster if it satisfied the following two criteria: (1) the cluster has the highest score (collective expression of the genes in the module) of this module when compared to the scores of the module in all the other clusters, and (2) the module score in the cluster is at least 1.2 fold greater than the second highest score of this module in all other clusters. After this module-to-cluster assignment, we filtered genes from each module if the expression percentage is less than 20% in the assigned cluster of the module. We first used the above approach to identify candidate gene modules for each cluster in WT and GMSD separately, then combined all the genes in the candidate modules, and repeated the module identification (using the find_gene_modules function) and module-to-cluster assignment analysis using the combined WT and GMSD data to obtain the final representative gene module for each cluster.

#### snRNA-seq gene signature analysis

Monocyte and macrophage subtypes were assessed for M1 or M2 signature scores using gene sets acquired from previously.^71–74^ Genes expressed in less than 50 cells were excluded. Seurat’s AddModuleScore function was used to calculate a M1 or M2 geneset score for each cell.

### 10x Visium HD spatial transcriptomics Profiling of Mouse Lung

Spatial transcriptomic profiling was performed using the Visium HD Spatial Gene Expression platform (10x Genomics) according to the manufacturer’s protocols. Formalin-fixed paraffin-embedded (FFPE) lung sections were deparaffinized, stained with hematoxylin and eosin (H&E), and imaged using a BZ-X800 microscope (Keyence) equipped with a 20× PlanApo objective (NA 0.75). Whole-slide images were acquired using automated focus with five z-slices and stitched using BZ-X800 Analyzer software without additional image processing. Following imaging, tissue sections were processed using the Visium HD FFPE workflow. Briefly, sections were destained, decrosslinked, and hybridized with the Visium HD Mouse Whole Transcriptome Probe Panel (10x Genomics). Following probe hybridization and ligation, tissue sections and Visium HD slides were loaded onto the Visium CytAssist instrument, where ligated probe products were released from the tissue and captured by spatially barcoded oligonucleotides on the Visium HD slide. Captured molecules were amplified and used for library preparation according to the manufacturer’s instructions. Libraries were purified using SPRIselect beads, assessed for quality using an Agilent 4150 TapeStation, quantified, pooled, and sequenced on a NovaSeq X sequencer (Illumina) using a single lane of a 10B flow cell. Sequencing was performed with 43 cycles for Read 1, 10 cycles each for the i7 and i5 index reads, and 50 cycles for Read 2. Spatial barcodes and unique molecular identifiers (UMIs) were encoded in Read 1, and transcript-derived sequences were captured in Read 2.

### 10x Visium HD spatial transcriptomic data analysis

The generated 10x Visium HD data was processed using 10x Genomics Space Ranger pipeline (v3.0.1) using 10x mouse mm10 reference genome and Mouse Transcriptome Probe Set v2.0. Downstream QC and analysis were performed using 8μm-bin resolution data using Seurat (v5.0). Bins with less than 10 UMIs were excluded. Data normalized was performed using Seurat’s LogNormalize method. Spatial patterns of gene expression were visualized using the SpatialFeaturePlot in Seurat. We selected regions of interest (ROI, each is a circled area with 640μm in diameter) in both WT and *Csf2ra*^KO^ Visium HD data based on histopathologic features identified on matched H&E images and overlaid spatial gene expression maps. 8 ROIs were selected for each condition. We created a pseudo-bulk gene expression profile for each ROIs by summing gene expression in the bins within each ROI and performed principal component analysis and differential expression (DE) analysis using DESeq2 pipeline. Genes with an adjusted *P* value < 0.1 and an absolute log2 fold change > 1 were considered significant. Statistics of the DE test was visualized as volcano plot using the EnhancedVolcano package in R, and the expression patterns of DEGs across pseudo-bulk profiles were visualized using heatmap.

### Statistical Analysis

Data are presented as mean ± s.e.m. Normality was evaluated using the Shapiro–Wilk test. Two-group comparisons were performed using a two-tailed Student’s *t*-test for normally distributed data and a two-sided Mann–Whitney *U* test for data that were not normally distributed. Statistical significance was defined as *P* < 0.05. Experiments were independently repeated at least twice with similar results. The exact value of *n* is provided in the corresponding figure legends. Group comparisons were performed using an unpaired two-tailed Student’s T-test for normally distributed data with equal variance or a Mann-Whitney *U* test for nonparametric data. Differences were considered statistically significant at *P* < 0.05.

## ACKNOWLEDGEMENTS

The authors would like to thank the Genomics Sequencing Facility, Research Flow Cytometry Facility and Integrated Pathology Research Facility of the Cincinnati Children’s Research Foundation, all of whom have provided extensive technical support. We also thank Madeline Bonfield and Jerilyn Gray for their support with flow cytometry panel design and data acquisition. This work was Supported by grants from the National Heart, Lung, and Blood Institute (NHLBI) R01 HL118243 (B.C.T. and C. T.), R01 HL85453 (B.C.T.), R01 HL149743 (P.A.) and 5U24 HL134763 subaward 1701192 (P.A.), Chan-Zuckerburg Initiative DAF2020-217687 (C.T. and B.C.T.) and Translational Pulmonary Science Center and Division of Pulmonary Biology, CCHMC.

## Author Contributions

BCT, PA, and CT designed the study. PA, CW, DR, MM, SS, MW, YM, TS, RK performed research. CW, CT, PA, MG, RK and BT analyzed data. PA and BCT wrote the initial draft of the manuscript. PA, CW, DR, SS, MW, BC, GW, CT, and BCT revised and approved the final manuscript.

