## Supplementary figures and images for "GM-CSF Programs Alveolar Macrophages to Maintain Pulmonary Lymphocyte Homeostasis"

### Supplemental Figures

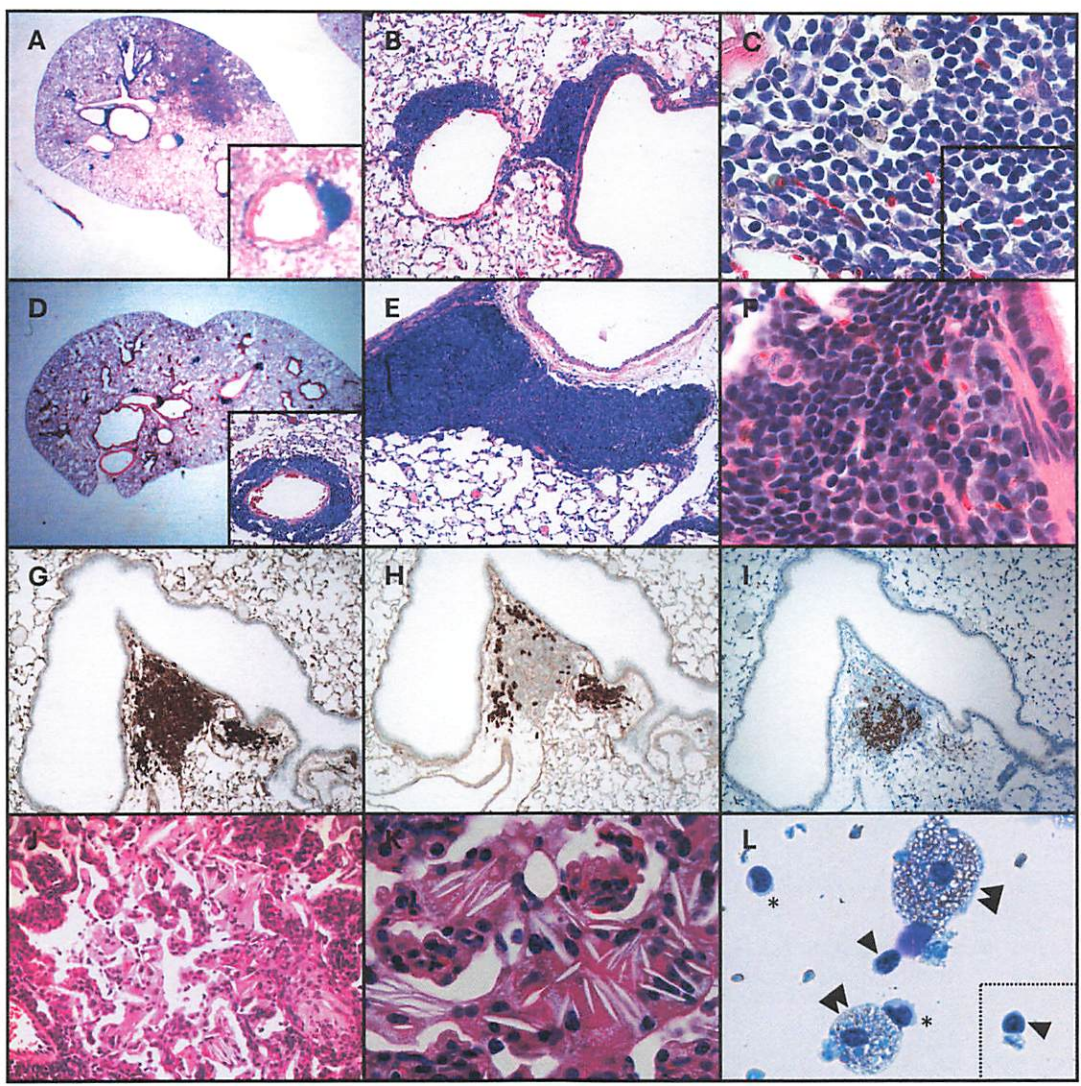

Figure S1

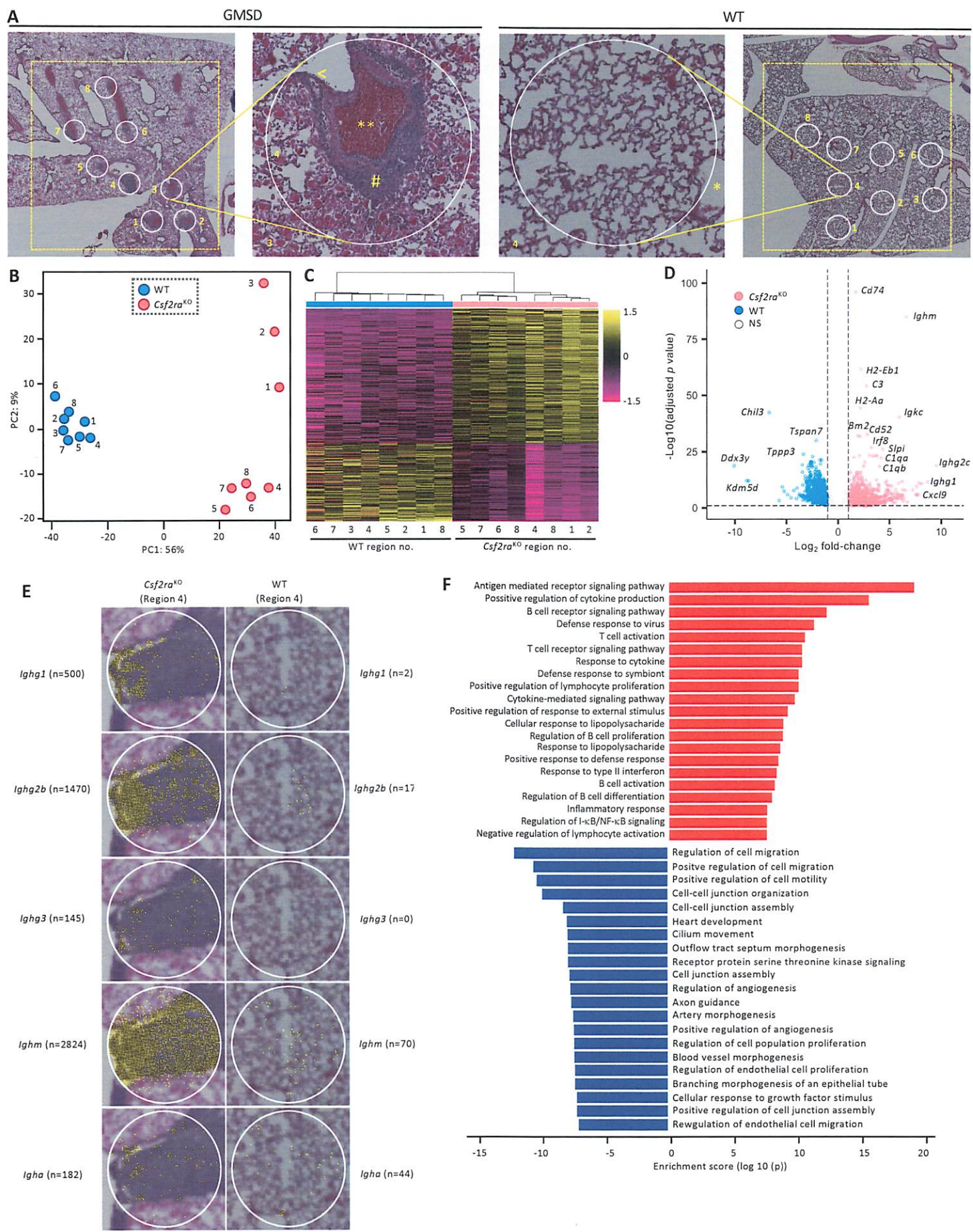

Figure S2

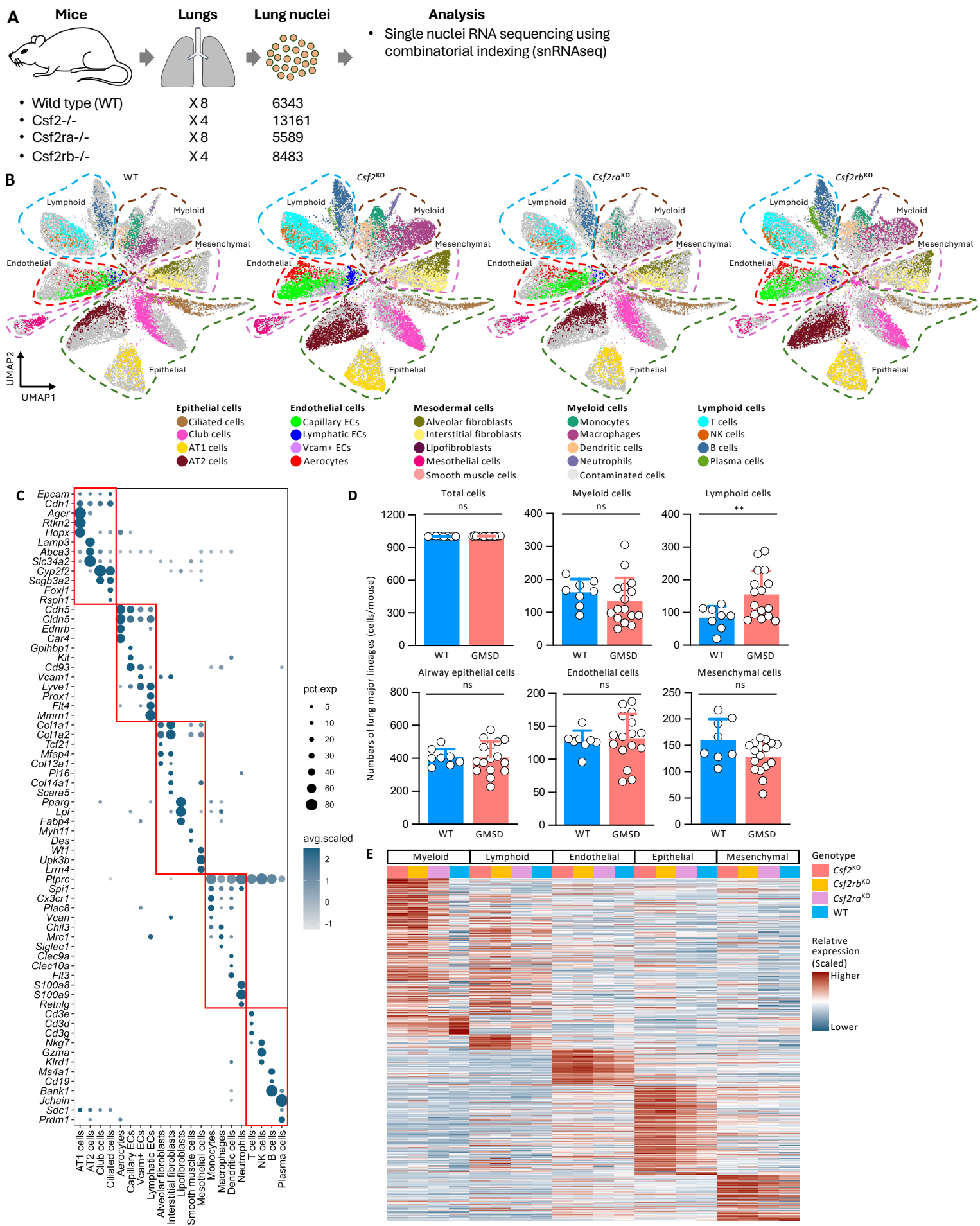

Figure S3

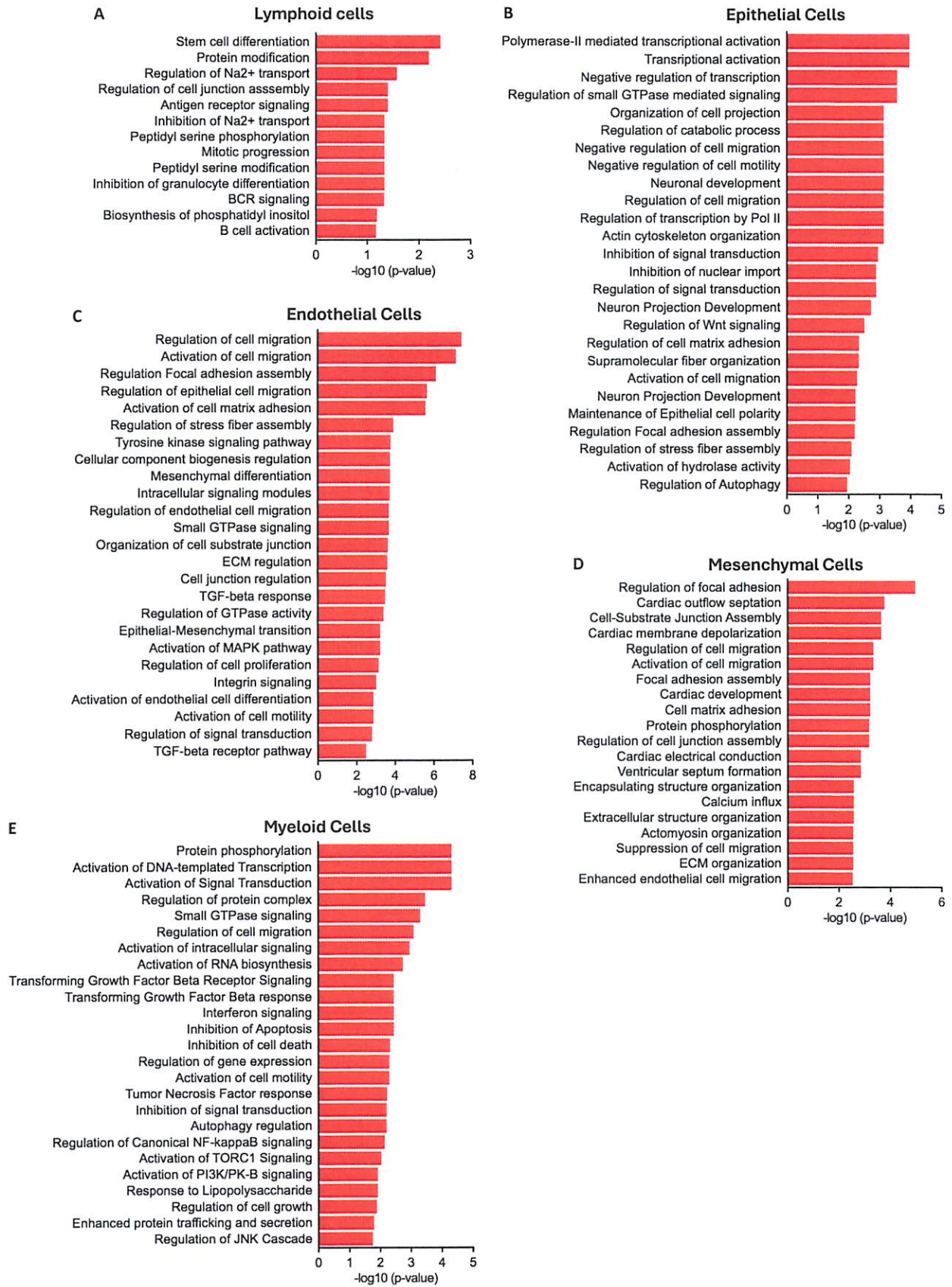

Figure S4

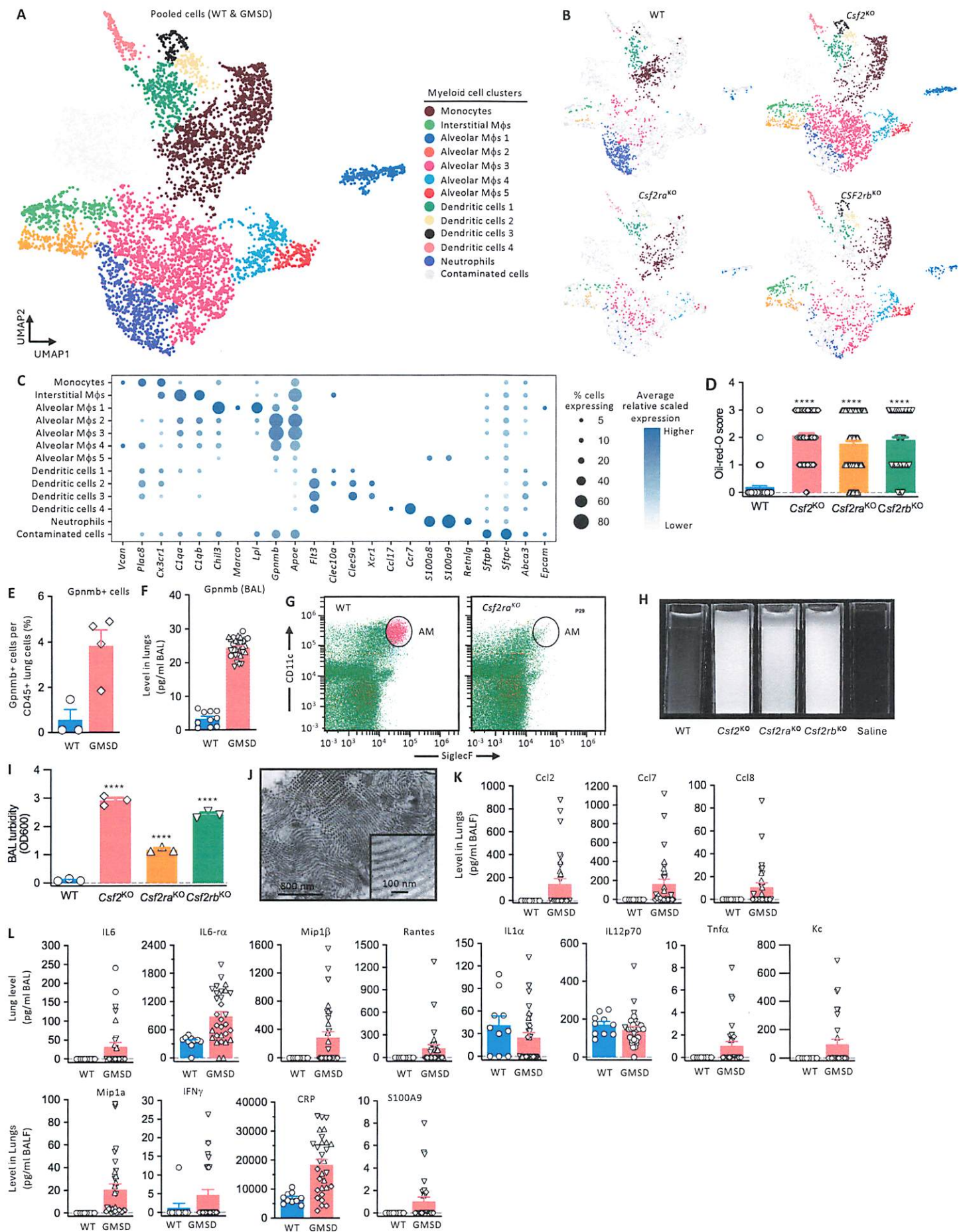

Figure S5

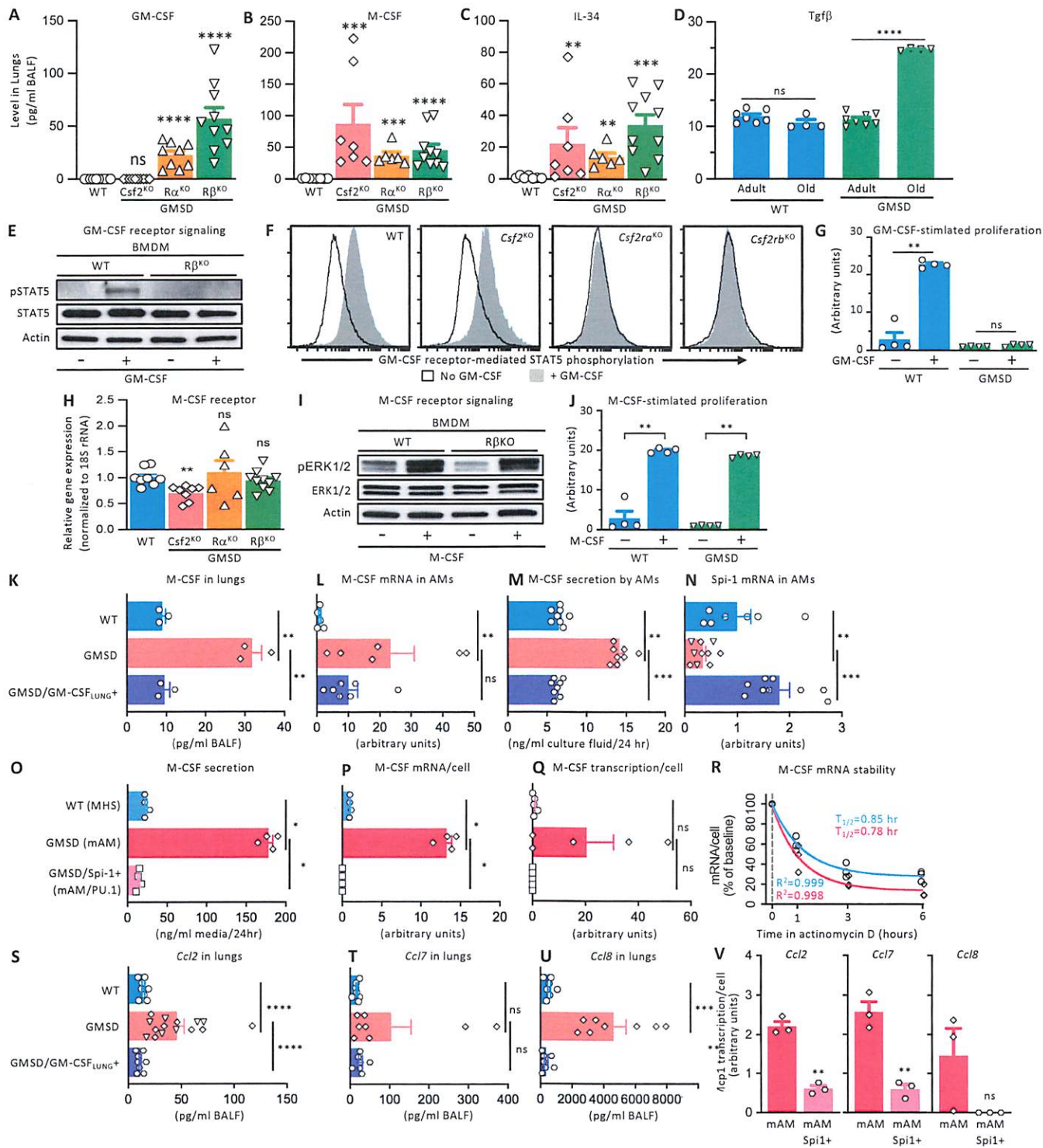

Figure S6

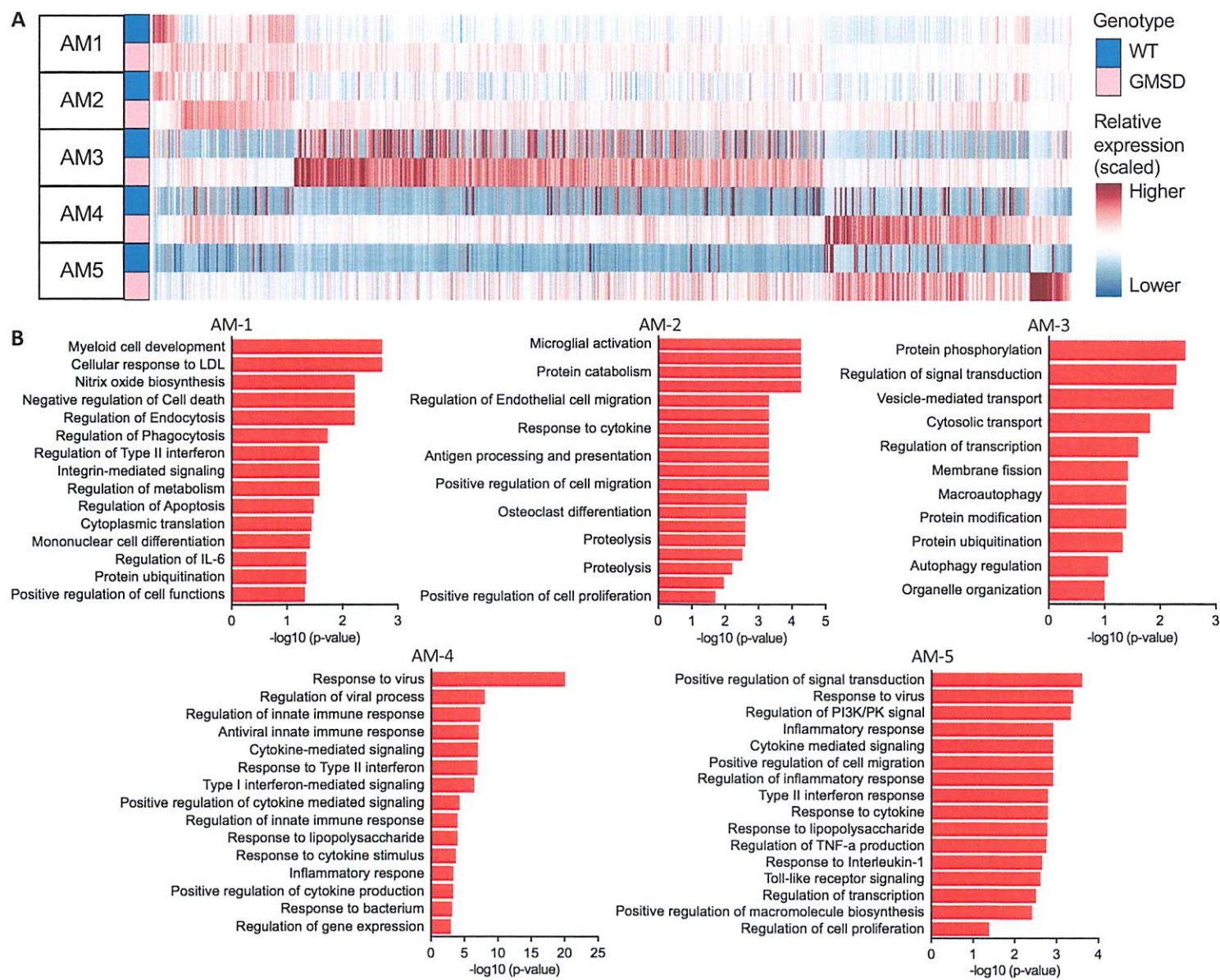

Figure S7

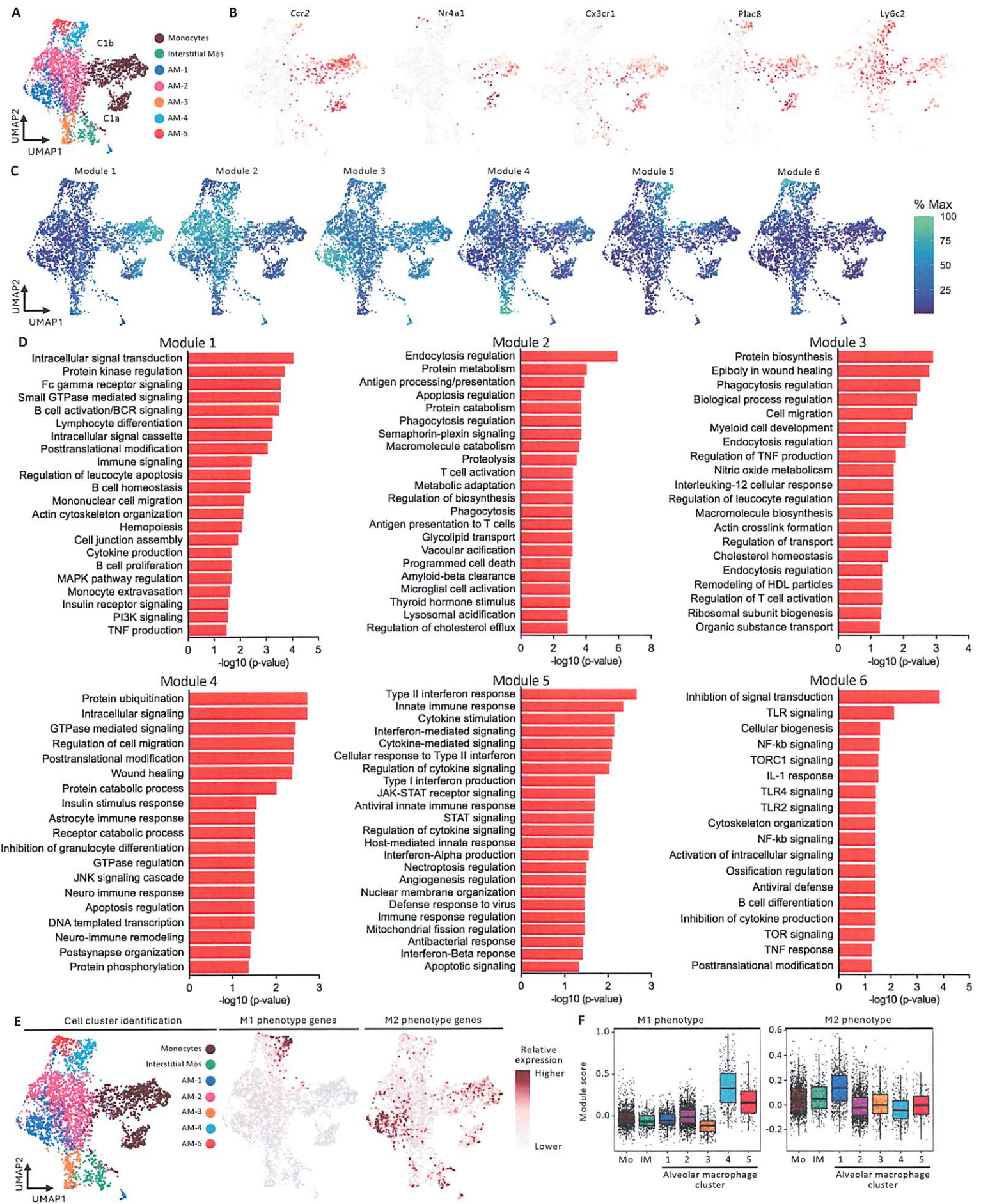

Figure S8
