## Supplemental text for "GM-CSF Programs Alveolar Macrophages to Maintain Pulmonary Lymphocyte Homeostasis"

### SUPPLEMENTAL FIGURE LEGENDS

#### Figure S1. Validation of pulmonary lymphocytosis and alveolar proteinosis phenotypes in GM-CSF signaling-deficient mice.

Representative histopathological features observed in *Csf2*<sup>-/-</sup>, *Csf2ra*<sup>-/-</sup>, and *Csf2rb*<sup>-/-</sup> (GMSD) mice.

(A-F) H&E-stained lung sections demonstrating progressive peribronchovascular immune cell accumulation (Magnification- A, D - 1x, B-20x, E-10x C, F, I -63x). Immunostaining for B220 (G), CD3 (H), Ki67 (I), Magnification-20x. C, F- High-power images show infiltrates composed predominantly of lymphocytes with occasional macrophages. (J and K) Intra-alveolar surfactant accumulation and cholesterol crystals in *Csf2ra*<sup>-/-</sup> lungs, Magnification 63x. (L) Diff-Quik staining of bronchoalveolar lavage (BAL) cells showing foamy alveolar macrophages, monocytes, and lymphocytes, Magnification-20x. Single arrowheads: lymphocytes; double arrowheads: foamy alveolar macrophages.

#### Figure S2. Spatial transcriptomic characterization of peribronchovascular lymphoid accumulations.

Spatial transcriptomic analysis identifies transcriptional remodeling within peribronchovascular immune aggregates in GMSD lungs. (A) Regions (1-8) selected for spatial transcriptomic profiling in WT and *Csf2ra*<sup>-/-</sup> lungs are marked in white circles. (B) Principal component analysis of spatially profiled regions. (C and D) Differential gene expression between WT and *Csf2ra*<sup>-/-</sup> peribronchovascular regions visualized by heatmap and volcano plot. (E) Spatial expression of immunoglobulin genes within lymphoid accumulations. 'n' represents number of bins positive for specific immunoglobulin expression. (F) KEGG pathway enrichment analysis of differentially expressed genes. < denotes pleural space, \* denotes airway, \*\* denotes blood vessel, # denotes peribronchovascular lymphocyte accumulations.

#### Figure S3. Extended quality control and cellular annotation of the lung snRNA-seq atlas.

Single-nucleus RNA sequencing defines cellular alterations across GM-CSF signaling-deficient lungs. (A) Experimental workflow. (B) UMAP visualization of integrated nuclei from WT, *Csf2*<sup>-/-</sup>, *Csf2ra*<sup>-/-</sup>, and

*Csf2rb*<sup>-/-</sup> lungs. (C) Marker gene expression used to identify major lung cell populations. (D) Quantification of epithelial, endothelial, mesenchymal, myeloid, and lymphoid populations across genotypes normalized to 1000 cells per mouse. (E) Heatmap showing lineage-specific differential gene expression. Each symbol represents data from one mouse. Data shown are the means ± SEM. Significance was assessed using Mann-Whitney U tests; NS-not significant, \*\*$P < 0.01$.

**Figure S4. KEGG pathway analysis of major lung cell lineages.**

Pathway enrichment analysis reveals lineage-specific transcriptional responses to loss of GM-CSF signaling. (A-E) KEGG pathway enrichment analyses for epithelial, endothelial, mesenchymal, myeloid, and lymphoid cell populations identified by snRNA-seq.

**Figure S5. Inflammatory cytokine, chemokine, and alveolar macrophage abnormalities in GMSD mice.**

Loss of GM-CSF signaling is associated with alveolar macrophage dysfunction and pulmonary inflammation. (A-B) UMAPs showing integrated and genotype-specific analyses of pulmonary myeloid cell populations. (C) Marker gene expression identifying individual myeloid subsets. (D) Quantification of Oil Red O-positive BAL cells. (E-F) Quantification of alveolar macrophages positive for *Gpnmb* expression measured by flow cytometry and soluble *Gpnmb* concentrations measured by ELISA in BAL fluid, respectively. (G) Representative flow plot of SiglecF<sup>+</sup>CD11c<sup>+</sup> cells in lung digest of WT and GMSD mouse. (H-I) Representative images and quantification of turbidity in BAL fluid of WT and GMSD mice. (J) Electron microscopy image demonstrating accumulation of surfactant-rich material within alveolar spaces, inset showing ultrastructure of surfactant in GMSD mice. (K-L) BAL inflammatory mediators, cytokines, and chemokines measured by Luminex assay.

**Figure S6. Enhanced M-CSF signaling and chemokine production in response to defective GM-CSF signaling.**

Defective GM-CSF signaling induces M-CSF expression and downstream inflammatory responses. (A-D) BAL concentrations of GM-CSF, M-CSF, IL-34, and TGF-β. (E and F) GM-CSF-induced STAT5 phosphorylation in bone marrow-derived macrophages (BMDMs) measured by immunoblotting and flow cytometry. (G) GM-CSF-dependent BMDM proliferation (n=4/condition). (H) Expression of *Csf1r* in AMs. (I and J) M-CSF receptor signaling and M-CSF-dependent proliferation in BMDMs. (n=4/condition) (K-M) Pulmonary M-CSF expression in WT, GMSD and GMSD mice expressing GM-CSF only in the lungs. (N) *Spi1* gene expression in AMs-(O-Q) Regulation of M-CSF expression in primary macrophage cell lines. (R) Analysis of *Csf1* mRNA stability (n=3 determinations/line). (S-U) Levels of CCL2, CCL7, and CCL8 in mice. (V) Chemokine gene expression in GMSD macrophage cell line expressing PU.1. Data shown are the means ± SEM. Significance was assessed using Mann-Whitney U tests; NS-not significant, \*P<0.05, \*\*P<0.01, \*\*\*P<0.001 and \*\*\*\*P<0.0001.

**Figure S7. Differential gene expression and pathway enrichment across alveolar macrophage states.**

Distinct alveolar macrophage populations emerge following loss of GM-CSF signaling. (A) Heatmap showing transcriptional signatures of alveolar macrophage clusters identified in WT and GMSD lungs. (B) KEGG pathway enrichment analysis of differentially expressed genes defining individual macrophage clusters. Abbreviations: GMSD=GM-CSF signaling deficient (i.e., *Csf2*<sup>-/-</sup>, *Csf2ra*<sup>-/-</sup> and *Csf2rb*<sup>-/-</sup>), WT=wild type. N=4-8 mice/genotype), AM-alveolar macrophages.

**Figure S8. Transcriptomic characterization of monocyte and alveolar macrophage populations in GMSD lungs.**

Integrated analysis of monocyte and alveolar macrophage populations reveals altered differentiation and activation programs in GMSD mice. (A) Integrated UMAP of monocyte and alveolar macrophage populations. (B) Expression of canonical monocyte-associated genes across clusters. (C) Module-specific gene expression programs visualized on UMAPs. (D) KEGG pathway enrichment analyses of identified transcriptional modules. (E) Distribution of M1- and M2-associated transcriptional signatures across monocyte and macrophage populations. (F) Quantification of M1 and M2 polarization scores by cluster.

Abbreviations: GMSD=GM-CSF signaling deficient (i.e., *Csf2*<sup>-/-</sup>, *Csf2ra*<sup>-/-</sup> and *Csf2rb*<sup>-/-</sup>), WT=wild type.

N=4-8 mice/genotype), AM-alveolar macrophages, IM-Interstitial macrophages.
