## Supplemental tables for "GM-CSF Programs Alveolar Macrophages to Maintain Pulmonary Lymphocyte Homeostasis"

Supplementary Table 1. Antibody panel for Spectral flow cytometry

| Antibody | Vendor | Catalog number | Working dilution |
| --- | --- | --- | --- |
| Siglec F PE | BD Pharmingen™ | Cat552126 | 1: 150 |
| CD45 efluor 450 | eBioscience™ | REF 48-0451-82 | 1: 150 |
| CD11c BV480 | BD Horizon™ | Cat565627 | 1: 150 |
| CD11b PEcy7 | BD Pharmingen™ | Cat552850 | 1: 150 |
| B220 APC | Invitrogen<br>eBioscience™ | REF 17-0452-82 | 1: 150 |
| CD3 ALEXAfluor 700 | BD Pharmingen™ | Cat561388 | 1: 150 |
| Ly6G BUV395 | BD Horizon™ | Cat565964 | 1: 150 |
| Zombie NIR | BioLegend® | Cat423105 | 1: 500 |

Supplementary Table 2. Real-time quantitative PCR primers to quantify gene expression

| **TaqMan gene expression assay details** |  |  |  |  |
| --- | --- | --- | --- | --- |
| Gene symbol | Protein encoded | RefSeq | Amplicon length (bp) | Catalogue number |
| *Spi1* | PU.1 | NM_011355.1 | 91 | Mm00488142_m1 |
| *Pparg* | Ppar-γ | NM_011146.3 | 101 | Mm01184322_m1 |
| *Cebpβ* | C/EBPβ | NM_001287738.1 | 105 | Mm07294206_s1 |
| *Cebpα* | C/EBPα | NM_001287514.1 | 99 | Mm00514283_s1 |
| *NF-κb1* | NF-kappa-B p105 | NM_008689.2 | 70 | Mm00476361_m1 |
| *STAT1* | STAT1 | NM_001205313.1 | 75 | Mm01257286_m1 |
| *Bach2* | BACH2 | NM_001109661.1 | 81 | Mm00464379_m1 |
| *Lgals3* | Galectin-3 | NM_001145953.1 | 108 | Mm00802901_m1 |
| *Mrc1* | MMR1 | NM_008625.2 | 70 | Mm01329359_m1 |
| *Chil3* | Chil3 | NM_009892.2 | 90 | Mm00657889_mH |
| *Lpl* | LPL | NM_008509.2 | 61 | Mm00434764_m1 |
| *Car4* | Car4 | NM_007607.2 | 79 | Mm00483021_m1 |
| *Plet1* | PLET1 | NM_029639.2 | 78 | Mm01170995_m1 |
| *Slfn4* | Slfn4 | NM_146032 | 63 | Mm01298330_m1 |
| *Apoe* | apoE | NM_009696.3 | 64 | Mm01307192_m1 |
| *Gpnmb* | Gpnmb | NM_053110.4 | 66 | Mm01328587_m1 |
| *Cxcl9* | Cxcl9 | NM_008599.4 | 64 | Mm00434946_m1 |
| *Cxcl10* | Cxcl10 | NM_021274.2 | 59 | Mm00445235_m1 |
| *Cxcl16* | Cxcl16 | NM_023158.6 | 67 | Mm00469712_m1 |
| *Csf1r* | Csf1r | NM_001037859.2 | 69 | Mm01266652_m1 |
| *Csf1* | Csf1 | NM_001113529.1 | 70 | Mm00432686_m1 |
| *18S* | 18S rRNA | X03205.1 | 187 | 4310893E |

Supplemental Table 3

| Figure S2. KEGG Pathway Enrichment Analysis of Upregulated Genes Identified by Spatial Transcriptomic Analysis |  |  |  |
| --- | --- | --- | --- |
| Term | Overlap | Adjusted P-value | Genes |
| Antigen Receptor-Mediated Signaling Pathway (GO:0050851) | 56/134 | 2.67E-21 | IGHM;ITK;PIK3CD;CD3G;PTPN22;SLA2;CD3E;CD3D;IGHG3;IGHG1;TRBC2;TRBC1;ZC3H12A;NFKBIZ;BLNK;IGLC3;CD38;CTLA4;IGLC1;SYK;PRKCB;THEMIS;LAX1;LAT2;ZAP70;CD8A;LCK;BTK;LCP2;MS4A1;SKAP1;BLK;RNF31;STOML2;THY1;MALT1;CD79B;CD79A;FCHO1;IGKC;CD19;INPP5D;PLCG2;NCKAP1L;TRDC;CSK;FYN;SH2B2;RFTN1;NFATC2;PTPRC;THEMIS2;BCL2;CD28;CD247;NFKBID |
| Positive Regulation Of Cytokine Production (GO:0001819) | 85/320 | 6.85E-18 | CD86;ITK;CD40;TRAF3IP3;IL23R;CD80;SERPINE1;PIK3CD;PTPN22;CLEC9A;TNF;PIK3CG;IFIH1;PYCARD;PNP;DHX58;CASP1;CYP1B1;FFAR2;JAK2;IKBKE;CCR2;HAVCR2;GBP5;FCER1G;SYK;IL16;CYBA;TRAF2;TLR1;FGR;SLC7A5;IRF4;IL1B;TLR9;IRF7;TLR7;IRF5;LCP1;LTB;RAB7B;TMEM106A;TLR2;PTGER4;CD274;CSF1R;STOML2;HIF1A;LY9;AIF1;TANK;RASGRP1;MALT1;HSPD1;C3;G3BP1;PLCG2;HMOX1;SLAMF6;CD14;CCL19;IL12RB1;PIBF1;CD74;IL33;XBP1;NOS2;RFTN1;LAPTM5;EIF2AK2;ISG15;SULF1;POU2F2;SCIMP;BATF;CCDC88B;LACC1;CD4;CD40LG;PTPRC;POU2AF1;CD28;HPSE;MYD88;CRLF2 |
| B Cell Receptor Signaling Pathway (GO:0050853) | 28/46 | 1.52E-15 | BLK;IGHM;ITK;PIK3CD;CD79B;IGHG3;CD79A;IGHG1;IGKC;CD19;BLNK;IGLC3;PLCG2;CD38;CTLA4;IGLC1;NCKAP1L;SH2B2;SYK;PRKCB;RFTN1;NFATC2;LAT2;PTPRC;LCK;BCL2;BTK;MS4A1 |
| Defense Response To Virus (GO:0051607) | 54/189 | 2.13E-12 | RTP4;IFITM1;CD40;IFIT1;ZDHHC1;TNF;LYST;IFIT3;IFIT2;PYCARD;IFIH1;SENP7;TRIM7;DHX58;CASP1;DHX15;TRIM26;TRIM21;IKBKE;GBP5;MORC3;RSAD2;RIPK3;RNASE6;ACOD1;AIM2;OAS2;OAS3;DDIT4;IRF7;TLR9;NCBP3;TLR7;TLR2;NLRC5;DDX60;SAMHD1;TANK;USP18;EXOSC4;G3BP1;PMAIP1;GBP2;STAT1;STAT2;EIF2AK2;ISG15;PML;BST2;CXCL10;PTPRC;BCL2;MAP3K14;MYD88 |
| T Cell Activation (GO:0042110) | 38/111 | 4.43E-11 | CD86;ITK;CD80;PIK3CD;CD3G;PTPN22;NLRC3;SLA2;CD3E;LY9;RASGRP1;CD3D;PIK3CG;HSPD1;TRBC2;TRBC1;SLAMF7;SLAMF6;TRDC;CCL19;TNFRSF4;CD74;TNFSF14;RHOH;FOXP3;ZAP70;CD4;PTPRC;JAML;CD8A;IRF4;LCK;TREML2;CD7;CD28;LCP1;CD247;PTPN2 |
| T Cell Receptor Signaling Pathway (GO:0050852) | 34/95 | 1.79E-10 | RNF31;ITK;STOML2;PIK3CD;CD3G;PTPN22;THY1;CD3E;CD3D;MALT1;FCHO1;TRBC2;INPP5D;NFKBIZ;TRBC1;ZC3H12A;PLCG2;CTLA4;CSK;FYN;TRDC;RFTN1;THEMIS;ZAP70;PTPRC;THEMIS2;CD8A;LCK;BTK;CD28;LCP2;CD247;NFKBID;SKAP1 |
| Response To Cytokine (GO:0034097) | 39/125 | 4.04E-10 | CD274;CSF1R;IFITM1;CD40;CIITA;SP100;SHMT2;FLT3;IL23R;ADAR;TNFRSF11A;CXCL16;REL;IRAK2;UBD;CASP3;KYN;TIMP1;JAK2;TRIM21;IKBKE;JAK3;CD74;STAT1;EIF2AK2;ISG15;PML;NFKB2;BST2;SELP;CH25H;COL3A1;ALDH1A2;IL1B;BCL2;REL;HCLS1;XAF1;MYD88 |
| Defense Response To Symbiont (GO:0140546) | 43/148 | 4.04E-10 | RTP4;IFITM1;CD40;NLRC5;IFIT1;DDX60;SAMHD1;LYST;TANK;IFIT3;IFIT2;PYCARD;IFIH1;EXOSC4;G3BP1;CASP1;DHX15;PMAIP1;GBP2;IKBKE;GBP5;RSAD2;RIPK3;STAT1;STAT2;RNASE6;ACOD1;EIF2AK2;ISG15;BST2;PTPRC;AIM2;OAS2;OAS3;DDIT4;BCL2;IRF7;TLR9;NCBP3;TLR7;MAP3K14;MYD88;TLR2 |

Supplemental Table 3

| Figure S2. KEGG Pathway Enrichment Analysis of Upregulated Genes Identified by Spatial Transcriptomic Analysis |  |  |  |
| --- | --- | --- | --- |
| Term | Overlap | Adjusted P-value | Genes |
| Positive Regulation Of Lymphocyte Proliferation (GO:0050671) | 29/74 | 4.67E-10 | CD86;SASH3;CD274;CD40;CD80;IL23R;IRS2;TNFRSF13C;CD3E;TNFSF13B;PNP;CCL5;CD38;NCKAP1L;CCL19;JAK2;TNFRSF4;CD74;VCAM1;NFATC2;PDCD1LG2;CCDC88B;CD40LG;PTPRC;IL1B;GPR183;TLR9;BCL2;CD28 |
| Cytokine-Mediated Signaling Pathway (GO:0019221) | 59/257 | 1.52E-09 | CXCL9;SPI1;CSF3R;FLT3;IL23R;CXCL13;TNF;CXCL5;IL27RA;PYCARD;IL18RAP;TNFSF11;JAK2;JAK3;CCR2;IL13RA1;FCER1G;SYK;GPR35;TRAF2;IRAK4;TRAF1;AIM2;OAS2;IL1B;IRF5;IL9R;BIRC3;CSF1R;CSF2RB;TNFRSF11A;IL2RG;LRP8;SOCS1;CCL8;CCL7;IRAK2;CCL5;CCL4;STAT4;CCL3;IL21R;PTK2B;CCL19;IL12RB1;CD300LF;IL33;CCL22;CCL20;STAT1;NFKBIA;CXCL10;CD4;CXCL12;IL2RA;PTPN6;IL7R;MYD88;CRLF2 |
| Positive Regulation Of Response To External Stimulus (GO:0032103) | 43/155 | 1.64E-09 | PTGER4;RNF31;SERPINE1;LY96;IFI35;TNF;PLA2G7;AIF1;PYCARD;PLAU;CCL5;CASP4;NFKBIZ;CASP1;CCL3;EMILIN2;CCL19;SLC15A4;GPSM3;CCR2;GSDMD;GBP5;IL33;CD180;PDE2A;LY86;ACOD1;NKG7;CYBA;IL16;SCIMP;NFKBIA;AIM2;TNIP1;IL1B;NINJ1;TLR9;ADAM8;TLR7;LCP1;S100A9;S100A8;TLR2 |
| Cellular Response To Lipopolysaccharide (GO:0071222) | 37/124 | 4.40E-09 | CD86;CD274;CXCL9;CD80;GFI1;SERPINE1;LY96;TNFAIP3;PTPN22;CXCL13;TNF;CXCL5;MTDH;PYCARD;UXT;ZFP36;SBNO2;CASP7;CCL5;ZC3H12A;PLCG2;ANKRD1;CASP1;CCL3;CD14;CCR5;GBP2;XBP1;NFKBIL1;CD180;PDCD1LG2;CXCL10;TNIP1;IL1B;TNIP3;MYD88;NFKBIB |
| Regulation Of B Cell Proliferation (GO:0030888) | 21/44 | 4.65E-09 | SASH3;CD74;CD40;TNFRSF13B;CD300A;NFATC2;IRS2;TNFRSF13C;IKZF3;TNFSF13B;PTPRC;GPR183;BCL2;MZB1;TLR9;CD38;CTLA4;NCKAP1L;FCGR2B;TNFRSF4;CD22 |
| Response To Lipopolysaccharide (GO:0032496) | 42/159 | 1.28E-08 | CD86;CD274;CXCL9;CD80;GFI1;IL23R;SERPINE1;LY96;TNFAIP3;PTPN22;TNFRSF11A;CXCL13;CXCL5;PYCARD;UXT;ZFP36;SBNO2;CASP7;ZC3H12A;ANKRD1;CASP1;CD14;CCR5;JAK2;GBP2;XBP1;NOS2;NFKBIL1;CD180;PDCD1LG2;SELP;CXCL10;ADAM17;SLPI;TNIP1;IL1B;TNIP3;TRIB1;S100A9;S100A8;MYD88;NFKBIB |
| Positive Regulation Of Defense Response (GO:0031349) | 36/124 | 1.55E-08 | PTGER4;RNF31;SERPINE1;IFI35;TNF;PLA2G7;PYCARD;CCL5;CASP4;NFKBIZ;CASP1;CCL3;EMILIN2;SLC15A4;GPSM3;CCR2;GSDMD;GBP5;IL33;PDE2A;ACOD1;NKG7;CYBA;IL16;NFKBIA;AIM2;TNIP1;IL1B;NINJ1;TLR9;ADAM8;TLR7;LCP1;S100A9;S100A8;TLR2 |
| Response To Type II Interferon (GO:0034341) | 28/80 | 1.55E-08 | CD40;CIITA;IFITM1;SP100;IL23R;AIF1;CXCL16;CCL8;CCL7;KYNLU;CCL5;UBD;CCL4;CASP1;CCL3;CCL19;GBP2;IL12RB1;TRIM21;GBP4;GBP5;CD74;CCL22;CCL20;STAT1;BST2;IRF8;TLR2 |
| B Cell Activation (GO:0042113) | 30/92 | 2.32E-08 | CD86;CD40;FLT3;PIK3CD;RASGRP1;MALT1;HSPD1;FCRLA;CD79B;CD79A;BLNK;PLCG2;JAK3;VCAM1;ITGA4;PRKCB;PAX5;LAX1;LAT2;ZAP70;CD40LG;PTPRC;BANK1;GPR183;BTK;BCL2;TCF3;MS4A1;CD22;PTPN2 |
| Regulation Of B Cell Differentiation (GO:0045577) | 13/19 | 6.22E-08 | XBP1;SPI1;SYK;IKZF3;IL4I1;SFRP1;PPP2R3C;ID2;INPP5D;SLAMF8;TLR9;CD27;NCKAP1L |
| Inflammatory Response (GO:0006954) | 52/236 | 7.37E-08 | CD40;CIITA;CXCL9;NRROS;GPR68;TUSC2;ITGB2;PIK3CD;CXCL13;ITGAL;TNF;CXCL5;PIK3CG;C3AR1;BLNK;CCR5;TNFRSF4;CCR2;CD96;CYBA;TLR1;IL1B;ADAM8;S100A9;S100A8;TLR |

**Supplemental Table 3**

| Figure S2. KEGG Pathway Enrichment Analysis of Upregulated Genes Identified by Spatial Transcriptomic Analysis |  |  |  |
| --- | --- | --- | --- |
| Term | Overlap | Adjusted P-value | Genes |
|  |  |  | 2;PTGER4;CSF1R;PTGER2;AIF1;RELB;CCL8;CCL7;IRAK2;CCL5;CCL4;CCL3;CD14;CCL19;PLA2G2D;XCR1;CCL22;VCAM1;NOS2;CCL20;NFKB2;CXCL10;CD40LG;IL2RA;REL;ACKR1;FCGR2B |
| Regulation Of I-kappaB kinase/NF-kappaB Signaling (GO:0043122) | 51/232 | 1.08E-07 | CD40;TNFAIP3;ARRB2;TNF;PYCARD;ZC3H12A;NUP62;DHX15;CASP1;TRIM21;IKBKE;UNC5CL;PARP1;PRKCB;SHISA5;RHOH;TRAF2;IRAK4;TRAF1;IL1B;TLR9;S100A4;TRIM59;LTB;ALPK1;TMEM106A;BIRC3;RNF31;LPAR1;DDX21;NLRC3;CXXC5;TANK;MALT1;MTDH;UBD;TIFA;PLCG2;HMOX1;CCL19;ECT2;CD74;STAT1;NEK6;BST2;CD4;TNIP1;TNIP3;REL;MAP3K14;MYD88 |
| Negative Regulation Of Lymphocyte Activation (GO:0051250) | 14/23 | 1.08E-07 | LST1;LAPTM5;FOXJ1;ILDR2;TNFAIP3;PTPN22;LAX1;SAMSN1;IL4I1;GPNMB;TNFAIP8L2;FCGR2B;JAK3;PIBF1 |

Supplemental Table 3

| Figure S2. KEGG Pathway Enrichment Analysis of Downregulated Genes Identified by Spatial Transcriptomic Analysis |  |  |  |
| --- | --- | --- | --- |
| Term | Overlap | Adjusted P-value | Genes |
| Regulation Of Cell Migration (GO:0030334) | 81/434 | 6.92E-13 | ACVRL1;FLT1;SERPINE2;ULK4;LAMC2;ADARB1;PTPRK;LDB2;FGF1;SYNE2;CX3CL1;MYLK;AMOT;CDH5;DACH1;CDH1;DAG1;KDR;PHACTR1;RAC1;ARPIN;PDGFRA;EDN1;DUSP3;LIMCH1;SEMA6D;APLP2;HGF;F2R;CLDN5;F7;RAP2A;CDH11;NUMB;MYADM;DPEP1;CDH13;ITGA6;EPHA1;VCL;ENG;SMIM22;BEX4;CD151;SEMA3C;SEMA3B;PDGFB;SEMA3G;PDGFA;PIK3R3;PRR5L;SEMA3E;SEMA3F;PPM1F;NKD1;PARD6B;PODXL;PDPN;NKX2-1;PLXNA1;SRGAP1;PLXNA4;SASH1;TCAF1;ACE;FZD4;CAV1;LAMB1;PLET1;TBX5;MAPK15;PTK2;VEGFA;SMAD7;TJP1;SH3RF2;KIF26A;CNTN1;AMOTL2;FERMT2;BCAR1 |
| Positive Regulation Of Cell Migration (GO:0030335) | 54/272 | 3.10E-09 | BMPR2;FLT1;LAMC2;FGF1;SYNE2;MYLK;CDH5;SYDE1;HYAL1;KDR;PDGFRA;EDN1;HGF;SEMA6D;F2R;GAB1;F7;MYADM;NUMB;CDH13;ITGA6;EPHA1;SMIM22;CD151;SEMA3C;SEMA3A;SEMA3B;PDGFB;SEMA3G;PDGFA;PIK3R3;SEMA3E;SEMA3F;PPM1F;VTN;RAB25;PODXL;PDPN;TGFB2;DMTN;SMURF2;FZD4;CAV1;FN1;DAB2IP;LAMB1;PLET1;PTK2;VEGFA;TJP1;SH3RF2;CNTN1;FERMT2;BCAR1 |
| Positive Regulation Of Cell Motility (GO:2000147) | 45/221 | 6.06E-08 | CD151;FLT1;SEMA3C;SEMA3B;PDGFB;SEMA3G;PDGFA;PIK3R3;LAMC2;SEMA3E;SEMA3F;FGF1;NTN1;SYNE2;PPM1F;MYLK;CDH5;PODXL;SPOCK2;PDPN;KDR;PDGFRA;EDN1;CAVIN1;FZD4;HGF;SEMA6D;CAV1;F2R;LAMB1;PLET1;PTK2;VEGFA;TJP1;F7;SH3RF2;MYADM;CNTN1;NUMB;CDH13;ITGA6;EPHA1;FERMT2;BCAR1;SMIM22 |
| Cell-Cell Junction Organization (GO:0045216) | 23/68 | 6.62E-08 | PRKCI;TGFB2;CXADR;CTNND1;F2R;GJC1;TJP1;CDH5;CLDN5;PARD6B;OCLN;MARVELD2;CDH1;GJA4;CDH11;CDHR3;MYADM;PKP2;CDH13;PKP4;TLN2;LIMS2;TJP2 |
| Cell-Cell Junction Assembly (GO:0007043) | 23/71 | 1.43E-07 | PATJ;RAMP2;JUP;CTNND1;SMAD7;GJC1;TJP1;CDH5;CLDN5;PARD6B;GJB2;CDH1;GJA5;GJA4;CDH11;CDHR3;PKP2;CTNNA1;CDH13;CD9;PKP4;TLN2;VCL |
| Heart Development (GO:0007507) | 35/171 | 3.60E-06 | ACVRL1;POPDC3;RAMP2;BMPR2;PDGFB;CCM2L;GATA5;PKD2;PDLIM1;PDLIM3;PDLIM2;SOX17;ERBB3;SOX18;FLRT3;GJA5;GPC3;PLCE1;UTY;TGFB2;CXADR;FOXF1;FN1;MICAL2;TBX5;PTK2;TBX3;BMP5;APLN;TBX2;VEGFA;BMP4;TGFB3;PKP2;TEK |
| Cilium Movement (GO:0003341) | 19/59 | 4.27E-06 | CFAP61;SPAG17;CCDC39;DNAH11;RSPH4A;SPEF2;CFAP73;SPEF1;DNAH5;DNAH9;RSPH9;NME5;CFAP206;TTC29;CFAP100;TTLL3;TEKT1;HYDIN;CFAP54 |
| Outflow Tract Septum Morphogenesis (GO:0003148) | 12/24 | 6.34E-06 | ROBO2;BMP4;TGFB2;BMPR2;SEMA3C;GATA6;PARVA;LRP2;SMAD6;FGFR2;TBX2;ENG |
| Transmembrane Receptor Protein Serine/Threonine Kinase Signaling Pathway (GO:0007178) | 29/130 | 6.34E-06 | ACVRL1;BMPR2;HPGD;TGFB1I1;SCX;PTPRK;TMEM100;CDH5;MAPK3;TGFB2;SMURF2;APLP2;SMAD9;FOS;SMAD6;ACVR2B;BMP6;PTK2;BMP5;SMAD7;RGMA;BMP4;TGFB3;CLDN5;BMP3;TWSG1;ID1;FERMT2;ENG |
| Cell Junction Assembly (GO:0034329) | 24/95 | 7.61E-06 | GPM6A;BDNF;CTNND1;FN1;GJC1;TJP1;CDH5;CLDN5;PARD6B;CDH1;FLRT3;SPOCK2;GJA4;CDH11;CDHR3;PKP2;CDH13;NRCAM;ITGA6;PKP4;TLN2;SHANK3;SHANK2;ADGRL3 |
| Regulation Of Angiogenesis (GO:0045765) | 38/205 | 7.75E-06 | ECM1;RAMP2;FLT1;NPR1;GATA6;PTPRM;SEMA3E;ISM1;GATA2;FGF1;TSPAN12;AQP1;AMOT;CDH5;BMPER;HYAL1;ALOX5;KDR;SASH1;TGFB2;PTGIS;JUP;ANXA3;CEMIP2;TNFSF1 |

**Supplemental Table 3**

| Figure S2. KEGG Pathway Enrichment Analysis of Downregulated Genes Identified by Spatial Transcriptomic Analysis |  |  |  |
| --- | --- | --- | --- |
| Term | Overlap | Adjusted P-value | Genes |
|  |  |  | 2;GAB1;DAB2IP;EMP2;VEGFD;KLF4;VEGFA;CLDN5;COL4A3;CNTN1;PRKD1;TEK;EPHA1;RAPGEF3 |
| Axon Guidance (GO:0007411) | 31/149 | 8.65E-06 | ROBO2;NOTCH3;SEMA3C;SEMA3A;SEMA3B;SEMA3G;PTPRM;SEMA3E;SEMA3F;NTN1;ARRHGAP35;EFNB2;EFNB1;ALCAM;CHN1;DAG1;PLXNA1;NRCAM;RAC1;ARHGEF40;NEO1;PLXNA4;BDNF;SEMA6D;PTK2;VEGFA;EFNA1;GAP43;GDNF;NFIB;CNTN1 |
| Artery Morphogenesis (GO:0048844) | 13/30 | 8.65E-06 | TGFB2;BMPR2;HPGD;FOXF1;LRP2;PKD2;SMAD7;TBX2;VEGFA;BMP4;GJA5;HES1;ENG |
| Positive Regulation Of Angiogenesis (GO:0045766) | 27/119 | 8.65E-06 | ECM1;RAMP2;FLT1;GATA6;FGF1;GATA2;AQP1;CDH5;BMPEP;HYAL1;KDR;SASH1;PTGIS;JUP;ANXA3;TNFSF12;GAB1;EMP2;VEGFD;KLF4;VEGFA;TJP1;CNTN1;PRKD1;TEK;EPHA1;RAPGEF3 |
| Regulation Of Cell Population Proliferation (GO:0042127) | 94/766 | 1.05E-05 | ACVRL1;BMPR2;IRS1;TGFB1I1;RTKN2;HILPDA;SCX;LAMC2;ADARB1;PTPRK;FGF1;PKD2;CX3CL1;RERG;KDR;ADAMTS8;SOX7;TNS2;PDGFRA;KLF11;EDN1;EDN3;TSC22D1;IGFBP3;F2R;ITGA1;P3H2;EMP2;LIFR;CGRRF1;TENT5B;BTC;ACE2;CLDN5;AZGP1;DDAH1;COL4A3;CDH13;ATF5;EPHA1;PLPP1;SHANK2;ENG;SMIM22;KANK2;BEX4;YAP1;CTBP2;CUL3;NPR3;PDGFB;LRP5;PDGFA;TOB1;EFNB2;DLL4;NUAK1;RAB25;ERBB3;NACC2;PDPN;HES1;IGFBP6;WNT2;JAG2;WWTR1;CDKN2B;TGFB2;CADM4;JUP;WNT3A;FN1;DAB2IP;IGF2;VEGFD;TBX5;SMAD6;PTPN14;MAPK15;TBX3;PTK2;BMP6;VEGFA;BMP5;ACER2;TJP1;BMP4;GDNF;NOX4;FGFR4;FGFR3;CRLF1;HSPA1B;FGFR2 |
| Blood Vessel Morphogenesis (GO:0048514) | 17/53 | 1.21E-05 | EGFL7;RAMP2;FLT1;CAV1;SLC1A1;LRP5;RASIP1;SMAD7;VEGFA;EFNB2;SOX17;SOX18;GJA5;ID1;KDR;HES1;ENG |
| Regulation Of Endothelial Cell Proliferation (GO:0001936) | 23/93 | 1.29E-05 | ECM1;EGFL7;BMPR2;ANGPT1;CAV2;CAV1;TIE1;TNFSF12;SORD;PDGFB;PTPRM;NRARP;VEGFD;VEGFA;BMP4;NR4A1;ALOX5;KDR;CNTN1;CDH13;ANG;PRKD1;TEK |
| Branching Morphogenesis Of An Epithelial Tube (GO:0048754) | 15/42 | 1.29E-05 | TIE1;LRP5;TCF21;NRARP;SEMA3E;PKD2;NPNT;RASIP1;VEGFA;DLL4;BMP4;GDNF;KDR;NKX2-1;ENG |
| Cellular Response To Growth Factor Stimulus (GO:0071363) | 31/155 | 1.44E-05 | ACVRL1;RAMP2;BMPR2;FLT1;STMN2;PDGFB;NDNF;LRP2;TMEM100;DLL4;HYAL1;KDR;WNT2;DUSP3;APLP2;HGF;DAB2IP;SMAD9;WNT7A;KLF4;ACVR2B;BMP6;SMAD7;MTSS2;VEGFA;BMP4;NR4A1;PDE3A;PRKD1;MAPT;BCAR1 |
| Positive Regulation Of Cell Junction Assembly (GO:1901890) | 18/61 | 1.69E-05 | CLSTN2;BDNF;CAV1;IRX3;CLSTN1;WNT7A;TBX5;PPM1F;VEGFA;PTPRD;ACE2;CLDN5;SRPX2;NPHP1;KDR;RAC1;TEK;FERMT2 |
| Regulation Of Endothelial Cell Migration (GO:0010594) | 22/88 | 1.70E-05 | ACVRL1;EDN1;BMPR2;ANGPT1;ANXA3;DAB2IP;PTPRM;EMP2;FGF1;PTK2;VEGFA;DLL4;BMP4;BMPEP;DNAJA4;KDR;RAC1;PRKD1;TEK;PLPP3;SASH1;BCAR1 |

**Supplemental Table 3**

| **Figure S4A. KEGG Pathway Enrichment Analysis of Upregulated Genes Identified by Single Nuclear RNA-Sequencing analysis in Lymphoid Cells** |  |  |  |  |
| --- | --- | --- | --- | --- |
| **Term** | **Short term** | **Overlap** | **Adjusted P-value** | **Genes** |
| Stem Cell Differentiation (GO:0048863) | Stem cell differentiation | 5/47 | 0.003735 | MEF2C;PTPRC;FBXL17;MLLT3;JARID2 |
| Protein Modification Process (GO:0036211) | Protein modification | 12/545 | 0.006325 | CAMK2D;PTPRC;PRKCB;PRKCE;STK17B;AKT3;ST8SIA4;PRKCA;BIRC6;STK4;SSH2;JAK1 |
| Regulation of Sodium Ion Transmembrane Transport (GO:1902305) | Regulation of Na<sup>2+</sup> transport | 3/20 | 0.026463 | CAMK2D;PRKCE;UTRN |
| Antigen Receptor-Mediated Signaling Pathway (GO:0050851) | Antigen receptor signaling | 5/114 | 0.039202 | DENND1B;MEF2C;PTPRC;PRKCB;INPP5D |
| Regulation of Cell Junction Assembly (GO:1901888) | Regulation of cell junction assembly | 3/25 | 0.039202 | MEF2C;RAP1A;PRKCA |
| Negative Regulation of Granulocyte Differentiation (GO:0030853) | Inhibition of granulocyte differentiation | 2/6 | 0.046215 | INPP5D;RUNX1 |
| Peptidyl-Serine Modification (GO:0018209) | Peptidyl serine modification | 4/75 | 0.046215 | CAMK2D;PRKCE;PRKCA;STK4 |
| Mitotic Nuclear Membrane Disassembly (GO:0007077) | Mitotic progression | 2/7 | 0.046215 | PRKCB;PRKCA |
| Peptidyl-Serine Phosphorylation (GO:0018105) | Peptidyl serine phosphorylation | 4/70 | 0.046215 | CAMK2D;PRKCE;PRKCA;STK4 |
| Negative Regulation of Sodium Ion Transport (GO:0010766) | Inhibition of Na<sup>2+</sup> transport | 2/7 | 0.046215 | CAMK2D;PRKCE |
| B Cell Receptor Signaling Pathway (GO:0050853) | BCR signaling | 3/33 | 0.046469 | MEF2C;PTPRC;PRKCB |
| Phosphatidylinositol Biosynthetic Process (GO:0006661) | Biosynthesis of phosphatidyl inositol | 4/91 | 0.063743 | ITPKB;INPP4B;INPP5D;PIP4K2A |
| B Cell Activation (GO:0042113) | B cell activation | 4/96 | 0.06663 | MEF2C;PTPRC;PRKCB;BANK1 |

**Supplemental Table 3**

| **Figure S4B. KEGG Pathway Enrichment Analysis of Upregulated Genes Identified by Single Nuclear RNA-Sequencing analysis in Epithelial Cells** |  |  |  |  |
| --- | --- | --- | --- | --- |
| **Term** | **Short term** | **Overlap** | **Adjusted P-value** | **Genes** |
| Positive Regulation of DNA-templated Transcription (GO:0045893) | Transcriptional activation | 54/1274 | 1.07E-04 | APP;MAML2;CHD6;RORA;NR3C1;RPS6KA3;PPP3CA;MECOM;ZMIZ1;TEAD1;MAP3K5;MEF2A;NCOA1;BCAS3;NCOA2;PPP1R12A;TCF12;ATRX;DYRK1A;LMO7;ARID1B;ETV5;RUNX1;SFPQ;MDFIC;IRF2;WAC;PHIP;PRKD1;MAML3;PRKN;NFAT5;KMT2C;FOXO3;MLLT3;STRN3;PDZD2;PRDM16;APBB2;TBL1X;RREB1;BRD4;WWOX;CREBBP;ZFHX3;TCF7L1;NFKB1;MLLT10;MYO1E;NFIA;KANSL1;CDK13;SSBP2;LPIN2 |
| Positive Regulation of Transcription by RNA Polymerase II (GO:0045944) | Polymerase-II mediated transcriptional activation | 45/983 | 1.07E-04 | PRKN;NFAT5;APP;MAML2;KMT2C;CHD6;RORA;FOXO3;NR3C1;STRN3;RPS6KA3;PPP3CA;PDZD2;ZMIZ1;APBB2;TBL1X;RREB1;TEAD1;BRD4;MEF2A;NCOA1;WWOX;BCAS3;NCOA2;CREBBP;ZFHX3;TCF7L1;PPP1R12A;TCF12;ATRX;LMO7;NFKB1;ETV5;RUNX1;MLLT10;MYO1E;SFPQ;NFIA;IRF2;PHIP;PRKD1;MAML3;CDK13;SSBP2;LPIN2 |
| Regulation of Small GTPase Mediated Signal Transduction (GO:0051056) | Regulation of small GTPase mediated signaling | 15/157 | 2.65E-04 | TRIO;ARHGEF12;DOCK9;ARHGEF28;DENND1A;DOCK7;ARHGAP5;MYO9A;ARHGAP4;AKAP13;ARHGAP31;WNK1;DLC1;SRGAP2;CGNL1 |
| Negative Regulation of DNA-templated Transcription (GO:0045892) | Negative regulation of transcription | 44/1006 | 2.65E-04 | PRKN;APP;WWC1;WWC2;ZBTB20;OTUD7B;PTPRK;FOXO3;HDAC8;NR3C1;PHF21A;STRN3;ATXN1;MECOM;CUX1;PRDM16;SCMH1;MBIP;APBB2;TBL1X;RREB1;MEF2A;MAGI1;CREBBP;ZFHX3;TCF7L1;RBF2;SMURF2;USP9X;WWP1;ARID5B;FOXN3;FOX2;NFKB1;ETV5;PIAS1;FOXP1;RUNX1;DLG1;SFPQ;NR6A1;MDFIC;IRF2;LCOR |
| Actin Filament Organization (GO:0007015) | Actin cytoskeleton organization | 14/158 | 7.29E-04 | CARMIL1;SVIL;NEBL;TPM1;COBL;SHROOM3;PPP1R9A;MYO1D;MYO1E;DLG1;MYO1B;MYO5B;PLS3;CGNL1 |
| Negative Regulation of Transcription by RNA Polymerase II (GO:0000122) | Regulation of transcription by Pol II | 34/732 | 7.29E-04 | PRKN;APP;WWC1;WWC2;ZBTB20;OTUD7B;FOXO3;HDAC8;NR3C1;PHF21A;STRN3; |

**Supplemental Table 3**

| **Figure S4B. KEGG Pathway Enrichment Analysis of Upregulated Genes Identified by Single Nuclear RNA-Sequencing analysis in Epithelial Cells** |  |  |  |  |
| --- | --- | --- | --- | --- |
| **Term** | **Short term** | **Overlap** | **Adjusted P-value** | **Genes** |
|  |  |  |  | ATXN1;CUX1;PRDM16;MBIP;APBB2;TBL1X;RREB1;MEF2A;CREBBP;ZFXH3;TCF7L1;SMURF2;USP9X;ARID5B;NFKB1;ETV5;PIAS1;RUNX1;DLG1;SFPQ;NR6A1;IRF2;LCOR |
| Regulation of Cell Migration (GO:0030334) | Regulation of cell migration | 26/479 | 7.29E-04 | ITGB1;MACF1;GSK3B;RORA;SEMA3E;PTPRJ;LDLRAD4;PTPRK;FOXO3;PPP3CA;MGAT5;PLXNA2;DAG1;CTSH;SRGAP2;PLXNA4;SPAG9;CARMIL1;LIMCH1;TPM1;PTK2;TJP1;MYO1E;AFDN;DLC1;NF1 |
| Neuron Development (GO:0048666) | Neuronal development | 14/159 | 7.29E-04 | APP;GSK3B;DOCK7;PBX3;NEDD4L;PTPRM;VPS13B;PTPRK;PPP1R9A;RUNX1;PTPRG;VAPA;WNK1;ATXN10 |
| Negative Regulation of Cell Motility (GO:2000146) | Negative regulation of cell motility | 13/139 | 7.29E-04 | GSK3B;LIMCH1;TPM1;PTPRJ;LDLRAD4;PTPRK;FOXO3;AFDN;PDZD2;DLC1;DAG1;NF1;SRGAP2 |
| Negative Regulation of Cell Migration (GO:0030336) | Negative regulation of cell migration | 15/184 | 7.29E-04 | GSK3B;LIMCH1;TPM1;PTPRM;PTPRJ;LDLRAD4;PTPRK;FOXO3;PTPRG;AFDN;DLC1;DAG1;NF1;SRGAP2;PHLDB2 |
| Regulation of Catabolic Process (GO:0009894) | Regulation of catabolic process | 10/81 | 7.29E-04 | ITGB1;PRKN;UVRAG;DRAM1;DAPK1;FAF1;LAMP3;DAPK2;WAC;ITPR1 |
| Plasma Membrane Bounded Cell Projection Organization (GO:0120036) | Organization of cell projection | 13/141 | 7.29E-04 | APP;CARMIL1;GSK3B;TPM1;DOCK7;NEDD4L;PTPRM;VPS13B;PTPRK;PPP1R9A;PTPRG;VAPA;ATXN10 |
| Negative Regulation of Intracellular Signal Transduction (GO:1902532) | Inhibition of signal transduction | 19/297 | 0.001098 | GSK3B;WWC1;WWC2;HUWE1;ITPR1;OTUD7B;PTPRJ;RORA;LDLRAD4;UBR1;DUSP16;STRN3;PDZD2;WNK1;DAG1;MAP3K20;NF1;SLC39A8;CGNL1 |
| Regulation of Intracellular Signal Transduction (GO:1902531) | Regulation of signal transduction | 19/302 | 0.001273 | TRIO;ARHGEF12;SGMS1;DAPK2;WWC1;ZFAND6;ARHGEF28;WWC2;ARHGAP5;MYO9A;MOB3B;ARHGAP44;AKAP13;ARHGAP31;DLG1;TAOK3;DLC1;PIP5K1B;SRGAP2 |
| Negative Regulation of Protein Localization to Nucleus (GO:1900181) | Inhibition of nuclear import | 6/26 | 0.001273 | APP;GSK3B;MDFIC;CTNNA1;OTUD7B;CLDN18 |

**Supplemental Table 3**

| **Figure S4B. KEGG Pathway Enrichment Analysis of Upregulated Genes Identified by Single Nuclear RNA-Sequencing analysis in Epithelial Cells** |  |  |  |  |
| --- | --- | --- | --- | --- |
| **Term** | **Short term** | **Overlap** | **Adjusted P-value** | **Genes** |
| Neuron Projection Development (GO:0031175) | Neuron Projection Development | 15/206 | 0.001846 | APP;GSK3B;TRIO;PPP1R12A;DOCK7;NEDD4L;PTPRM;VPS13B;PTPRK;PPP1R9A;PTPRG;TAOK3;VAPA;ATXN10;SRGAP2 |
| Regulation of Canonical Wnt Signaling Pathway (GO:0060828) | Regulation of Wnt signaling | 15/216 | 0.003041 | PRKN;APP;GSK3B;SMURF2;USP34;MLLT3;NFKB1;PDZD2;ZNRF3;WNK1;GNAQ;TBL1X;CDK14;WLS;RBMS3 |
| Regulation of Cell-Matrix Adhesion (GO:0001952) | Regulation of cell matrix adhesion | 8/65 | 0.004509 | GSK3B;MACF1;LIMCH1;NF1;CASK;SEMA3E;UTRN;PTK2 |
| Supramolecular Fiber Organization (GO:0097435) | Supramolecular fiber organization | 19/339 | 0.004597 | PRKN;APP;CARMIL1;USP9X;TPM1;SHROOM3;PPP1R9A;THSD4;MYO1D;MYO1E;DLG1;MYO1B;PDZD2;BMP1;MYO5B;NF1;B2M;NCKAP5;CGNL1 |
| Positive Regulation of Cell Migration (GO:0030335) | Activation of cell migration | 19/344 | 0.005226 | ITGB1;SPAG9;CARMIL1;NRP1;BCAS3;SMURF2;ANXA3;RORA;SEMA3E;PTK2;FOXP1;TJP1;PPP3CA;MYO1E;MGAT5;CTSH;PRKD1;RREB1;DOCK1 |
| Establishment or Maintenance of Epithelial Cell Apical/Basal Polarity (GO:0045197) | Maintenance of Epithelial cell polarity | 6/36 | 0.005899 | PATJ;DLG1;MTCL1;PARD3;MYO9A;PARD3B |
| Neuron Projection Morphogenesis (GO:0048812) | Neuron Projection Development | 12/157 | 0.005899 | MEF2A;APP;FARP1;TRIO;ALCAM;PPP1R12A;TAOK3;PARD3;DOCK7;ANK3;IQGAP1;SRGAP2 |
| Regulation of Focal Adhesion Assembly (GO:0051893) | Regulation Focal adhesion assembly | 7/53 | 0.006309 | NRP1;MACF1;LIMCH1;DLC1;PTPRJ;PHLDB2;PTK2 |
| Regulation of Stress Fiber Assembly (GO:0051492) | Regulation of stress fiber assembly | 8/75 | 0.007898 | TJP1;NRP1;CARMIL1;LIMCH1;DLC1;TPM1;PHLDB2;CGNL1 |
| Positive Regulation of Hydrolase Activity (GO:0051345) | Activation of hydrolase activity | 9/98 | 0.008788 | ITGB1;PDZD2;DOCK9;NF1;SNX13;CTSH;PKP4;SRGAP2;RAPGEF6 |
| Regulation of Autophagy (GO:0010506) | Regulation of Autophagy | 14/230 | 0.010901 | PRKN;UVRAG;GSK3B;DAPK1;DAPK2;ITPR1;AMBRA1;FOXO3;DRAM1;WNK1;LAMP3;EXOC4;WAC;PRKD1 |

**Supplemental Table 3**

| **Figure S4C. KEGG Pathway Enrichment Analysis of Upregulated Genes Identified by Single Nuclear RNA-Sequencing analysis in Endothelial Cells** |  |  |  |  |
| --- | --- | --- | --- | --- |
| **Term** | **Short term** | **Overlap** | **Adjusted P-value** | **Genes** |
| Regulation of Cell Migration (GO:0030334) | Regulation of cell migration | 24/479 | 3.44E-08 | GSK3B;NOTCH1;ACE;MCTP1;SEMA3C;ROCK1;ROCK2;ANKRD44;LDB2;PTPRK;PTK2;SYNE2;IGF1R;TJP1;AFDN;DACH1;PDGFD;KIT;PLXNA2;NF1;NUMB;PHACTR1;FERMT2;SASH1 |
| Positive Regulation of Cell Migration (GO:0030335) | Activation of cell migration | 20/344 | 6.58E-08 | NRP1;SEMA6A;NOTCH1;BMPR2;SEMA3C;ROCK2;PRKCE;PTK2;SYNE2;IGF1R;TJP1;PDGFD;KIT;NUMB;RAPGEF2;TEK;RIN2;DOCK1;FERMT2;SASH1 |
| Regulation of Focal Adhesion Assembly (GO:0051893) | Regulation Focal adhesion assembly | 9/53 | 7.16E-07 | NRP1;ROCK1;ROCK2;LDB2;TEK;PHLDB2;PTK2;FERMT2;LIMS1 |
| Regulation of Epithelial Cell Migration (GO:0010632) | Regulation of epithelial cell migration | 8/43 | 2.11E-06 | BMPR2;ANKRD44;PRKCE;MCC;DOCK1;SASH1;PTK2;PTPRG |
| Positive Regulation of Cell-Matrix Adhesion (GO:0001954) | Activation of cell matrix adhesion | 8/45 | 2.47E-06 | GSK3B;NRP1;ROCK1;CD36;TEK;RIN2;FERMT2;LIMS1 |
| Regulation of Stress Fiber Assembly (GO:0051492) | Regulation of stress fiber assembly | 8/75 | 1.15E-04 | TJP1;NRP1;ROCK1;ROCK2;CD47;PHLDB2;WASF2;FERMT2 |
| Cell Surface Receptor Protein Tyrosine Kinase Signaling Pathway (GO:0007169) | Tyrosine kinase signaling pathway | 14/296 | 1.54E-04 | NRP1;GSK3B;CAV2;KALRN;PTK2;IGF1R;PTPRG;SIPA1L1;COL4A2;PDGFD;KIT;RAPGEF2;TEK;PLCB1 |
| Mesenchymal Cell Differentiation (GO:0048762) | Mesenchymal differentiation | 7/58 | 1.73E-04 | GSK3B;NOTCH1;ROCK1;SEMA3C;ROCK2;FBXL17;LIMS1 |
| Regulation of Cellular Component Biogenesis (GO:0044087) | Cellular component biogenesis regulation | 6/37 | 1.73E-04 | TJP1;NOTCH1;ROCK1;ROCK2;RAPGEF2;CD36 |
| Intracellular Signaling Cassette (GO:0141124) | Intracellular signaling modules | 15/354 | 1.78E-04 | MEF2A;VAV3;CREBBP;JUN;MCTP1;GAB2;NCALD;IGF1R;ARHGAP31;ZMIZ1;NF1;RAPGEF2;CD36;RIN2;DGKH |
| Small GTPase-mediated Signal Transduction (GO:0007264) | Small GTPase signaling | 11/188 | 2.01E-04 | RB1;VAV3;ARHGAP31;ROCK1;ROCK2;ELMO1;NF1;ARHGEF3;ARHGAP29;RAPGEF2;RIN2 |
| Regulation of Endothelial Cell Migration (GO:0010594) | Regulation of endothelial cell migration | 8/88 | 2.01E-04 | NRP1;ROCK2;NF1;PTPRM;TEK;RIN2;SASH1;PTK2 |
| Positive Regulation of Cell-Substrate Junction Organization (GO:0150117) | Organization of cell substrate junction | 5/23 | 2.18E-04 | NRP1;ROCK1;TEK;FERMT2;LIMS1 |

**Supplemental Table 3**

| **Figure S4C. KEGG Pathway Enrichment Analysis of Upregulated Genes Identified by Single Nuclear RNA-Sequencing analysis in Endothelial Cells** |  |  |  |  |
| --- | --- | --- | --- | --- |
| **Term** | **Short term** | **Overlap** | **Adjusted P-value** | **Genes** |
| Regulation of Cell-Matrix Adhesion (GO:0001952) | ECM regulation | 7/65 | 2.36E-04 | GSK3B;ROCK1;ROCK2;NF1;LDB2;CD36;PTK2 |
| Regulation of Cell Junction Assembly (GO:1901888) | Cell junction regulation | 5/25 | 2.79E-04 | TJP1;SETD5;ROCK1;ROCK2;RAPGEF2 |
| Cellular Response to Transforming Growth Factor Beta Stimulus (GO:0071560) | TGF-beta response | 8/98 | 3.12E-04 | SKI;JUN;ZMIZ1;PTPRK;NR3C1;PTK2;FERMT2;LIMS1 |
| Regulation of GTPase Activity (GO:0043087) | Regulation of GTPase activity | 9/134 | 3.69E-04 | VAV3;FGD5;SIPA1L1;CAV2;NF1;RAPGEF2;PKP4;PTK2;LIMS1 |
| Epithelial to Mesenchymal Transition (GO:0001837) | Epithelial-Mesenchymal transition | 6/51 | 5.53E-04 | GSK3B;NOTCH1;ROCK1;ROCK2;TBX3;LIMS1 |
| Positive Regulation of MAPK Cascade (GO:0043410) | Activation of MAPK pathway | 14/362 | 5.55E-04 | NRP1;NOTCH1;ROCK1;CAV2;ROCK2;IGF1R;PDGFD;KIT;RAPGEF2;CD36;TEK;PLCB1;FERMT2;SASH1 |
| Regulation of Cell Population Proliferation (GO:0042127) | Regulation of cell proliferation | 21/769 | 6.85E-04 | JUN;RBFOX2;NOTCH1;BMPR2;ITGA1;PTPRK;GAB2;SMARCA2;PTK2;TBX3;IGF1R;SKI;TJP1;PDGFD;ACER3;KIT;RAPGEF2;IGFBP7;BIRC6;CD47;LIMS1 |
| Integrin-Mediated Signaling Pathway (GO:0007229) | Integrin signaling | 7/86 | 9.14E-04 | NRP1;DST;ITGA1;CD47;DOCK1;PTK2;FERMT2 |
| Positive Regulation of Cell Motility (GO:2000147) | Activation of cell motility | 11/249 | 0.001288 | TJP1;NOTCH1;SEMA3C;ROCK2;PDGFD;KIT;NUMB;PTK2;FERMT2;SYNE2;IGF1R |
| Positive Regulation of Cell Differentiation (GO:0045597) | Activation of endothelial cell differentiation | 12/297 | 0.001288 | GSK3B;BMPR2;ZEB1;ZBTB16;RAPGEF2;CD36;PLCB1;RIN2;SMARCA2;ARID1B;PTK2;FERMT2 |
| Regulation of Signal Transduction (GO:0009966) | Regulation of signal transduction | 10/211 | 0.001475 | NRP1;GIT2;CREBBP;FBXL17;IGFBP7;RGS12;CD36;CD47;PLCB1;NCALD |
| Transforming Growth Factor Beta Receptor Signaling Pathway (GO:0007179) | TGF-beta receptor pathway | 6/76 | 0.003079 | SKI;JUN;ZMIZ1;PTPRK;PTK2;FERMT2 |

**Supplemental Table 3**

| **Figure S4D. KEGG Pathway Enrichment Analysis of Upregulated Genes Identified by Single Nuclear RNA-Sequencing analysis in Mesenchymal Cells** |  |  |  |  |
| --- | --- | --- | --- | --- |
| **Term** | **Short term** | **Overlap** | **Adjusted P-value** | **Genes** |
| Regulation of Focal Adhesion Assembly (GO:0051893) | Regulation of focal adhesion | 9/53 | 0.00001 | NRP1;MACF1;ROCK1;LIMCH1;PEAK1;DLC1; ARHGAP6;PHLDB2;LIMS1 |
| Outflow Tract Septum Morphogenesis (GO:0003148) | Cardiac outflow septation | 6/24 | 0.00017 | ROBO2;NRP1;BMPR2;PARVA;ZFPM2;TGFB2 |
| Membrane Depolarization During Cardiac Muscle Cell Action Potential (GO:0086012) | Cardiac membrane depolarization | 5/16 | 0.00022 | CACNB2;CACNA2D1;CACNA1D;CACNA1C;SLC8A1 |
| Cell-Substrate Junction Assembly (GO:0007044) | Cell-Substrate Junction Assembly | 6/29 | 0.00022 | DST;PEAK1;DLC1;CDH11;PTPRK;ARHGAP6 |
| Positive Regulation of Cell Migration (GO:0030335) | Activation of cell migration | 16/344 | 0.00045 | ITGB1;NRP1;PDGFRA;BCAS3;SPARC;BMPR2; SMURF2;ANGPT1;PRKCA;RORA;LAMB1;MYLK;IGF1R;FOXP1;COL1A1;NIPBL |
| Regulation of Cell Migration (GO:0030334) | Regulation of cell migration | 19/479 | 0.00045 | ITGB1;MACF1;PDGFRA;ROCK1;LIMCH1;ANKRD44;MAGI2;PRKCA;RORA;LAMB1;LDLRAD4;PTPRK;MYLK;IGF1R;COL1A1;DLC1;CDH11; PHACTR1;SLIT2 |
| Cell-Matrix Adhesion (GO:0007160) | Cell matrix adhesion | 9/107 | 0.00060 | ITGB1;COL3A1;PEAK1;DLC1;CDH11;ITGA8;PTPRK;NPNT;ARHGAP6 |
| Heart Development (GO:0007507) | Cardiac development | 11/171 | 0.00060 | MEF2A;AKAP13;NIPBL;COL3A1;BMPR2;HEG1;DLC1;CACNA1C;ZFPM2;FBN1;TGFB2 |
| Focal Adhesion Assembly (GO:0048041) | Focal adhesion assembly | 5/22 | 0.00060 | PEAK1;DLC1;CDH11;PTPRK;ARHGAP6 |
| Negative Regulation of Cell Junction Assembly (GO:1901889) | Regulation of cell junction assembly | 5/23 | 0.00065 | ROBO2;ROCK1;DLC1;ARHGAP6;PHLDB2 |
| Protein Phosphorylation (GO:0006468) | Protein phosphorylation | 16/373 | 0.00065 | PDGFRA;CAMK2D;ROCK1;DAPK1;PRKCA;CD42BPA;MYLK;IGF1R;GRK5;PEAK1;AKT3;MAP3K20;BIRC6;MAP4K3;PRKG1;JAK1 |
| Ventricular Septum Morphogenesis (GO:0060412) | Ventricular septum formation | 5/28 | 0.00140 | ROBO2;BMPR2;SLIT2;ZFPM2;TGFB2 |
| Cardiac Conduction (GO:0061337) | Cardiac electrical conduction | 6/47 | 0.00140 | MEF2A;CACNB2;CAMK2D;CACNA2D1;CACNA1D;CACNA1C |
| Extracellular Structure Organization (GO:0043062) | Extracellular structure organization | 8/111 | 0.00260 | ITGB1;ADAMTS2;COL3A1;COL4A2;COL4A1; MMP2;COL5A2;NPNT |
| Calcium Ion Import (GO:0070509) | Calcium influx | 5/34 | 0.00260 | CACNB2;CACNA2D1;CACNA1D;CACNA1C;SLC8A1 |

**Supplemental Table 3**

| **Figure S4D. KEGG Pathway Enrichment Analysis of Upregulated Genes Identified by Single Nuclear RNA-Sequencing analysis in Mesenchymal Cells** |  |  |  |  |
| --- | --- | --- | --- | --- |
| **Term** | **Short term** | **Overlap** | **Adjusted P-value** | **Genes** |
| Extracellular Matrix Organization (GO:0030198) | ECM organization | 10/182 | 0.00272 | COL1A1;ITGB1;ADAMTS2;COL3A1;COL4A2;COL4A1;MMP2;COL5A2;HMCN1;NPNT |
| External Encapsulating Structure Organization (GO:0045229) | Encapsulating structure organization | 8/111 | 0.00260 | ITGB1;ADAMTS2;COL3A1;COL4A2;COL4A1;MMP2;COL5A2;NPNT |
| Negative Regulation of Cell Migration (GO:0030336) | Suppression of cell migration | 10/184 | 0.00272 | COL3A1;LIMCH1;DLC1;MAGI2;CDH11;LDLRAD4;PTPRK;SLIT2;PHLDB2;PTPRG |
| Positive Regulation of Endothelial Cell Migration (GO:0010595) | Enhanced endothelial cell migration | 7/88 | 0.00294 | NRP1;BCAS3;SPARC;ANGPT1;AKT3;PRKCA;FOXO1 |
| Actomyosin Structure Organization (GO:0031032) | Actomyosin organization | 7/85 | 0.00272 | ITGB1;ROCK1;EPB41L2;MYH9;PHACTR1;CD42BPA;MYH10 |

**Supplemental Table 3**

| **Figure S4E. KEGG Pathway Enrichment Analysis of Upregulated Genes Identified by Single Nuclear RNA-Sequencing analysis in Myeloid Cells** |  |  |  |  |
| --- | --- | --- | --- | --- |
| **Term** | **Short term** | **Overlap** | **Adjusted P-value** | **Genes** |
| Positive Regulation of Signal Transduction (GO:0009967) | Activation of Signal Transduction | 19/256 | 0.00005 | ITGB1;MACF1;CD63;CD74;PRKCB;STAT3;CD180;SORL1;TGFB1;C3;SKI;ZMIZ1;MGAT5;BMP2K;CTSH;PTK2B;PHIP;SNX5;LIMS1 |
| Positive Regulation of DNA-templated Transcription (GO:0045893) | Activation of DNA-templated Transcription | 47/1274 | 0.00005 | KMT2E;NOTCH2;SP100;PID1;MAML2;TCF20;LITAF;STRN3;NSD3;ZMIZ1;NSD1;NAMPT;EP300;APOE;RREB1;SF3B1;SNX5;MEF2A;ATF7IP;NCOA2;CD74;CREBBP;PPP1R12A;STAT1;TCF12;STAT3;DYRK1A;MRTFA;MITF;ARID1B;TGFB1;MED13L;SKI;MLLT10;ZEB2;ELF2;MDFIC;KANSL1;TBL1XR1;KAT6A;IRF2;PHIP;MAML3;CDK13;SQSTM1;PICALM;NFE2L2 |
| Phosphorylation (GO:0016310) | Protein phosphorylation | 20/298 | 0.00005 | GSK3B;PRKCB;CSNK1A1;STAT3;DYRK1A;HCK;TAOK3;WINK1;IRAK2;TAOK1;AKT3;MAPKAPK2;SIK3;TLK1;PTK2B;BIRC6;FNIP2;MAP4K3;JAK1;MAP4K4 |
| Regulation of Protein-Containing Complex Assembly (GO:0043254) | Regulation of protein complex | 12/126 | 0.00033 | RB1;SH3GLB1;LGALS3;GSK3B;HCK;LAMP2;EP300;APOE;FNIP1;TANK;FNIP2;SORL1 |
| Regulation of Small GTPase Mediated Signal Transduction (GO:0051056) | Small GTPase signaling | 13/157 | 0.00048 | VAV3;DENND1A;ARHGAP17;MYO9B;ARHGAP15;ARHGAP25;VAV1;GNA13;PREX1;WINK1;SRGAP2;DOCK2;SQSTM1 |
| Regulation of Cell Migration (GO:0030334) | Regulation of cell migration | 23/479 | 0.00078 | ITGB1;SPAG9;MACF1;CD74;GSK3B;YTHDF3;SEMA4D;STAT3;MITF;DLRAD4;SORL1;TGFB1;CXCL16;SDCBP;MMP14;ADAM17;GPNMB;MGAT5;CTSH;PTK2B;PLXNC1;SRGAP2;MAP4K4 |
| Positive Regulation of RNA Biosynthetic Process (GO:1902680) | Activation of RNA biosynthesis | 24/555 | 0.00177 | KMT2E;ATF7IP;CD74;CREBBP;STAT1;STAT3;DYRK1A;MITF;ARID1B;TGFB1;NSD3;ELF2;MDFIC;KANSL1;TBL1XR1;NSD1;KAT6A;EP300;PHIP;APOE;RREB1;SNX5;PICALM;NFE2L2 |
| Negative Regulation of Apoptotic Process (GO:0043066) | Inhibition of Apoptosis | 21/475 | 0.00342 | RB1;NOTCH2;ITGB1;CD74;GSK3B;SEMA4D;ANXA4;CBL;BABAM2;HCK;SON;TAX1BP1;CTSH;PTK2B;PHIP;BIRC6;CPEB4;LGMN;MAP4K4;GRINA;NFE2L2 |
| Interferon-Mediated Signaling Pathway (GO:0140888) | Interferon signaling | 7/55 | 0.00342 | IFITM3;HCK;SP100;STAT1;IFNGR2;TANK;JAK1 |

**Supplemental Table 3**

| **Figure S4E. KEGG Pathway Enrichment Analysis of Upregulated Genes Identified by Single Nuclear RNA-Sequencing analysis in Myeloid Cells** |  |  |  |  |
| --- | --- | --- | --- | --- |
| **Term** | **Short term** | **Overlap** | **Adjusted P-value** | **Genes** |
| Cellular Response to Transforming Growth Factor Beta Stimulus (GO:0071560) | Transforming Growth Factor Beta response | 9/98 | 0.00342 | ITGB1;SKI;USP15;NRROS;ZMIZ1;STAT3;TGFB1;TGFB2;LIMS1 |
| Negative Regulation of Programmed Cell Death (GO:0043069) | Inhibition of cell death | 18/382 | 0.00464 | NOTCH2;CD74;GSK3B;SEMA4D;ANXA4;CBL;BAM2;HCK;SON;FTH1;TAX1BP1;CTSH;PTK2B;PHIP;BIRC6;SQSTM1;MAP4K4;NFE2L2 |
| Positive Regulation of Cell Motility (GO:2000147) | Activation of cell motility | 14/249 | 0.00484 | ITGB1;SPAG9;SEMA4D;STAT3;TGFB1;CXCL16;SDCBP;MMP14;ADAM17;GPNMB;MGAT5;CTSH;PTK2B;MAP4K4 |
| Transforming Growth Factor Beta Receptor Signaling Pathway (GO:0007179) | Transforming Growth Factor Beta Receptor Signaling | 8/76 | 0.00342 | ITGB1;SKI;USP15;NRROS;ZMIZ1;STAT3;TGFB1;TGFB2 |
| Cellular Response to Tumor Necrosis Factor (GO:0071356) | Tumor Necrosis Factor response | 8/87 | 0.00579 | PID1;STAT1;ERBIN;PTK2B;TNFRSF1B;TANK;LIMS1;NFE2L2 |
| Regulation of Autophagy (GO:0010506) | Autophagy regulation | 13/230 | 0.00600 | SH3GLB1;ATP6V1A;UVRAG;GSK3B;STAT3;EEF1A1;WNK1;EXOC4;PIP4K2A;EP300;TAB2;SQSTM1;SNX5 |
| Regulation of Canonical NF-kappaB Signal Transduction (GO:0043122) | Regulation of Canonical NF-kappaB signaling | 14/270 | 0.00692 | ANKRD17;CD74;PRKCB;STAT1;STAT3;ZC3HAV1;LITAF;TANK;IRAK2;NAMPT;PELI1;TAB2;SQSTM1;LIMS1 |
| Intracellular Signaling Cassette (GO:0141124) | Activation of intracellular signaling | 19/354 | 0.00107 | MEF2A;VAV3;CREBBP;STAT3;GAB2;ARHGAP15;TANK;VAV1;PIK3R5;ADAM17;P2RX4;TAOK3;ZMIZ1;IRAK2;MAPKAPK2;EP300;TAB2;MAP4K4;NFE2L2 |
| Regulation of Gene Expression (GO:0010468) | Regulation of gene expression | 36/1127 | 0.00484 | RB1;KMT2E;NOTCH2;GSK3B;PID1;CELF1;JMJD1C;LITAF;SLC8A1;AFF1;NSD3;NSD1;NAMPT;CTSH;APOE;RREB1;ATF7IP;MSR1;NCOA2;CD74;MBNL1;CREBBP;MBNL2;STAT3;MITF;ASH1L;PARP14;TGFB1;FOXP1;SON;KAT6A;IRF2;LGMN;LIMS1;PICALM;NFE2L2 |
| Negative Regulation of Intracellular Signal Transduction (GO:1902532) | Inhibition of signal transduction | 15/297 | 0.00600 | GSK3B;PHLPP1;STAT1;STAT3;ARHGAP17;LDLRAD4;TANK;ARHGAP25;STRN3;SKI;PREX1;WNK1;IRAK2;TAX1BP1;FNIP1 |
| Positive Regulation of TORC1 Signaling (GO:1904263) | Activation of TORC1 Signaling | 6/52 | 0.00909 | CSNK1A1;CUL3;SIK3;EP300;FNIP1;FNIP2 |

**Supplemental Table 3**

| **Figure S4E. KEGG Pathway Enrichment Analysis of Upregulated Genes Identified by Single Nuclear RNA-Sequencing analysis in Myeloid Cells** |  |  |  |  |
| --- | --- | --- | --- | --- |
| **Term** | **Short term** | **Overlap** | **Adjusted P-value** | **Genes** |
| Response to Lipopolysaccharide (GO:0032496) | Response to Lipopolysaccharide | 10/161 | 0.01181 | ADAM17;CD180;MAPKAPK2;PELI1;ERBIN;TAB2;TNFRSF1B;CD68;B2M;FOXP1 |
| + Reg of Phosphatidylinositol 3-Kinase/Prot Kinase B Signal Transduction (GO:0051897) | Activation of PI3K/PK-B signaling | 10/162 | 0.01181 | ITGB1;VAV3;OSBPL8;P2RX4;SEMA4D;PTK2B;CBL;TGFB1;VAV1;PIK3R5 |
| Regulation of JNK Cascade (GO:0046328) | Regulation of JNK Cascade | 8/115 | 0.01737 | SDCBP;NCOR1;PHLPP1;MDFIC;TAOK3;TAOK1;PTK2B;MAP4K4 |
| Positive Regulation of Cell Growth (GO:0030307) | Regulation of cell growth | 8/107 | 0.01266 | ITSN2;MACF1;SDCBP;ADAM17;MMP14;SEMA4D;TGFB1;CXCL16 |
| post-Golgi Vesicle-Mediated Transport (GO:0006892) | Enhanced protein trafficking and secretion | 6/61 | 0.01573 | VPS13C;EXOC4;VTI1A;VPS54;MYO5A;SORL1 |

**Supplemental Table 3**

| **Figure S7. KEGG Pathway Enrichment Analysis of Upregulated Genes in Alveolar Macrophage Subpopulation 1 (AM1) Identified by Single-Nucleus RNA Sequencing** |  |  |  |  |
| --- | --- | --- | --- | --- |
| **Term** | **Short term** | **Overlap** | **Adjusted P-value** | **Genes** |
| Cellular Response to Low-Density Lipoprotein Particle Stimulus (GO:0071404) | Cellular response to LDL | 4/25 | 0.001908921 | APP;LPL;CD9;TXNIP |
| Myeloid Cell Development (GO:0061515) | Myeloid cell development | 4/26 | 0.001908921 | APP;CEBPB;ANXA2;FLNA |
| Positive Regulation of Endocytosis (GO:0045807) | Regulation of Endocytosis | 5/85 | 0.005972564 | CAMK1D;AXL;PPT1;SIRPA;CD47 |
| Negative Regulation of Programmed Cell Death (GO:0043069) | Negative regulation of Cell death | 9/382 | 0.005972564 | AXL;ARHGDIA;LAPTM5;PPT1;FLNA; CTNNB1;TMBIM6;CD44;PTMS |
| Regulation of Nitric Oxide Biosynthetic Process (GO:0045428) | Nitric oxide biosynthesis | 4/45 | 0.005972564 | APP;HSP90AB1;SIRPA;CD47 |
| Regulation of Phagocytosis (GO:0050764) | Regulation of Phagocytosis | 4/62 | 0.01816753 | CAMK1D;RACK1;SIRPA;CD47 |
| Positive Regulation of Metabolic Process (GO:0009893) | Regulation of metabolism | 5/147 | 0.025782956 | CEBPB;CAMK1D;UCP2;SIRPA;ADIPO R2 |
| Regulation of Apoptotic Process (GO:0042981) | Regulation of Apoptosis | 10/704 | 0.032611119 | APP;AXL;ARHGDIA;RACK1;PPT1;FLNA;CTNNB1;TMBIM6;CD44;PTMS |
| Integrin-Mediated Signaling Pathway (GO:0007229) | Integrin-mediated signaling | 4/86 | 0.025782956 | DST;ITGAX;MYH9;CD47 |
| Regulation of Protein Ubiquitination (GO:0031396) | Protein ubiquitination | 4/113 | 0.044553594 | HSP90AB1;MYCBP2;LAPTM5;CTNN B1 |
| Regulation of Interleukin-6 Production (GO:0032675) | Regulation of IL-6 | 4/114 | 0.044553594 | APP;SIRPA;LPL;CD47 |
| Regulation of Type II Interferon Production (GO:0032649) | Regulation of Type II interferon | 4/88 | 0.025782956 | AXL;LAPTM5;SIRPA;CD47 |
| Mononuclear Cell Differentiation (GO:1903131) | Mononuclear cell differentiation | 3/46 | 0.03805633 | AXL;UCP2;MYH9 |
| Cytoplasmic Translation (GO:0002181) | Cytoplasmic translation | 4/101 | 0.035801798 | RPLP1;RPS5;RACK1;RPL8 |
| Positive Regulation of Multicellular Organismal Process (GO:0051240) | Positive regulation of cell functions | 7/407 | 0.046857519 | APP;CEBPB;UCP2;ITGAX;LPL;CTNNB 1;ADIPOR2 |

**Supplemental Table 3**

| **Figure S7. KEGG Pathway Enrichment Analysis of Upregulated Genes in Alveolar Macrophage Subpopulation 2 (AM2) Identified by Single-Nucleus RNA Sequencing** |  |  |  |  |
| --- | --- | --- | --- | --- |
| **Term** | **Short term** | **Overlap** | **Adjusted P-value** | **Genes** |
| Phagocytosis (GO:0006909) | Phagocytosis | 8/71 | 5.32E-05 | ITGB1;MSR1;ITGAM;TYROBP;CD93;SLC11A1;PLD4;TREM2 |
| Protein Catabolic Process (GO:0030163) | Protein catabolism | 9/107 | 5.32E-05 | MMP14;CTSL;LAMP2;CTS;CTSH;CTSD;CTSS;LGMN;CTSB |
| Macrophage Activation (GO:0042116) | Macrophage activation | 6/35 | 5.32E-05 | C1QA;ITGAM;CD93;SLC11A1;TREM2;TMEM106A |
| Microglial Cell Activation (GO:0001774) | Microglial activation | 5/19 | 5.32E-05 | C1QA;GRN;ITGAM;TYROBP;TREM2 |
| Regulation of Cell Migration (GO:0030334) | Positive regulation of cell migration | 15/479 | 4.79E-04 | ITGB1;CSF1R;IGSF8;CD74;GRN;PLXND1;SOD2;CXCL16;BST2;MMP14;RAP2A;GPNMB;PLXNA1;CTSH;EMILIN2 |
| Positive Regulation of Endocytosis (GO:0045807) | Postitive regulation of endocytosis | 7/85 | 4.79E-04 | SLC11A1;TREM2;CYBA;APOE;CALR;IL2RG;B2M |
| Antigen Processing and Presentation of Exogenous Peptide Antigen (GO:0002478) | Antigen processing and presentation | 5/31 | 4.79E-04 | CTSL;CTSD;CTSS;MPEG1;LGMN |
| Vacuolar Acidification (GO:0007035) | Vacuolar acidification | 5/31 | 4.79E-04 | GRN;ATP6V0B;LAMP1;SLC11A1;LAMP2 |
| Response to Cytokine (GO:0034097) | Response to cytokine | 8/123 | 4.79E-04 | BST2;CSF1R;CD74;PID1;SLC11A1;SELENOP;TIMP2;CXCL16 |
| Protein Metabolic Process (GO:0019538) | Protein metabolism | 17/614 | 4.79E-04 | CTSA;CTS;RRBP1;CTSS;EEF1A1;MMP14;CTSL;ANPEP;LAMP2;CTSH;MAN2B1;APOE;CALR;B2M;CTSD;LGMN;CTSB |
| Positive Regulation of Endothelial Cell Migration (GO:0010595) | Regulation of Endothelial cell migration | 7/88 | 4.79E-04 | NRP1;GRN;P2RX4;BSG;CALR;PLPP3;LGMN |
| Integrin-Mediated Signaling Pathway (GO:0007229) | Integrin-mediated signaling | 6/86 | 0.002937 | ITGB1;NRP1;ITGAM;ITGB5;FN1;PLPP3 |
| Regulation of Interleukin-6 Production (GO:0032675) | Regulation of IL-6 production | 7/114 | 0.002193 | CD74;TYROBP;GPNMB;BSG;CYBA;TREM2;TMEM106A |
| Proteolysis Involved in Protein Catabolic Process (GO:0051603) | Proteolysis | 6/80 | 0.00241 | CTSL;CTS;CTSH;CTSS;CTSB;LGMN |
| Positive Regulation of Response to External Stimulus (GO:0032103) | Response to external stimulus | 8/163 | 0.00241 | LGALS3;CEBPA;GRN;LGALS1;EMILIN2;CYBA;TREM2;PLA2G7 |

**Supplemental Table 3**

| **Figure S7. KEGG Pathway Enrichment Analysis of Upregulated Genes in Alveolar Macrophage Subpopulation 2 (AM2) Identified by Single-Nucleus RNA Sequencing** |  |  |  |  |
| --- | --- | --- | --- | --- |
| **Term** | **Short term** | **Overlap** | **Adjusted P-value** | **Genes** |
| Positive Regulation of Cell Population Proliferation (GO:0008284) | Positive regulation of cell proliferation | 11/484 | 0.01906 | BST2;CSF1R;CD74;GRN;PID1;FN1;CTSH;CALR;IL7R;LGMN;GNAI2 |
| Proteolysis (GO:0006508) | Proteolysis | 10/321 | 0.006047 | CTSA;MMP14;CTSL;ANPEP;CTSZ;CTSH;CTSD;CTSS;LGMN;CTSB |
| Osteoclast Differentiation (GO:0030316) | Osteoclast differentiation | 4/24 | 0.002326 | CSF1R;TYROBP;ATP6AP1;TREM2 |
| Regulation of Fibroblast Proliferation (GO:0048145) | Fibroblast proliferation | 4/46 | 0.010828 | CD74;CD300A;FTH1;FN1 |

**Supplemental Table 3**

| **Figure S7. KEGG Pathway Enrichment Analysis of Upregulated Genes in Alveolar Macrophage Subpopulation 3 (AM3) Identified by Single-Nucleus RNA Sequencing** |  |  |  |  |
| --- | --- | --- | --- | --- |
| **Term** | **Short term** | **Overlap** | **Adjusted P-value** | **Genes** |
| Protein Phosphorylation (GO:0006468) | Protein phosphorylation | 18/373 | 0.003531 | SRPK2;MAP3K3;PRKCB;DAPK1;NEK7;VRK2;NLK;MERTK;MAPK14;TGFB1;HIPK2;ERN1;TBCK;TEC;TLK1;SIK2;MAP4K3;MAP4K4 |
| Regulation of Small GTPase Mediated Signal Transduction (GO:0051056) | Regulation of signal transduction | 11/157 | 0.005097 | ARHGAP22;FGD4;OPHN1;DENND1A;DENND4C;DOCK7;ARHGAP15;DOCK2;ARHGAP25;ARHGAP6;ARHGAP12 |
| Positive Regulation of Transcription by RNA Polymerase II (GO:0045944) | Regulation of transcription | 28/983 | 0.024613 | PRKN;MAML2;FOXO3;CDC73;APBB2;RREB1;MAP2K5;MLXIP;STAT5B;NCOA2;WVVOX;BCAS3;ZFHX3;KDM4C;TCF12;MRTFB;MRTFA;MITF;IGF1;ST18;HIPK2;ASXL1;MLLT10;MYO1E;NFIA;MAML3;ERCC6;PLPP3 |
| Protein Modification by Small Protein Conjugation (GO:0032446) | Protein modification | 13/316 | 0.041117 | PRKN;UHRF2;FBXW7;NSMCE2;ATG10;FBXL17;WWP1;FBXO32;PIAS2;ITCH;ZNRFB3;ATG7;RNF111 |
| Macroautophagy (GO:0016236) | Macroautophagy | 9/162 | 0.041117 | PRKN;UVRAG;TBC1D5;ATG10;VTI1A;PIP4K2A;WDFY3;AMBRA1;ATG7 |
| Protein Ubiquitination (GO:0016567) | Protein ubiquitination | 14/369 | 0.046631 | PRKN;UBE2H;UBE3C;RNF13;UHRF2;FBXW7;FBXL17;WWP1;AMBRA1;FBXO32;ITCH;ZNRFB3;RNF111;SASH1 |
| Regulation of Autophagy (GO:0010506) | Autophagy regulation | 10/230 | 0.086822 | RPTOR;PRKN;ERN1;UVRAG;DAPK1;EXOC4;PIP4K2A;AMBRA1;FOXO3;ATP6V0D2 |
| post-Golgi Vesicle-Mediated Transport (GO:0006892) | Vesicle-mediated transport | 7/61 | 0.005772 | KIF16B;VPS13C;EXOC4;VTI1A;EXOC6B;MYO5A;EXOC2 |
| Cytosolic Transport (GO:0016482) | Cytosolic transport | 8/99 | 0.015078 | TMEM87B;PIKFYVE;KIF16B;TBC1D5;VPS50;VPS13C;VTI1A;TBC1D23 |
| Membrane Fission (GO:0090148) | Membrane fission | 5/42 | 0.037657 | MVB12B;EXOC6B;EXOC4;TMCC1;EXOC2 |
| Organelle Organization (GO:0006996) | Organelle organization | 14/437 | 0.100448 | PRKN;DST;COG5;MAPKAP1;VPS13C;VPS13B;FGD4;TMEM135;PIKFYVE;ZMYM4;ATXN7;DYM;TMCC1;ATP7A |

Supplemental Table 3

| Figure S7. KEGG Pathway Enrichment Analysis of Upregulated Genes in Alveolar Macrophage Subpopulation 4 (AM4) Identified by Single-Nucleus RNA Sequencing |  |  |  |  |
| --- | --- | --- | --- | --- |
| Term | Short term | Overlap | Adjusted P-value | Genes |
| Defense Response to Virus (GO:0051607) | Response to virus | 24/198 | 7.55E-21 | IFITM3;ZBP1;RTP4;GBP5;GBP7;RSD2;STAT2;NLRC5;EIF2AK2;ISG15;IFI T1;DDX60;SAMHD1;USP18;TANK;IFI T3;IFIT2;IFIH1;CXCL10;ZNFX1;IRF1; OAS3;IRF7;GBP2 |
| Negative Regulation of Viral Process (GO:0048525) | Regulation of viral process | 10/66 | 7.99E-09 | IFITM3;IFIH1;ZFP36;SP100;ZNFX1;R SAD2;OAS3;EIF2AK2;ISG15;IFIT1 |
| Negative Regulation of Innate Immune Response (GO:0045824) | Regulation of innate immune response | 9/57 | 3.92E-08 | OAS3;STAT2;NLRC5;SLAMF8;TNFAIP 3;TRAFD1;ISG15;SAMHD1;USP18 |
| Antiviral Innate Immune Response (GO:0140374) | Antiviral innate immune response | 9/62 | 6.46E-08 | IFIH1;ZBP1;CXCL10;OAS3;EIF2AK2;I FIT1;USP18;IFIT3;IFIT2 |
| Negative Regulation of Cytokine-Mediated Signaling Pathway (GO:0001960) | Cytokine-mediated signaling | 9/67 | 8.83E-08 | USP25;IL1RN;OAS3;STAT2;NLRC5;IS G15;SAMHD1;PARP14;USP18 |
| Cellular Response to Type II Interferon (GO:0071346) | Response to Type II interferon | 8/47 | 1.01E-07 | GBP5;SP100;GBP7;IRF1;CCL2;LGALS 9;GBP2;JAK2 |
| Regulation of Type I Interferon-Mediated Signaling Pathway (GO:0060338) | Type I interferon-mediated signaling | 7/36 | 3.23E-07 | ZBP1;STAT2;OAS3;NLRC5;ISG15;SA MHD1;USP18 |
| Positive Regulation of Cytokine-Mediated Signaling Pathway (GO:0001961) | Positive regulation of cytokine mediated signaling | 6/47 | 4.50E-05 | ZBP1;CD40;CASP4;NLRC5;PARP14;P ARP9 |
| Regulation of Innate Immune Response (GO:0045088) | Regulation of innate immune response | 7/86 | 9.94E-05 | GBP5;IRF1;TNFAIP3;TRAFD1;IFI35;P ARP9;SAMHD1 |
| Cellular Response to Lipopolysaccharide (GO:0071222) | Response to lipopolysaccharide | 8/125 | 1.02E-04 | CD86;CXCL10;ZFP36;TNFAIP3;CCL2; JAK2;GBP2;NFKB1 |
| Cellular Response to Cytokine Stimulus (GO:0071345) | Response to cytokine stimulus | 11/289 | 1.82E-04 | GBP5;ZFP36;GBP7;IRF1;CCL2;LGALS 9;IFIT1;JAK2;GBP2;TANK;NFKB1 |
| Inflammatory Response (GO:0006954) | Inflammatory response | 10/261 | 4.02E-04 | ZBP1;CXCL10;IL1RN;CD40;NAMPT;N INJ1;CASP4;CCL2;LGALS9;NFKB1 |
| Defense Response to Bacterium (GO:0042742) | Response to bacterium | 9/221 | 6.29E-04 | GBP7;RNF213;ZNFX1;OAS3;CASP4;S LAMF8;ISG15;GBP2;NFKB1 |

**Supplemental Table 3**

|  |  |  |  |  |
| --- | --- | --- | --- | --- |
| Negative Regulation of Gene Expression<br>(GO:0010629) | Regulation of gene expression | 11/364 | 9.22E-04 | SRGN;ZFP36;DAXX;REST;GBP7;KDR;<br>EIF2AK2;LGALS9;PARP14;PARP9;NF<br>KB1 |
| Positive Regulation of Cytokine<br>Production (GO:0001819) | Positive regulation of cytokine<br>production | 11/325 | 4.34E-04 | CD86;IFIH1;GBP5;IRF1;IRF7;EIF2AK2<br>;ISG15;LGALS9;JAK2;TANK;MALT1 |

**Supplemental Table 3**

| **Figure S7. KEGG Pathway Enrichment Analysis of Upregulated Genes in Alveolar Macrophage Subpopulation 5 (AM5) Identified by Single-Nucleus RNA Sequencing** |  |  |  |  |
| --- | --- | --- | --- | --- |
| **Term** | **Short term** | **Overlap** | **Adjusted P-value** | **Genes** |
| Positive Regulation of Intracellular Signal Transduction (GO:1902533) | Positive regulation of signal transduction | 18/652 | 2.48E-04 | SMAD3;RSAD2;SEMA4D;DDX60;THBS1;VAV1;MALT1;IGF1R;IRAK2;IL1B;NAMPT;CCL4;CCL3;PTK2B;JAK2;FNIP1;PIK3AP1;TLR2 |
| Defense Response to Virus (GO:0051607) | Response to virus | 10/198 | 4.02E-04 | RSAD2;IRF1;IRF2;NLRC5;ACOD1;IFIT1;DDX60;LYST;BCL2L1;TLR2 |
| + Reg of Phosphatidylinositol 3-Kinase/Prot Kinase B Signal Transduction (GO:0051897) | Regulation of PI3K/PK signal | 9/162 | 4.54E-04 | SEMA4D;IL1B;CCL3;PTK2B;JAK2;PIK3AP1;THBS1;VAV1;IGF1R |
| Regulation of Inflammatory Response (GO:0050727) | Regulation of inflammatory response | 10/249 | 0.00121222 | NFKBIA;TNIP1;IL1B;CCL3;ACOD1;TNFAIP3;JAK2;TNFRSF1B;PIK3AP1;TLR2 |
| Positive Regulation of Cell Motility (GO:2000147) | Positive regulation of cell migration | 10/249 | 0.00121222 | SPAG9;CSF1;SEMA4D;SMOX;IL1B;CCL4;CCL3;PTK2B;THBS1;IGF1R |
| Cytokine-Mediated Signaling Pathway (GO:0019221) | Cytokine mediated signaling | 10/255 | 0.00121222 | NFKBIA;CSF3R;IRAK2;IL1B;CCL4;CCL3;PTK2B;IL1RAP;JAK2;TNFRSF1B |
| Inflammatory Response (GO:0006954) | Inflammatory response | 10/261 | 0.00121222 | IL1RN;IRAK2;IL1B;CCRL2;NAMPT;MAPKAPK2;CCL4;CCL3;THBS1;TLR2 |
| Cellular Response to Type II Interferon (GO:0071346) | Type II interferon response | 5/47 | 0.00158751 | SP100;IRF1;CCL3;JAK2;TLR2 |
| Response to Cytokine (GO:0034097) | Response to cytokine | 7/123 | 0.00161524 | CD274;SP100;IRAK2;IL1B;MAPKAPK2;JAK2;BCL2L1 |
| Regulation of Tumor Necrosis Factor Production (GO:0032680) | Regulation of TNF-α production | 7/131 | 0.00173433 | ARFGEF2;MAPKAPK2;CCL3;TNFAIP3;JAK2;THBS1;TLR2 |
| Cellular Response to Interleukin-1 (GO:0071347) | Response to Interleukin-1 | 5/57 | 0.00217661 | NFKBIA;IRAK2;IL1B;CCL3;IL1RAP |
| Cell Surface Toll-Like Receptor Signaling Pathway (GO:0140895) | Toll-like receptor signaling | 4/30 | 0.00237285 | NFKBIA;IRAK2;PIK3AP1;TLR2 |

**Supplemental Table 3**

| **Figure S7. KEGG Pathway Enrichment Analysis of Upregulated Genes in Alveolar Macrophage Subpopulation 5 (AM5) Identified by Single-Nucleus RNA Sequencing** |  |  |  |  |
| --- | --- | --- | --- | --- |
| **Term** | **Short term** | **Overlap** | **Adjusted P-value** | **Genes** |
| Cellular Response to Lipopolysaccharide (GO:0071222) | Response to Lipopolysaccharide | 7/125 | 0.00165634 | CD274;TNIP1;IRAK2;IL1B;TNFAIP3;JAK2;TNFRSF1B |
| Positive Regulation of Macromolecule Biosynthetic Process (GO:0010557) | Positive regulation of Macromolecule biosynthesis | 10/355 | 0.00381215 | SMAD3;CSF1;SMOX;IL1B;NAMPT;NLRC5;CCL3;JAK2;NFE2L2;TLR2 |
| Positive Regulation of DNA-templated Transcription (GO:0045893) | Regulation of transcription | 21/1274 | 0.00310108 | ATF7IP;SP100;SMAD3;SMOX;DYRK1A;NLRC5;ETS2;RUNX1;NFKBIA;KLF6;ELF2;TNIP1;IL1B;IRF1;NAMPT;IRF2;JAK2;ATF3;NFE2L2;PICALM;TLR2 |
| Regulation of Cell Population Proliferation (GO:0042127) | Regulation of cell proliferation | 12/769 | 0.04062219 | CSF3R;SMAD3;CSF1;IL1B;IRF1;NAMPT;MDM2;PTK2B;JAK2;THBS1;IGF1R;RUNX1 |

**Supplemental Table 3**

| **Figure S8D. KEGG Pathway Enrichment Analysis of Upregulated Genes in Gene Module 1 Identified by Pseudotime Trajectory Analysis** |  |  |  |  |
| --- | --- | --- | --- | --- |
| **Term** | **Short term** | **Overlap** | **Adjusted P-value** | **Genes** |
| Regulation of Intracellular Signal Transduction (GO:1902531) | Intracellular signal transduction | 15/340 | 8.47E-05 | LYN;VAV3;CSF1R;DOCK8;ITPR1;ARHGAP15;ARHGAP26;MOB3B;AKAP13;FGD4;ARHGAP30;RASA3;TAOK3;TAB2;DOCK2 |
| Positive Regulation of Protein Kinase Activity (GO:0045860) | Protein kinase regulation | 7/57 | 1.85E-04 | CSF1R;CCDC88A;RAP1A;PTPRC;NEDD9;LCP2;TAB2 |
| Regulation of Small GTPase Mediated Signal Transduction (GO:0051056) | Small GTPase mediated signaling | 10/164 | 2.64E-04 | VAV3;DOCK11;AKAP13;FGD4;ARHGAP30;DENND4A;DOCK8;ARHGAP15;ARHGAP26;DOCK2 |
| Fc-gamma Receptor Signaling Pathway Involved in Phagocytosis (GO:0038096) | Fc gamma receptor signaling | 5/23 | 2.64E-04 | LYN;VAV3;FGR;PRKCE;PRKCD |
| B Cell Activation (GO:0042113) | B cell activation/BCR signaling | 8/103 | 3.04E-04 | TPD52;LYN;MEF2C;PTPRC;ITGA4;NFAM1;PIK3CD;PTPRJ |
| Lymphocyte Differentiation (GO:0030098) | Lymphocyte differentiation | 7/81 | 5.35E-04 | TPD52;PTPRC;ITGA4;NFAM1;PIK3CD;PTPRJ;IKZF1 |
| Intracellular Signaling Cassette (GO:0141124) | Intracellular signal cassette | 13/362 | 5.83E-04 | VAV3;CX3CR1;MEF2C;PRKCD;PIK3CD;ARHGAP15;CAMKK2;CCDC88A;AKAP13;ARHGAP30;TAOK3;TAB2;CCR2 |
| Protein Modification Process (GO:0036211) | Posttranslational modification | 14/439 | 8.40E-04 | LYN;PPP1R12A;PRKCD;F13A1;PIK3CD;SSH2;CAMKK2;FGR;STK10;PTPRC;STK24;TAOK3;STK17B;ST8SIA4 |
| Antigen Receptor-Mediated Signaling Pathway (GO:0050851) | Immune signaling | 7/120 | 0.00347783 | VAV3;MEF2C;PTPRC;NFAM1;PIK3CD;PTPRJ;LCP2 |
| B Cell Homeostasis (GO:0001782) | B cell homeostasis | 3/10 | 0.003967283 | LYN;DOCK11;MEF2C |
| Positive Regulation of Leukocyte Apoptotic Process (GO:2000108) | Regulation of leucocyte apoptosis | 3/10 | 0.003967283 | LYN;MEF2C;PIK3CD |
| Regulation of Mononuclear Cell Migration (GO:0071675) | Mononuclear cell migration | 3/12 | 0.006812952 | LYN;CSF1R;STK10 |

**Supplemental Table 3**

| **Figure S8D. KEGG Pathway Enrichment Analysis of Upregulated Genes in Gene Module 1 Identified by Pseudotime Trajectory Analysis** |  |  |  |  |
| --- | --- | --- | --- | --- |
| **Term** | **Short term** | **Overlap** | **Adjusted P-value** | **Genes** |
| Regulation of Actin Filament-Based Process (GO:0032970) | Actin cytoskeleton organization | 6/98 | 0.00731309 | CSF1R;CCDC88A;PRKCE;PRKCD;NEDD9;CORO1A |
| Hemopoiesis (GO:0030097) | Hemopoiesis | 6/102 | 0.008328927 | NOTCH2;LYN;CSF1R;PTPRC;KLF3;ETV6 |
| Regulation of Cell Junction Assembly (GO:1901888) | Cell junction assembly | 4/39 | 0.011546923 | RAP1B;MEF2C;RAP1A;PDLIM5 |
| Regulation of MAPK Cascade (GO:0043408) | MAPK pathway regulation | 9/292 | 0.02087605 | LYN;CSF1R;AKAP13;MEF2C;PTPRC;TAOK3;PRKCD;TAB2;PTPRJ |
| B Cell Proliferation (GO:0042100) | B cell proliferation | 3/20 | 0.02087605 | LYN;MEF2C;PTPRC |
| Positive Regulation of Cytokine Production (GO:0001819) | Cytokine production | 10/358 | 0.02087605 | FGR;CSF1R;PTPRC;NFAM1;PIK3CD;POU2F2;PLCB1;SORL1;CCR2;IL17RA |
| Regulation of Monocyte Extravasation (GO:2000437) | Monocyte extravasation | 2/5 | 0.02431396 | PLCB1;CCR2 |
| Regulation of Insulin Receptor Signaling Pathway (GO:0046626) | Insulin receptor signaling | 4/53 | 0.027717056 | PID1;PRKCD;PIP4K2A;SORL1 |
| Phosphatidylinositol-mediated Signaling (GO:0048015) | PI3K signaling | 3/24 | 0.029285294 | PLCL1;PIK3CD;PLCB1 |
| Regulation of Tumor Necrosis Factor Production (GO:0032680) | TNF production | 6/146 | 0.032692798 | CX3CR1;PTPRC;IFNGR1;PTPRJ;VSIR;CCR2 |

**Supplemental Table 3**

| **Figure S8D. KEGG Pathway Enrichment Analysis of Upregulated Genes in Gene Module 2 Identified by Pseudotime Trajectory Analysis** |  |  |  |  |
| --- | --- | --- | --- | --- |
| **Term** | **Short term** | **Overlap** | **Adjusted P-value** | **Genes** |
| Positive Regulation of Endocytosis (GO:0045807) | Endocytosis regulation | 10/85 | 1.02E-06 | C3;LRP1;AXL;SLC11A1;TREM2;CYBA;APOE;CALR;IL2RG;B2M |
| Protein Metabolic Process (GO:0019538) | Protein metabolism | 20/669 | 7.90E-05 | RPL4;CTSA;CTSZ;RRBP1;CTSS;CNDP2;EEF1A1;C3;MMP14;CAPNS1;CTSL;TPP1;APOE;CALR;B2M;CTSD;CFB;LGMN;CTSB;MSRB1 |
| Antigen Processing and Presentation of Exogenous Peptide Antigen (GO:0002478) | Antigen processing/presentation | 6/36 | 1.19E-04 | CTSL;B2M;CTSD;CTSS;MPEG1;LGMN |
| Semaphorin-plexin Signaling Pathway (GO:0071526) | Semaphorin-plexin signaling | 6/41 | 1.79E-04 | TYROBP;PLXND1;ARHGDI1;PLXNA1;PLXNB2;TREM2 |
| Positive Regulation of Phagocytosis (GO:0050766) | Phagocytosis regulation | 7/67 | 1.79E-04 | C3;SLC11A1;TREM2;CYBA;CALR;IL2RG;CFB |
| Protein Catabolic Process (GO:0030163) | Protein catabolism | 10/178 | 1.79E-04 | SEM1;MMP14;PDCD6IP;CTSL;CTSZ;TPP1;CTSD;CTSS;LGMN;CTSB |
| Negative Regulation of Apoptotic Process (GO:0043066) | Apoptosis regulation | 16/497 | 1.79E-04 | CD74;GRN;HSPA5;TREM2;IL2RG;SOX2;HSP90B1;PTMS;PRDX5;AXL;ARHGDI1;CFL1;TM6SF1;MCL1;LGMN;GRIN1 |
| Macromolecule Catabolic Process (GO:0009057) | Macromolecule catabolism | 7/74 | 2.31E-04 | MMP14;CTSL;CTSZ;TPP1;CTSS;CTSB;LGMN |
| Proteolysis (GO:0006508) | Proteolysis | 13/358 | 3.52E-04 | CTSA;CTSZ;CTSS;CNDP2;C3;MMP14;CAPNS1;CTSL;CPD;TPP1;CTSD;LGMN;CTSB |
| Positive Regulation of Biosynthetic Process (GO:0009891) | Regulation of biosynthesis | 7/91 | 6.00E-04 | MTPN;PRDX5;HSP90AB1;P2RX4;CYBA;APOE;VIM |
| Cellular Response to Oxygen-Containing Compound (GO:1901701) | Metabolic adaptation | 13/382 | 6.00E-04 | LRP1;TREM2;HSP90B1;PRDX5;TYROBP;P2RX4;AXL;GNAS;VIM;CCR5;CD68;B2M;LGMN |
| T Cell Activation Involved in Immune Response (GO:0002286) | T cell activation | 5/35 | 6.00E-04 | CD74;TYROBP;SLC11A1;PLXNA1;TREM2 |
| Vacuolar Acidification (GO:0007035) | Vacuolar acidification | 5/36 | 6.45E-04 | GRN;LAMP1;ATP6AP1;CREG1;SLC11A1 |
| Glycolipid Transport (GO:0046836) | Glycolipid transport | 3/6 | 6.45E-04 | NPC2;PSAP;PLTP |

**Supplemental Table 3**

| **Figure S8D. KEGG Pathway Enrichment Analysis of Upregulated Genes in Gene Module 2 Identified by Pseudotime Trajectory Analysis** |  |  |  |  |
| --- | --- | --- | --- | --- |
| **Term** | **Short term** | **Overlap** | **Adjusted P-value** | **Genes** |
| T Cell Act via T Cell Rec Contact With Antigen Bound to MHC Mol on Antigen Presenting Cell (GO:0002291) | Antigen presentation to T cells | 3/6 | 6.45E-04 | TYROBP;PLXNA1;TREM2 |
| Phagocytosis (GO:0006909) | Phagocytosis | 7/98 | 6.45E-04 | MSR1;TYROBP;LRP1;AXL;SLC11A1;MYH9;TREM2 |
| Negative Regulation of Programmed Cell Death (GO:0043069) | Programmed cell death | 13/410 | 8.37E-04 | CD74;HSPA5;TREM2;IL2RG;HSP90B1;PTMS;PRDX5;AXL;FTH1;ARHGDIA;CFL1;TM6IM6;MCL1 |
| Cellular Response to Thyroid Hormone Stimulus (GO:0097067) | Thyroid hormone stimulus | 3/7 | 8.94E-04 | CTSL;CTSS;CTSB |
| Microglial Cell Activation Involved in Immune Response (GO:0002282) | Microglial cell activation | 3/7 | 8.94E-04 | GRN;TYROBP;TREM2 |
| Positive Regulation of Amyloid-Beta Clearance (GO:1900223) | Amyloid-beta clearance | 3/7 | 8.94E-04 | LRP1;TREM2;APOE |
| Lysosomal Lumen Acidification (GO:0007042) | Lysosomal acidification | 4/23 | 0.00132 | GRN;ATP6AP1;LAMP1;CREG1 |
| Positive Regulation of Cholesterol Efflux (GO:0010875) | Regulation of cholesterol efflux | 4/23 | 0.00132 | LRP1;TREM2;APOE;PLTP |

**Supplemental Table 3**

| **Figure S8D. KEGG Pathway Enrichment Analysis of Upregulated Genes in Gene Module 3 Identified by Pseudotime Trajectory Analysis** |  |  |  |  |
| --- | --- | --- | --- | --- |
| **Term** | **Short term** | **Overlap** | **Adjusted P-value** | **Genes** |
| Translation (GO:0006412) | Protein biosynthesis | 7/251 | 0.0012 | RPS14;RPS9;RPLP1;RPS5;RACK1;RPL8;EEF2 |
| Epiboly Involved in Wound Healing (GO:0090505) | Epiboly in wound healing | 3/15 | 0.001572 | FLNA;LCP1;CD44 |
| Regulation of Phagocytosis (GO:0050764) | Phagocytosis regulation | 4/63 | 0.002942 | CAMK1D;RACK1;SIRPA;CD47 |
| Regulation of Biosynthetic Process (GO:0009889) | Biological process regulation | 3/24 | 0.003734 | HVCN1;SIRPA;CD47 |
| Wound Healing, Spreading of Cells (GO:0044319) | Cell migration | 3/28 | 0.005144 | FLNA;LCP1;CD44 |
| Myeloid Cell Development (GO:0061515) | Myeloid cell development | 3/34 | 0.008133 | CEBPB;ANXA2;FLNA |
| Negative Regulation of Endocytosis (GO:0045806) | Endocytosis regulation | 3/36 | 0.008595 | RACK1;SIRPA;CD47 |
| Regulation of Tumor Necrosis Factor Superfamily Cytokine Production (GO:1903555) | Regulation of TNF production | 2/9 | 0.016699 | SIRPA;CD47 |
| Positive Regulation of Leukocyte Cell-Cell Adhesion (GO:1903039) | Regulation of leucocyte regulation | 3/51 | 0.019561 | SIRPA;CD47;CD44 |
| Cellular Response to Interleukin-12 (GO:0071349) | Interleuking-12 cellular response | 2/11 | 0.019561 | SIRPA;CD47 |
| Regulation of Nitric Oxide Metabolic Process (GO:0080164) | Nitric oxide metabolism | 2/11 | 0.019561 | SIRPA;CD47 |
| Macromolecule Biosynthetic Process (GO:0009059) | Macromolecule biosynthesis | 5/243 | 0.019895 | RPS14;RPLP1;RPS5;RPL8;EEF2 |
| Positive Regulation of Transport (GO:0051050) | Regulation of transport | 3/59 | 0.022467 | CEBPB;ANXA2;ACTB |
| Actin Crosslink Formation (GO:0051764) | Actin crosslink formation | 2/13 | 0.022467 | FLNA;LCP1 |
| Regulation of Cholesterol Storage (GO:0010885) | Cholesterol homeostasis | 2/16 | 0.028965 | LPL;ABCG1 |
| Regulation of T Cell Activation (GO:0050863) | Regulation of T cell activation | 3/83 | 0.044548 | LAPTM5;SIRPA;CD47 |
| High-density Lipoprotein Particle Remodeling (GO:0034375) | Remodeling of HDL particles | 2/21 | 0.044548 | LPL;ABCG1 |

**Supplemental Table 3**

| Figure S8D. KEGG Pathway Enrichment Analysis of Upregulated Genes in Gene Module 3 Identified by Pseudotime Trajectory Analysis |  |  |  |  |
| --- | --- | --- | --- | --- |
| Term | Short term | Overlap | Adjusted P-value | Genes |
| Positive Regulation of Endocytosis<br>(GO:0045807) | Endocytosis regulation | 3/85 | 0.044548 | CAMK1D;SIRPA;CD47 |
| Ribosomal Small Subunit Biogenesis<br>(GO:0042274) | Ribosomal subunit biogenesis | 3/88 | 0.047088 | RPS14;RPS9;RPS5 |
| Carbohydrate Derivative Transport<br>(GO:1901264) | Organic substance transport | 2/24 | 0.052057 | ABCG1;CD44 |

**Supplemental Table 3**

| **Figure S8D. KEGG Pathway Enrichment Analysis of Upregulated Genes in Gene Module 4 Identified by Pseudotime Trajectory Analysis** |  |  |  |  |
| --- | --- | --- | --- | --- |
| **Term** | **Short term** | **Overlap** | **Adjusted P-value** | **Genes** |
| Regulation of Intracellular Signal Transduction (GO:1902531) | Intracellular signaling | 20/340 | 0.001805 | APP;TNFAIP8;TRIO;FAM13B;MYO9B;ARHGAP6;DHRS3;SLC8A1;HIPK2;STRN3;ARHGAP22;TBCK;ARHGAP10;DLG1;GNA12;NF1;RICTOR;PLPP3;SRGAP2;MAP4K4 |
| Protein Ubiquitination (GO:0016567) | Protein ubiquitination | 21/372 | 0.001805 | PRKN;RNF13;UHRF2;FBXW11;FBXW7;TNKS;FBXL17;RC3H1;NEDD4L;WWP1;FBXO32;CBL;FBXO11;MIB1;ZNR3;SPOP;TRIP12;ARIH1;RNF111;SASH1;PCMTD1 |
| Regulation of Small GTPase Mediated Signal Transduction (GO:0051056) | GTPase mediated signaling | 13/164 | 0.00337 | GIT2;TRIO;FAM13B;DENND1A;DENND4C;MYO9B;ARHGAP25;ARHGAP6;ARHGAP22;ARHGAP10;OPHN1;RAF1;SRGAP2 |
| Protein Modification by Small Protein Conjugation (GO:0032446) | Posttranslational modification | 18/313 | 0.003818 | PRKN;UHRF2;FBXW11;FBXW7;FBXL17;NEDD4L;WWP1;FBXO32;CBL;FBXO11;MIB1;PIAS1;ZNR3;ZMIZ1;ARIH1;ATG7;RNF111;PCMTD1 |
| Positive Regulation of Cell Migration (GO:0030335) | Regulation of cell migration | 19/349 | 0.003818 | BCAS3;NRP2;SMURF2;IGF1;TGFB1;SHTN1;PPP3CA;RAP2A;APC;MGAT5;RAPGEF2;RICTOR;HPSE;PLPP3;RREB1;DOCK1;SASH1;MAP4K4;CLASP2 |
| Wound Healing (GO:0042060) | Wound healing | 9/83 | 0.004133 | PPP3CA;MACF1;DST;MYOF;NF1;IGF1;HPSE;RAF1;TGFB1 |
| Modification-dependent Protein Catabolic Process (GO:0019941) | Protein catabolic process | 14/224 | 0.009462 | PRKN;SMURF2;RNF13;FBXW11;RC3H1;NEDD4L;CBL;FBXO11;MIB1;ZNR3;TRIP12;ARIH1;RNF130;RNF111 |
| Cellular Response to Insulin Stimulus (GO:0032869) | Insulin stimulus response | 9/109 | 0.027391 | BCAR3;RAB31;PIK3CA;APC;MAPKAP1;AKT3;DENND4C;MYO5A;IGF1 |
| Negative Regulation of Granulocyte Differentiation (GO:0030853) | Inhibition of granulocyte differentiation | 3/7 | 0.0299 | INPP5D;C1QC;RUNX1 |
| Receptor Catabolic Process (GO:0032801) | Receptor catabolic process | 4/17 | 0.0299 | SH3GLB1;UVRAG;ZNR3;NEDD4L |
| Astrocyte Development (GO:0014002) | Astrocyte immune response | 4/17 | 0.0299 | APP;C1QA;CDK6;NF1 |

**Supplemental Table 3**

| **Figure S8D. KEGG Pathway Enrichment Analysis of Upregulated Genes in Gene Module 4 Identified by Pseudotime Trajectory Analysis** |  |  |  |  |
| --- | --- | --- | --- | --- |
| **Term** | **Short term** | **Overlap** | **Adjusted P-value** | **Genes** |
| Positive Regulation of DNA-templated Transcription (GO:0045893) | DNA templated transcription | 40/1304 | 0.031593 | PRKN;APP;MAML2;TNKS;FOXO3;STRN3;LRP6;PPP3CA;SCAF8;ZMIZ1;NSD1;APBB2;RREB1;MAP2K5;MEF2A;MLXIP;WWOX;BCAS3;FBXW11;TCF12;MRTFB;MRTFA;MITF;IGF1;ARID1B;ST18;TGFB1;RUNX1;MED13L;HIPK2;PIAS1;MLLT10;ZEB2;MDFIC;NFIA;KANS1;ASXL2;ERCC6;PLPP3;PICALM |
| Negative Regulation of Muscle Cell Apoptotic Process (GO:0010656) | Apoptosis regulation | 4/18 | 0.031593 | IGF1;LARGE1;MAP2K5;LRP6 |
| Neuron Development (GO:0048666) | Neuro immune response | 11/177 | 0.031593 | APP;OPHN1;TAOK1;NEDD4L;VPS13B;RAPGEF2;THOC2;TBC1D23;ERCC6;LRP12;RUNX1 |
| Regulation of JNK Cascade (GO:0046328) | JNK signaling cascade | 9/122 | 0.031593 | PRKN;APP;RAP2A;RNF13;MDFIC;TAOK1;FCGR2B;MAP4K4;HIPK2 |
| Regulation of GTPase Activity (GO:0043087) | GTPase regulation | 8/97 | 0.031593 | BCAR3;LRCH1;RABGAP1;NF1;SNX13;RAPGEF2;SRGAP2;GPR137B |
| Neuron Projection Development (GO:0031175) | Neuro-immune remodeling | 12/214 | 0.037191 | APP;SHTN1;TRIO;OPHN1;NEDD4L;RAPGEF2;VPS13B;TBC1D23;ERCC6;SRGAP2;LRP12;MAP4K4 |
| Regulation of Postsynapse Organization (GO:0099175) | Postsynapse organization | 5/36 | 0.03866 | ARHGAP22;APP;FCGR2B;DOCK1;TANC2 |
| Peptidyl-serine Phosphorylation (GO:0018105) | Protein phosphorylation | 5/37 | 0.041883 | SMG1;PIKFYVE;TNKS;TGFB1;HIPK2 |

**Supplemental Table 3**

| **Figure S8D. KEGG Pathway Enrichment Analysis of Upregulated Genes in Gene Module 5 Identified by Pseudotime Trajectory Analysis** |  |  |  |  |
| --- | --- | --- | --- | --- |
| **Term** | **Short term** | **Overlap** | **Adjusted P-value** | **Genes** |
| Defense Response to Virus (GO:0051607) | Antiviral defense | 12/207 | 3.25E-11 | IFITM3;RTP4;ZBP1;GBP7;ZNXF1;STAT1;STAT2;IRF7;ISG15;SAMHD1;IFIT3;IFIT2 |
| Regulation of Type I Interferon-Mediated Signaling Pathway (GO:0060338) | Type I interferon signaling | 5/38 | 7.59E-06 | ZBP1;STAT2;ISG15;PTPN6;SAMHD1 |
| Regulation of Viral Genome Replication (GO:0045069) | Viral genome replication | 4/66 | 0.001628 | IFITM3;GBP7;ZNXF1;ISG15 |
| Negative Regulation of Viral Process (GO:0048525) | Viral process | 4/67 | 0.001628 | IFITM3;ZNXF1;STAT1;ISG15 |
| Response to Cytokine (GO:0034097) | Cytokine response | 5/143 | 0.001654 | IFITM3;STAT1;ISG15;IFIT3;IFIT2 |
| Response to Type II Interferon (GO:0034341) | Type II interferon response | 4/77 | 0.002117 | IFITM3;GBP7;STAT1;GBP3 |
| Regulation of Innate Immune Response (GO:0045088) | Innate immune response | 4/95 | 0.004286 | TRAFD1;PTPN6;PARP9;SAMHD1 |
| Cellular Response to Cytokine Stimulus (GO:0071345) | Cytokine stimulation | 6/332 | 0.006882 | GBP7;STAT1;PTPN6;IFIT3;IFIT2;GBP3 |
| Type I Interferon-Mediated Signaling Pathway (GO:0060337) | Interferon-mediated signaling | 3/44 | 0.006961 | IFITM3;STAT1;STAT2 |
| Regulation of Cytokine-Mediated Signaling Pathway (GO:0001959) | Cytokine-mediated signaling | 3/48 | 0.007639 | ZBP1;SPPL2A;PTPN6 |
| Cellular Response to Type II Interferon (GO:0071346) | Cellular response to Type II interferon | 3/50 | 0.008011 | GBP7;STAT1;GBP3 |
| Positive Regulation of Cytokine-Mediated Signaling Pathway (GO:0001961) | Regulation of cytokine signaling | 3/53 | 0.00889 | ZBP1;CD40;PARP9 |
| Cell Surface Receptor Signaling Pathway via JAK-STAT (GO:0007259) | JAK-STAT receptor signaling | 3/73 | 0.019006 | CD40;STAT1;STAT2 |
| Positive Regulation of Type I Interferon Production (GO:0032481) | Type I interferon production | 3/73 | 0.019006 | STAT1;IRF7;ISG15 |
| Antiviral Innate Immune Response (GO:0140374) | Antiviral innate immune response | 3/75 | 0.019481 | ZBP1;IFIT3;IFIT2 |

**Supplemental Table 3**

| **Figure S8D. KEGG Pathway Enrichment Analysis of Upregulated Genes in Gene Module 5 Identified by Pseudotime Trajectory Analysis** |  |  |  |  |
| --- | --- | --- | --- | --- |
| **Term** | **Short term** | **Overlap** | **Adjusted P-value** | **Genes** |
| Negative Regulation of Cytokine-Mediated Signaling Pathway (GO:0001960) | Regulation of cytokine signaling | 3/78 | 0.020296 | STAT2;ISG15;SAMHD1 |
| Cell Surface Receptor Signaling Pathway via STAT (GO:0097696) | STAT signaling | 3/80 | 0.020296 | CD40;STAT1;STAT2 |
| Host-mediated Suppression of Symbiont Invasion (GO:0046597) | Host-mediated innate response | 2/20 | 0.021204 | IFITM3;LY6E |
| Positive Regulation of Interferon-Alpha Production (GO:0032727) | Interferon-Alpha production | 2/23 | 0.026932 | STAT1;IRF7 |
| Regulation of Necroptotic Process (GO:0060544) | Necroptosis regulation | 2/25 | 0.030562 | ZBP1;CFLAR |
| Regulation of Angiogenesis (GO:0045765) | Angiogenesis regulation | 4/213 | 0.031451 | CD40;STAT1;KDR;PTPN6 |
| Regulation of Mitochondrial Fission (GO:0090140) | Mitochondrial fission regulation | 2/28 | 0.033148 | STAT2;KDR |
| Regulation of Immune Response (GO:0050776) | Immune response regulation | 3/105 | 0.033148 | SPPL2A;IRF7;SAMHD1 |
| Positive Regulation of Defense Response to Virus by Host (GO:0002230) | Defense response to virus | 2/29 | 0.033148 | STAT1;PARP9 |
| Nuclear Membrane Organization (GO:0071763) | Nuclear membrane organization | 2/29 | 0.033148 | CHMP4B;TOR1AIP1 |
| Defense Response to Bacterium (GO:0042742) | Antibacterial response | 4/236 | 0.037063 | GBP7;ZNF1;ISG15;GBP3 |
| Response to Interferon-Beta (GO:0035456) | Interferon-Beta response | 2/32 | 0.037864 | IFITM3;STAT1 |
| Extrinsic Apoptotic Signaling Pathway via Death Domain Receptors (GO:0008625) | Apoptotic signaling | 2/36 | 0.046388 | DAXX;CFLAR |

**Supplemental Table 3**

| **Figure S8D. KEGG Pathway Enrichment Analysis of Upregulated Genes in Gene Module 6 Identified by Pseudotime Trajectory Analysis** |  |  |  |  |
| --- | --- | --- | --- | --- |
| **Term** | **Short term** | **Overlap** | **Adjusted P-value** | **Genes** |
| Negative Regulation of Intracellular Signal Transduction (GO:1902532) | Inhibition of signal transduction | 12/354 | 1.34E-04 | NFKBIA;IRAK2;PLEK;NLRC5;MDM2;PDE7B;FNIP1;TANK;THBS1;FNIP2;IGF1R;BCL2L1 |
| Cell Surface Toll-Like Receptor Signaling Pathway (GO:0140895) | TLR signaling | 4/31 | 0.007347707 | NFKBIA;IRAK2;PIK3AP1;TLR2 |
| Positive Regulation of Cellular Component Biogenesis (GO:0044089) | Cellular biogenesis | 5/90 | 0.025196906 | SDCBP;SMAD3;PLEK;FNIP1;FNIP2 |
| Regulation of Canonical NF-kappaB Signal Transduction (GO:0043122) | NF-kb signaling | 8/301 | 0.026139902 | NFKBIA;IRAK2;PELI1;NLRC5;LITAF;SQSTM1;TANK;TLR2 |
| Cellular Response to Interleukin-1 (GO:0071347) | IL-1 response | 4/58 | 0.029250884 | NFKBIA;IRAK2;ACOD1;TANK |
| Regulation of TORC1 Signaling (GO:1903432) | TORC1 signaling | 5/109 | 0.029250884 | ATP6V1A;SIK3;PELI1;FNIP1;FNIP2 |
| Toll-like Receptor 2 Signaling Pathway (GO:0034134) | TLR2 signaling | 2/6 | 0.037019895 | PIK3AP1;TLR2 |
| Regulation of Toll-Like Receptor 4 Signaling Pathway (GO:0034143) | TLR4 signaling | 3/29 | 0.037019895 | ZNRF1;ACOD1;SQSTM1 |
| Negative Regulation of Cytokine Production (GO:0001818) | Inhibition of cytokine production | 6/200 | 0.038201051 | SRGN;NFKBIA;CD274;SMAD3;ACOD1;THBS1 |
| Marginal Zone B Cell Differentiation (GO:0002315) | B cell differentiation | 2/7 | 0.038201051 | DOCK10;PTK2B |
| Defense Response to Virus (GO:0051607) | Antiviral defense | 6/207 | 0.038201051 | NLRC5;ACOD1;LYST;TANK;TLR2;BCL2L1 |
| Negative Regulation of Ossification (GO:0030279) | Ossification regulation | 3/33 | 0.038201051 | SRGN;SMAD3;PTK2B |
| Positive Regulation of Intracellular Signal Transduction (GO:1902533) | Activation of intracellular signaling | 11/707 | 0.038201051 | OSBPL8;SMAD3;IRAK2;PELI1;PTK2B;FNIP1;PIK3AP1;LITAF;THBS1;IGF1R;TLR2 |
| Negative Regulation of Canonical NF-kappaB Signal Transduction (GO:0043124) | NF-kb signaling | 4/80 | 0.038201051 | NFKBIA;IRAK2;NLRC5;TANK |
| Cortical Cytoskeleton Organization (GO:0030865) | Cytoskeleton organization | 3/35 | 0.038201051 | FMNL2;PLEK;PTK2B |

**Supplemental Table 3**

| **Figure S8D. KEGG Pathway Enrichment Analysis of Upregulated Genes in Gene Module 6 Identified by Pseudotime Trajectory Analysis** |  |  |  |  |
| --- | --- | --- | --- | --- |
| **Term** | **Short term** | **Overlap** | **Adjusted P-value** | **Genes** |
| Positive Regulation of TOR Signaling<br>(GO:0032008) | TOR signaling | 4/84 | 0.041227661 | ATP6V1A;SIK3;FNIP1;FNIP2 |
| Peptidyl-tyrosine Autophosphorylation<br>(GO:0038083) | Posttranslational modification | 2/10 | 0.054332718 | PTK2B;IGF1R |
| Cellular Response to Tumor Necrosis<br>Factor (GO:0071356) | TNF response | 4/93 | 0.054332718 | NFKBIA;ACOD1;PTK2B;TANK |
